# Exceptions to the rule: Why does resistance evolution not undermine antibiotic therapy in all bacterial infections?

**DOI:** 10.1101/2021.12.15.472803

**Authors:** Amrita Bhattacharya, Anton Aluquin, David A Kennedy

## Abstract

Antibiotic resistance poses one of the greatest public health challenges of the 21^st^ century. Yet not all pathogens are equally affected by resistance evolution. Why? Here we examine what underlies variation in antibiotic resistance across human bacterial pathogens and the drugs used to treat them. We document the observed prevalence of antibiotic resistance for ‘pathogen x drug’ combinations across 57 different human bacterial pathogens and 53 antibiotics from 15 drug classes used to treat them. Using AIC-based model selection we analyze 14 different traits of bacteria and antibiotics that are believed to be important in resistance evolution. Using these data, we identify the traits that best explain observed variation in resistance evolution. Our results show that nosocomial pathogens and indirectly transmitted pathogens are significantly associated with increased prevalence of resistance whereas zoonotic pathogens, specifically those with wild animal reservoirs, are associated with reduced prevalence of resistance. We found partial support for associations between drug resistance and gram classification, human microbiome reservoirs, horizontal gene transfer, and documented human-to human transfer. Global drug use, time since drug discovery, mechanism of drug action, and environmental reservoirs did not emerge as statistically robust predictors of drug resistance in our analyses. To the best of our knowledge this work is the first systematic analysis of resistance across such a wide range of human bacterial pathogens, encompassing the vast majority of common bacterial pathogens. Insights from our study may help guide public health policies and future studies on resistance control.

## Introduction

The emergence and spread of antibiotic resistance poses one of the greatest public heath challenges of the 21^st^ century^1^. Antibiotics impose selection on pathogens to survive, replicate, and transmit in their presence leading to the evolution of antibiotic resistance^2,3^. Yet, drug-resistance evolution does not undermine all antibiotic therapies equally. For example, syphilis, which is caused by the bacterium *Treponema pallidum,* has been treated with penicillin as first-line therapy since 1944, yet a clinical case of penicillin resistance has never been documented^4^. Likewise, antibiotic treatments for Lyme disease, brucellosis, and chlamydia have remained effective despite long histories of drug treatment^4,5^. Here we explore why some ‘pathogen x drug’ combinations are less affected by resistance evolution than others.

Insights about the evolution of drug resistance in clinical and laboratory settings have largely emerged from studies examining pathogens where drug resistance is widespread. For example, efforts to document patterns of drug resistance at national or global scales have typically focused on 13-18 common pathogens where resistance evolution poses the greatest threat to public health^6–8^. Less is known about cases where drug resistance has not yet undermined treatment efficacy. Understanding what drives variation in drug resistance among pathogens can offer important insights to guide antibiotic use in sustainable ways.

Efforts to understand resistance evolution have highlighted many factors that may be important drivers of high levels of resistance^9,10^. These include over-prescription or misuse of antibiotics^10,11^, antibiotic use in farming and agriculture^10,12^, hospital transmission of resistant infections^13^, human-to-human transmission of pathogens^10,14^, gram-classification of bacteria^15,16^, mechanism of drug action^17^, horizontal transfer of antibiotic resistance genes^18–20^, antibiotic release into environmental reservoirs^14,20^, and by-stander selection within the human microbiome^21–23^. Yet, many outstanding questions remain. How well do these factors predict resistance evolution? How important are these factors relative to each other? How might predictors of drug resistance interact and affect the evolution of resistance? Are there strong predictors for low resistance?

Here we investigate these questions by documenting the variation in drug resistance across common human bacterial pathogens and identifying drug and pathogen features that explain the observed variation. We examined 182 unique ‘Pathogen x Drug Class’ combinations, including 57 pathogens and 53 antibiotics within 15 drug classes, derived from a modern medical microbiology textbook^24^. Note that not every drug class is used to treat every pathogen resulting in fewer ‘Pathogen x Drug Class’ combinations than the product of pathogen and drug class numbers. We documented resistance levels using two different approaches that we term the “Expert Opinion Method” (EOM) and “Algorithmic Review Method” (ARM) (Figure 1, Supplemental Tables 1 and 2, see also Methods). Here and throughout the manuscript we use the term ‘resistance levels’ to indicate the qualitative or quantitative prevalence of resistance. In the EOM approach all 182 ‘pathogen x drug class’ combinations were classified into one of three resistance classes: Very Rare/None (0), Rare (1), or Not Rare (2) (Figure 1a). In the ARM approach, the search was expanded to include specific antibiotics within every drug class. The prevalence of resistance (number of isolates resistant out of total number studied) for 376 ‘pathogen x antibiotic’ combinations were acquired from papers published between January 2016 and June 2020. Data were only included from papers that met a strict set of criteria (Figure 1b).

**Figure 1:**
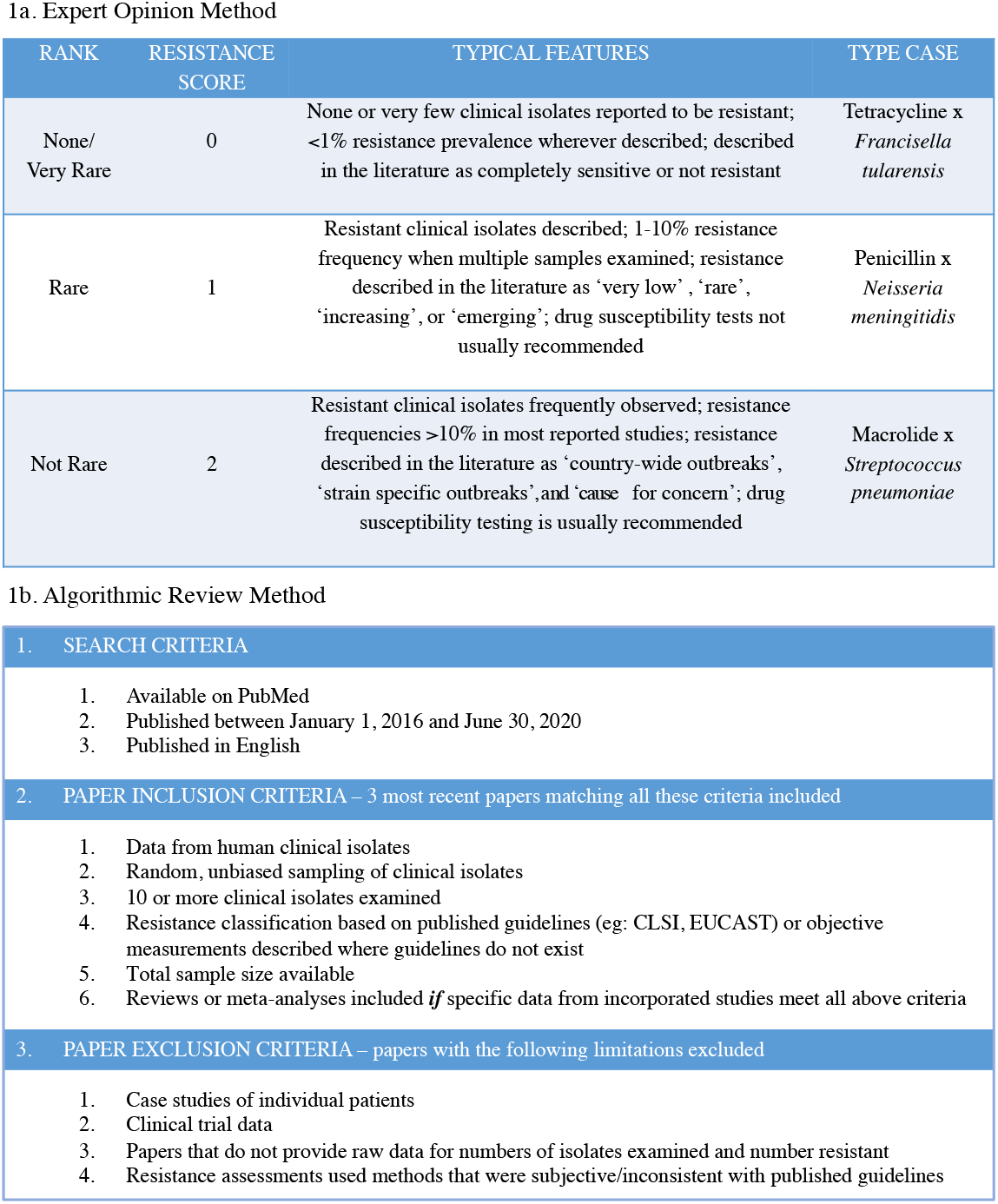
The data collection schemes used for the (a) Expert Opinion Method and (b) Algorithmic Review Method.

Both approaches have strengths and weaknesses. While the ARM data are precise and objective, this method offers lower coverage of the ‘pathogen x drug’ combinations since the necessary data are not available for all combinations of drugs and pathogens (Supplemental Table 2). The EOM method in contrast yields complete coverage of all 182 combinations (Supplemental Table 1) since it allows us to integrate multiple types of data from multiple sources, but it is less precise and is susceptible to researcher bias. We thus use these two complementary approaches. We examined 14 different factors believed to be associated with resistance evolution to identify factors that best explain the variation in resistance (Table 1). To the best of our knowledge this work is the first systematic investigation of resistance evolution to include such a wide range of human bacterial pathogens and predictor variables.

**Table 1.**

| Factor | Levels | Definition |
| --- | --- | --- |
| Nosocomial | Yes/ No | Bacteria that can be acquired from the hospital environment after admission |
| Zoonosis | Yes/No | Bacteria that replicate within animal reservoirs in addition to their infected human hosts. Sustained human to human transmission is independent of this classification |
| Naturally competent | Yes/No | Bacteria contain genetic machinery for horizontal gene transfer via uptake of DNA from the external environment |
| Conjugation | Yes/No | Bacteria contain genetic machinery for horizontal gene transfer between bacterial cells via direct cell-to-cell contact |
| Commensal | Yes/No | Bacteria are opportunistic pathogens can be found within the microbiome of healthy human hosts |
| Transmission type | Direct / Indirect | Mode of transmission. Direct transmission describes bacteria that transfer from reservoir to host via direct, physical contact (e.g.: wound infections caused by contact with contaminated soil, or sexually transmitted infections). Indirect transmission describes bacteria that transfer from reservoir to host via was suspended air particles (airborne), animate intermediaries (vectors), or inanimate objects (vehicles such as canned food, contaminated water) |
| Human transmission | Yes/No | Bacteria where transmission has been documented to occur from one human host to another whether through direct or indirect modes |
| Pathogen type | Gram Positive,<br>Gram Negative,<br>Anaerobic,<br>Other | Bacterial classification based on gram staining or respiration mode as typically discussed in medical microbiology |
| Research effort pathogen | Continuous | Research effort as described by the total number of publications returned for each bacterial pathogen name when searching on the database PubMed |
| Research effort drug/antibiotic | Continuous | Research effort as described by the total number of publications returned for each drug class, or antibiotic name when searching on the database PubMed |
| Drug mechanism | Bactericidal/<br>Bacteriostatic | Broad mechanism of drug action. Bactericidal drugs act by killing pathogens and bacteriostatic drugs inhibit the growth of pathogens without killing them |
| Drug/antibiotic date | Continuous | The year a drug class or antibiotic was first discovered defined by the date of the earliest reference found on Pubmed |
| Global drug use | Continuous | The average consumption of drugs across China, India, and USA (the three countries with highest total drug use). Data are from 2015, the most recent year for which data were publicly available |
| Environmental reservoir | Yes/No | Bacteria that replicate in environmental reservoirs outside the human host. Sustained human to human transmission is independent of this classification |

## Results

### Antibiotic resistance evolution varies widely within and between pathogens and drugs

Using our two approaches, EOM and ARM, we measured resistance across various pathogen x drug combinations (Figure 1, Supplemental Tables 3 and 4). For EOM, resistance was scored 0 (Very Rare/None), 1 (Rare), or 2 (Not Rare) with 0 meaning that resistance was extremely rare, and 2 meaning that resistance was common in at least some geographic locations (Figure 1a). For ARM, resistance prevalence for each ‘pathogen x antibiotic’ combination was calculated from counts of pathogen isolates that were resistant and not resistant in up to three recent studies (see Methods). Resistance was found to vary widely between pathogens (Figure 2A, C) and drugs (Figure 2B, D) across both methods. For the EOM dataset, the overall mean resistance score was 1.05 (median: 1.0) with a standard deviation of 0.84. For the ARM dataset, the overall mean percentage of resistant isolates was 24.29% (median: 9.02%) with a standard deviation of 28.69%.

**Figure 2:**
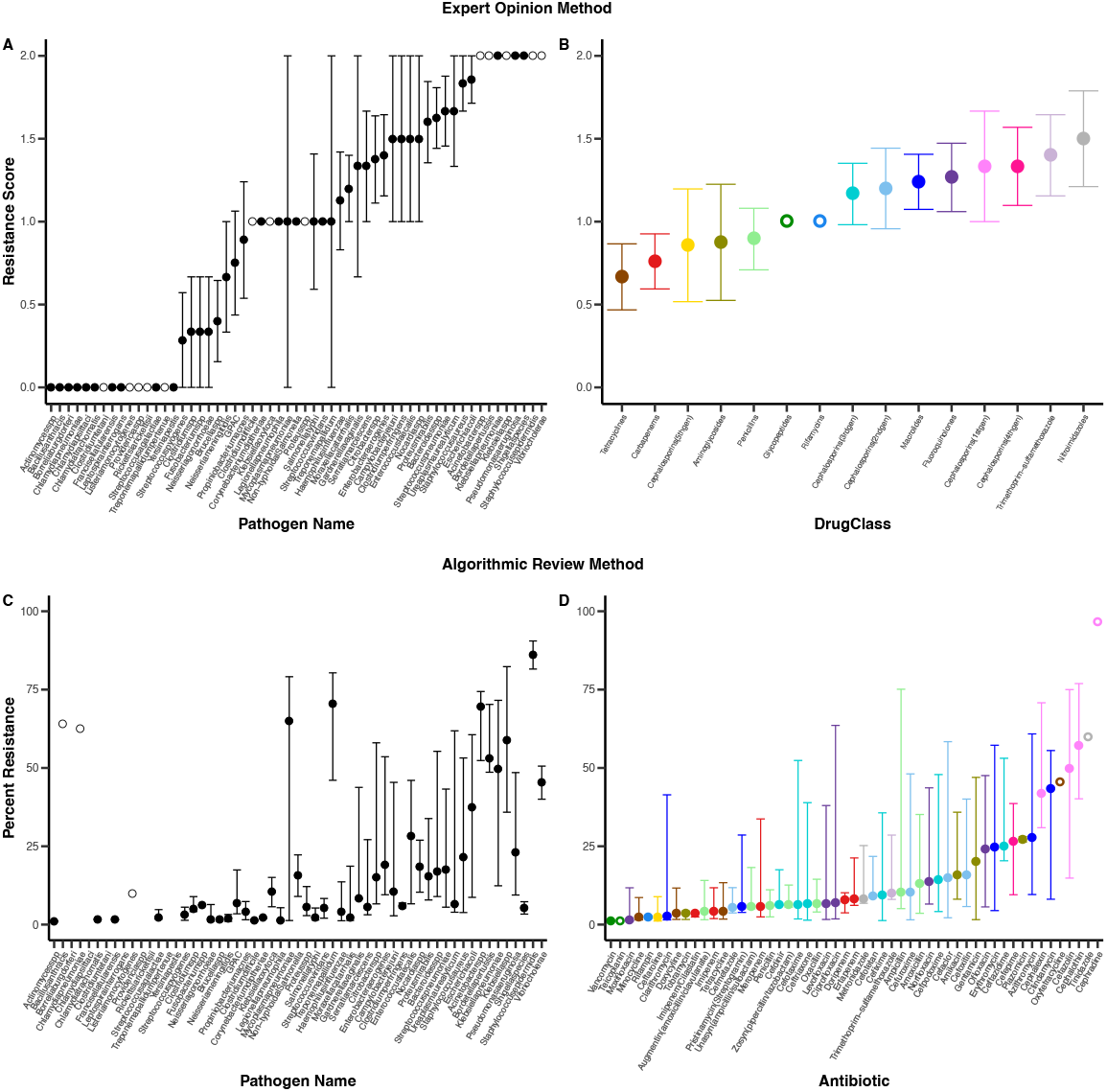
Drug resistance levels vary across pathogens and drugs. Panels A and B show data from the Expert Opinion Method (EOM) and panels C and D show data from the Algorithmic Review Method (ARM). Filled circles in panels A and B depict the mean resistance score +/− 1 standard error across, respectively, pathogens and drug classes in the EOM dataset. Resistance scores 0, 1, and 2 represent the categories Very Rare/None, Rare, and Not Rare as described in Figure 1A. Open circles depict resistance scores based on a single pathogen by drug class combination. Filled circles in panels C and D show the median resistance percentage with error bars depicting the 25^th^ and 75^th^ quantiles for the pathogens and antibiotics in the ARM dataset respectively. Open circles depict resistance scores based on a single datapoint. Note that no ARM data were available for 8 pathogens (missing circles in panel C). These missing datapoints are all pathogens that had low resistance in the EOM dataset.

### Single factor analyses reveal factors that significantly associate with variation in resistance

To test for correlations between our focal factors and resistance levels we performed single factor regressions using mixed effects models in R (linear models for EOM, and logistic models for ARM, see Methods for details). Each regression included a single independent variable as well as, to correct for pseudo replication, random effects of ‘pathogen’, ‘drug class’, and, for the ARM data, ‘antibiotic’. These analyses revealed factors that significantly correlate with variation in resistance for the EOM and ARM datasets (Figure 3 and Figure 4). For the EOM dataset, these analyses revealed ‘nosocomial’, ‘commensal’, ‘conjugation’, ‘natural competence’, ‘zoonosis’, ‘research effort pathogen’, ‘research effort drug’, ‘global drug use’ and ‘drug date’ as significant factors (Figure 3). For the ARM dataset, only ‘nosocomial’ emerged as a statistically significant factor but ‘transmission type’ and ‘human transmission’ showed a trend towards significance (Figure 4). Notably, many of the drug and pathogen factors have significant correlations among them creating opportunities for false positive and false negative trends (Figure 5). The additional analyses described below were performed to account for these correlations and examine the relative importance of different factors.

**Figure 3:**
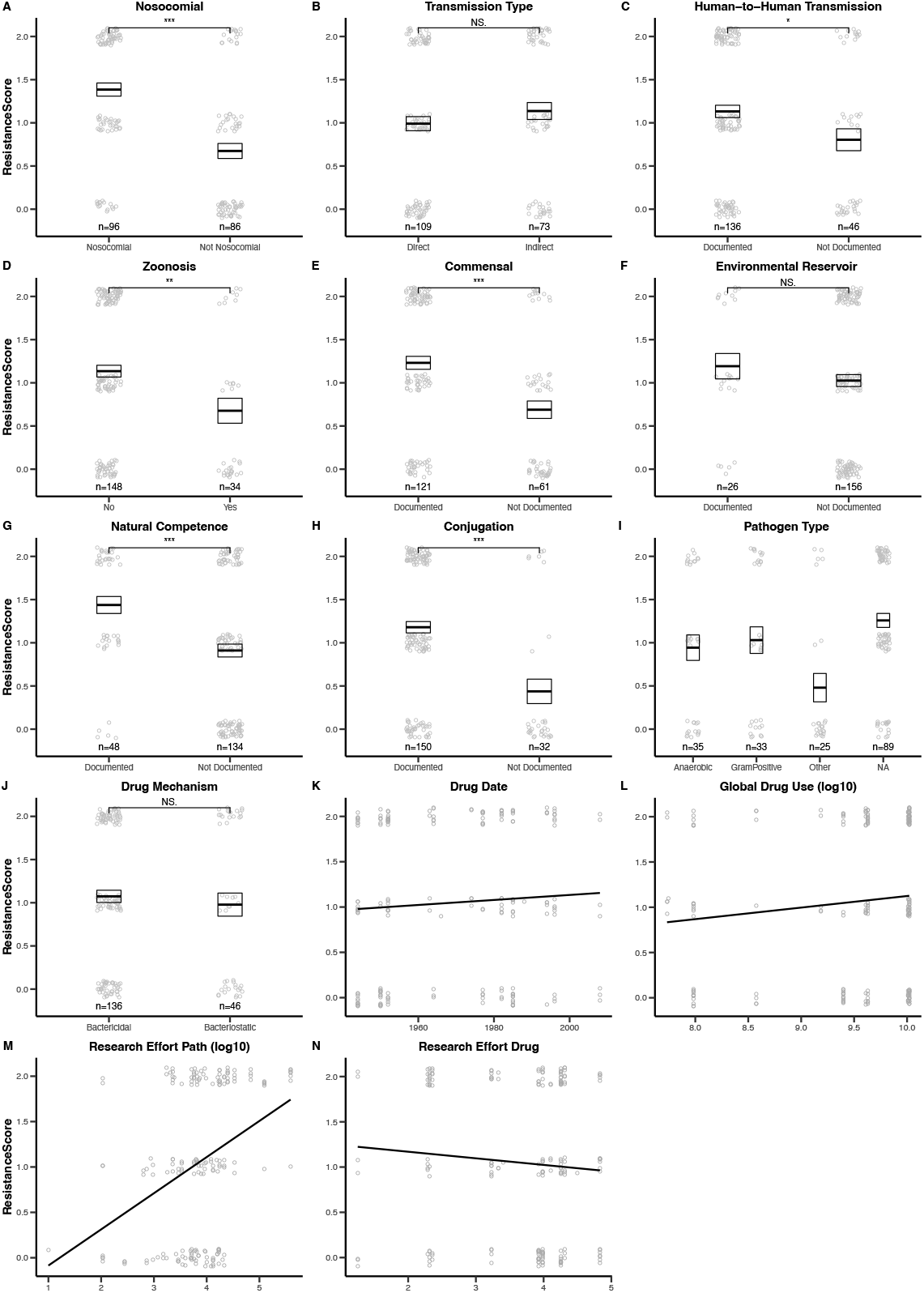
Single factor analyses of drug resistance evolution in EOM dataset. Panels show the correlation between EOM resistance levels and the factors described in Table 1. Grey open circles are the raw data points. Black box plots show the mean +/− 1 standard error. Significance levels represented as * (p < 0.05), ** (p < 0.01), *** (p < 0.001), or # (p <0.1). The presence of a regression line indicates a significant association for continuous factors (panels K-N). Points are jittered to aid visualization. Points in panels K-N are jittered only on the vertical axis.

**Figure 4:**
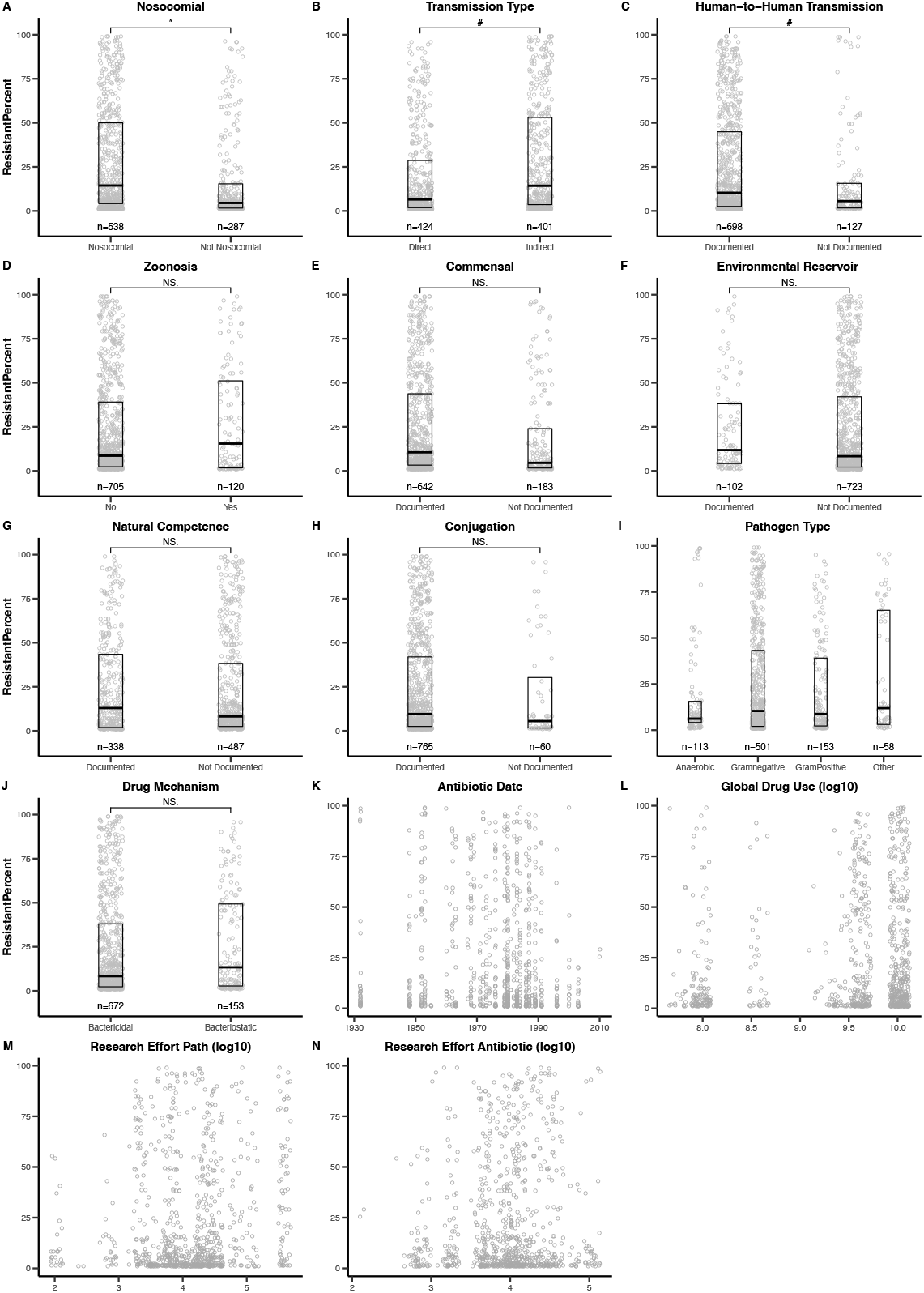
Single factor analyses of drug resistance evolution in ARM dataset. Panels show the correlation between ARM resistance levels and the factors described in Table 1. Grey open circles are the raw data points. Black box plots show the median and first and third quartiles. Significance levels represented as * (p < 0.05), ** (p < 0.01), *** (p < 0.001), or # (p <0.1). For panels N-K, the lack of regression lines indicate no significant associations. Points are jittered on the horizontal axis to aid visualization.

**Figure 5:**
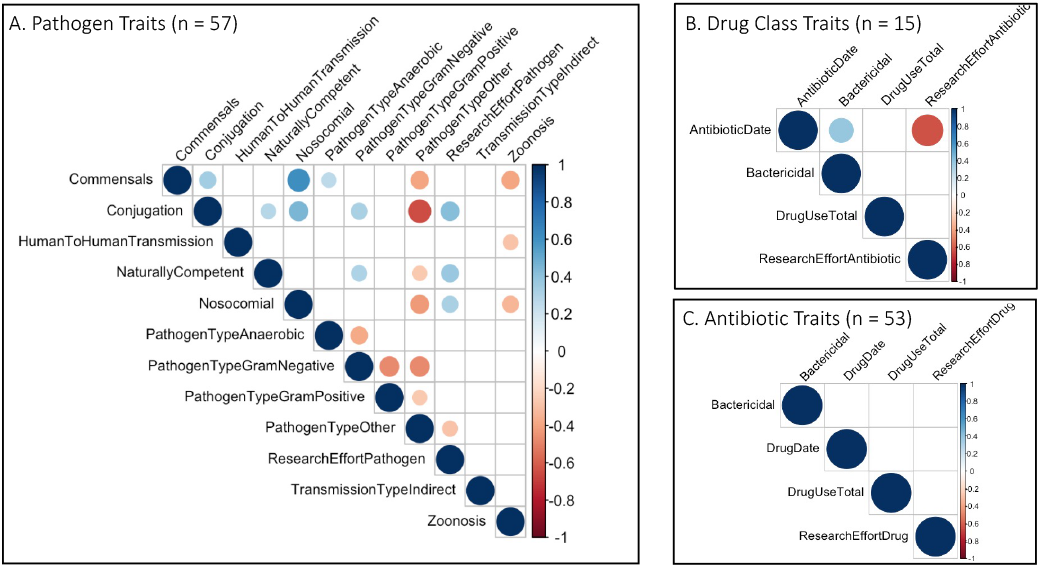
Significant correlations between pathogen traits, drug class factors, and antibiotic factors. Significant correlations (p < 0.05) between different pathogen traits (panel A), between different drug class traits used in EOM dataset (panel B), and between different antibiotic traits used in the ARM dataset (panel C). Color and size of the symbols represent the direction and size of correlations.

### Relative importance of factors in explaining the observed variation in drug resistance

Data from both methods were analyzed using a suite of mixed effects models (linear models for EOM, and logistic models for ARM, see Methods for details). The suite of models consisted of all combinations including or excluding each of our 14 explanatory variables. Each EOM model also included random effects of ‘pathogen’ and ‘drug class’ to correct for pseudo replication, while each ARM model included these random effects in addition to a random effect of ‘antibiotic’. We assessed the relative importance of the 14 factors using two AIC-based metrics: ΔAIC scores and AIC weights of factors (Figure 6, see Methods for details). Considering the two metrics together, ‘nosocomial’ and ‘transmission type’ emerge as important factors across both the EOM and ARM datasets (Figure 6). For the EOM dataset only, ‘zoonosis’, ‘pathogen type’, ‘conjugation’, ‘commensal’, and ‘research effort pathogen’ emerge as important factors. For the ARM dataset only, ‘human transmission’ and ‘pathogen type’ emerge as important factors. ‘Environmental reservoir’, ‘global drug use’, and ‘mechanism of drug action (bactericidal/bacteriostatic)’ do not emerge as important in either of the two datasets.

**Figure 6:**
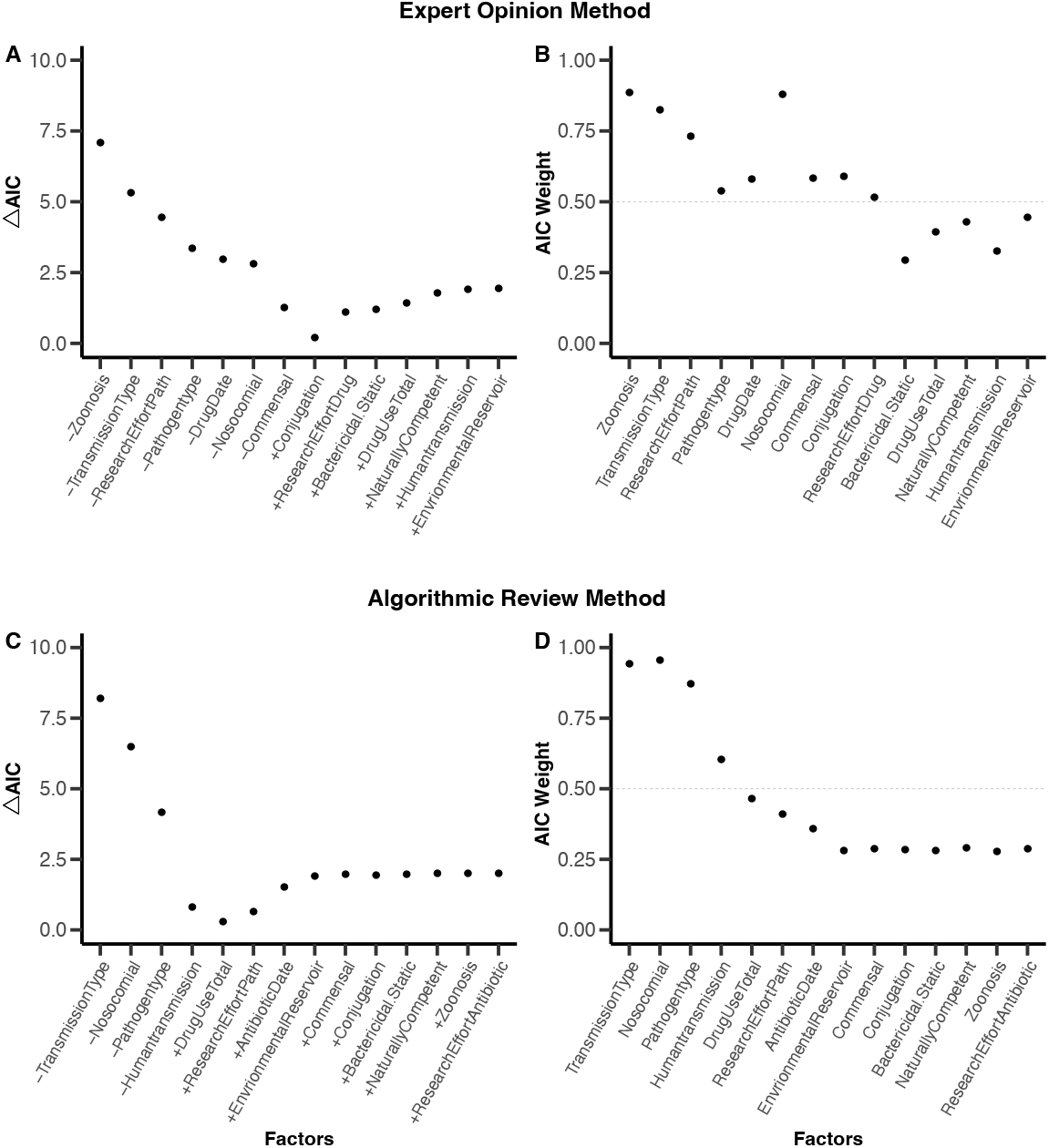
Relative order of importance of factors in explaining variation in drug resistance levels. Panels A and B represent the ΔAIC scores and AIC weights respectively of factors for the Expert Opinion Method dataset. ΔAIC scores represent the difference in AIC scores between the best fitting model with the lowest AIC score and a model with one factor added or removed (denoted respectively by + and – signs on x-axis labels). Factors are arranged in increasing order of importance according to their ΔAIC scores. AIC weights of a factor represent the relative likelihood of all models that include the factor. All factors above the dashed line have AIC weight values greater than 0.5. Factors in panels A and B are arranged in the same order. Panels C and D represent the ΔAIC scores and AIC weights of factors respectively for the ARM dataset. Factors are arranged in increasing order of importance of factors based on their ΔAIC score in Panel C. The same order is maintained in Panel D.

### Relative importance of factors is robust to the removal of individual factors from analyses

To examine the robustness of our results, the AIC weights of all factors were calculated in 14 different scenarios. In each scenario, we eliminated one of the 14 factors, reran the analysis, and recalculated the AIC weights for the 13 remaining factors. Figures 7A and 7B show the AIC weights of factors for all 14 scenarios. For both the EOM and ARM datasets, the relative importance of factors is, in general, robust to the absence of individual factors. In other words, factors that have high AIC weights (>0.5) in the full analysis, generally continue to have high AIC weights (>0.5) when single factors are removed. However, in both EOM and ARM when the factor ‘commensal’ is eliminated from analyses, ‘natural competence’ becomes highly important (AIC weight of factor > 0.8. These results are consistent with the possibility that natural competence is an important factor in explaining resistance evolution but that its contribution is masked when commensal is included as a factor in the analysis. This may be because 8 out of the 12 pathogens in our study that are classified as ‘naturally competent’ are also classified as ‘commensal’.

**Figure 7:**
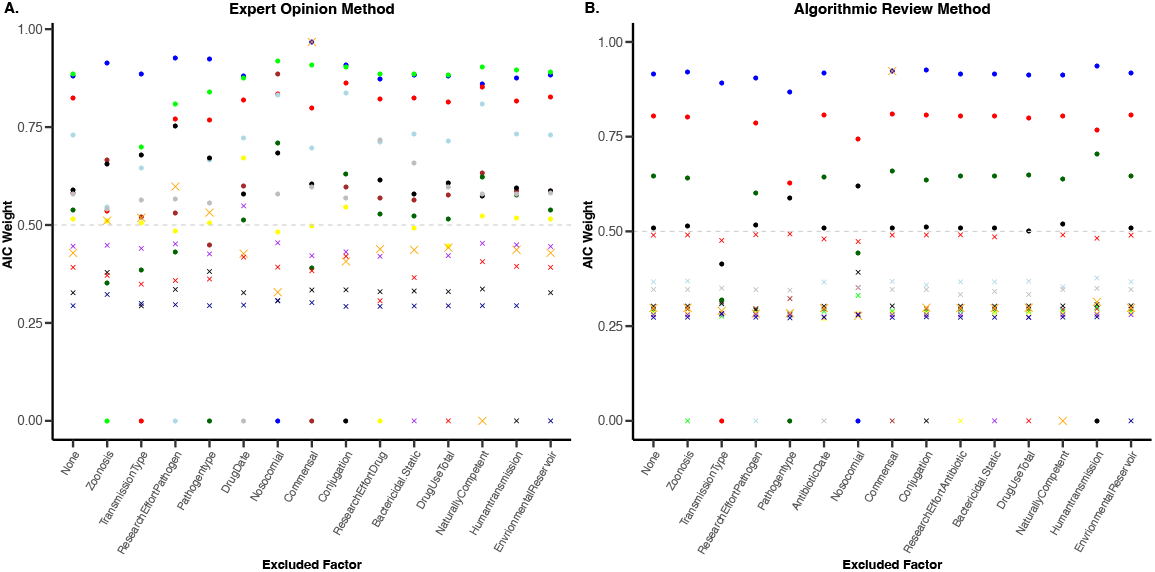
Relative importance of factors is robust to the exclusion of other factors from analysis. Panels A and B represent results from the Expert Opinion Method and Algorithmic Review Method respectively. In both panels the y-axis represents AIC weights with values greater than 0.5 (marked by dashed line). The x-axis lists the factor excluded from each analysis. The unique color-symbol combinations represent each factor. Where no factors were excluded (None) the AIC weights represent results from the full analysis including all 14 factors, also shown in Figure 6. In both EOM and ARM datasets, the AIC weight of ‘natural competence’ increased substantially when ‘commensal’ was removed from the analysis.

### Hospital transmission and indirect transmission modes are associated with high resistance levels while wild-animal zoonotic reservoirs are associated with low resistance levels

Using our full set of models, we calculated AIC-weighted parameter estimates for each of our 14 factors to quantify the correlation between resistance and each factor (Methods). In the EOM analysis, consistent with the AIC weights of factors (Figure 6B), parameter estimates reveal that the factors ‘nosocomial’, ‘transmission type’, ‘research effort pathogen’ and ‘zoonosis’ have significant effects on resistance levels (i.e., 95% confidence intervals do not overlap zero, Figure 8A). While being nosocomial and having indirect transmission are significantly associated with high resistance, being zoonotic is significantly associated with low resistance. Among zoonotic pathogens, those with wild animal reservoirs have substantially lower resistance than those with domesticated animal reservoirs (Supplemental Figure 1). Similarly, in the ARM analysis, the three factors with the highest AIC weights (Figure 6D) namely ‘nosocomial’, ‘transmission type’, and ‘pathogen type’ also have model-averaged parameter estimates that are significantly different than zero. Being nosocomial, having indirect transmission modes, and being grampositive or unclassified by gram-staining (other) are all associated with high levels of resistance (Figure 8B).

**Figure 8:**
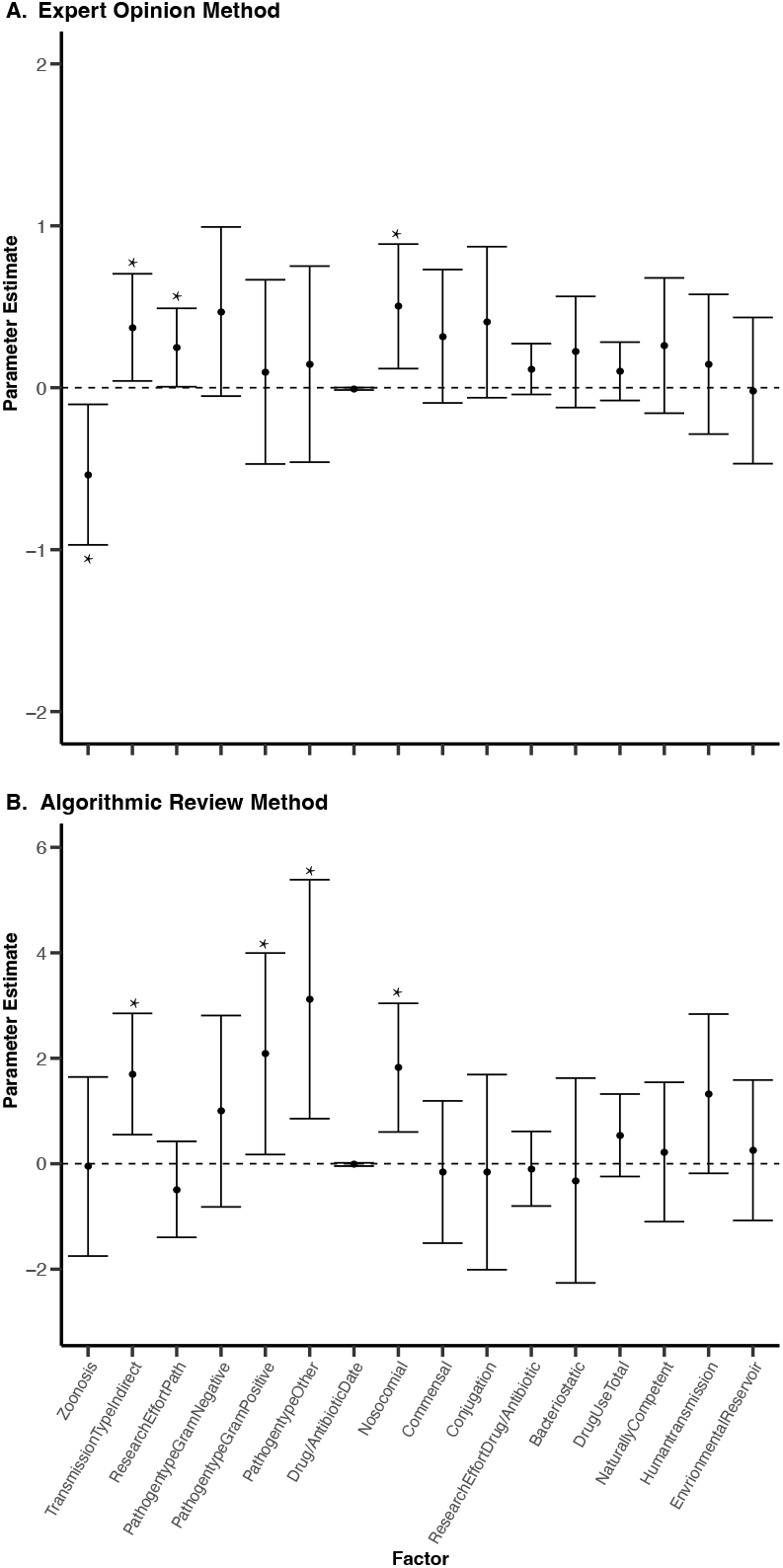
Model averaged parameter estimates of all factors across both EOM and ARM datasets. Model-averaged parameter estimates for each factor were calculated across all models that included the factor. Panels A and B represent results for the Expert Opinion Method and Algorithmic Review Method respectively. Points show model-estimated effect sizes with error bars depicting 95% confidence intervals. Note that for factors with continuous variables, estimated effect sizes are unscaled. Estimates are determined to be significant and marked with an asterisk if the error bars do not overlap ‘0’ (dashed line).

## Discussion

Despite being a huge concern for many human bacterial diseases, the evolution of antibiotic resistance has not undermined treatment efficacy of all ‘pathogen x drug’ combinations. Understanding why may be useful for addressing the ongoing antibiotic resistance crisis. Here, we documented the variation in antibiotic resistance across 57 human bacterial pathogens and 53 antibiotics across 15 drug classes. We explored associations between resistance levels and 14 factors theorized to be important to resistance evolution (Table 1). Our results show that the factors ‘nosocomial’ and ‘indirect transmission’ are associated with increased levels of resistance while ‘zoonosis’, particularly with wild animal reservoirs, is associated with reduced levels of resistance. Factors such as ‘pathogen type’, ‘horizontal gene transfer’, ‘commensalism’, and ‘human-to-human transmission’ may also be important predictors of resistance evolution, but the data for these are less compelling. All results are summarized in Table 2.

**Table 2:**
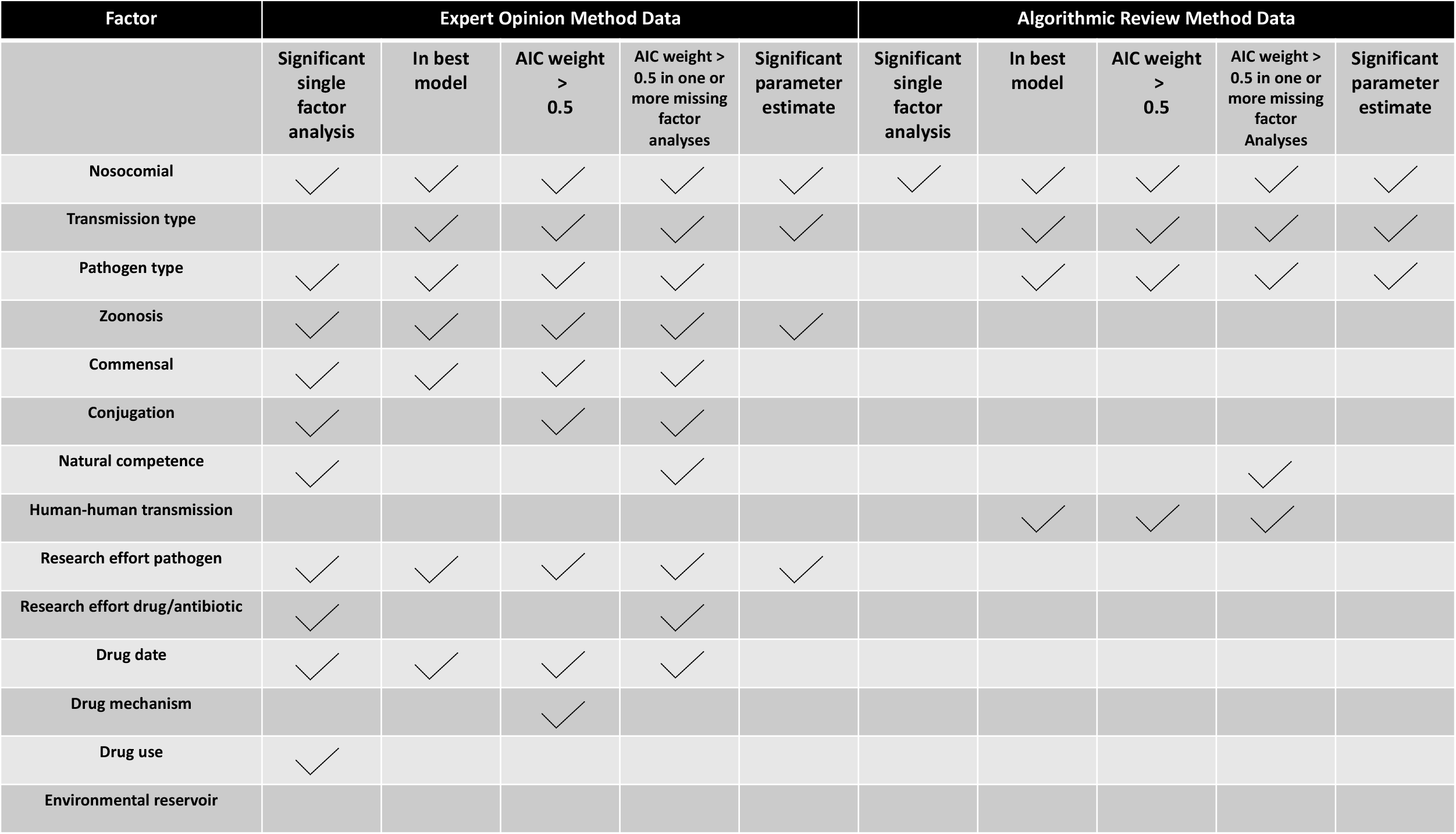
Summary of Results.

According to the WHO, 7% of all hospitalized patients in high-income countries and 10% in low and middle-income countries will acquire at least one nosocomial infection^25^. Many of these infections are caused by pathogens resistant to one or more antibiotics, with some studies reporting proportions as high as 30-50%^26–28^. Consistent with these figures, our results show that nosocomial pathogens are associated with high levels of resistance across both the EOM and ARM datasets (Figure 8). In fact, our results demonstrate that among the various factors implicated in resistance evolution, being nosocomial is one of the most statistically robust predictors (Table 2). The high prevalence of antibiotic resistance among nosocomial pathogens is largely attributed to the interplay of high antibiotic usage, which may impose selection for resistance, and imperfect infection control practices, which may promote transmission of resistant pathogens in hospitals^29–31^. These results highlight the importance of developing protocols to further minimize pathogen transmission within hospitals, for example, by improving sterilization and hygiene practices.

To the best of our knowledge our results are the first to demonstrate a significant relationship between drug resistance and the transmission mode of bacterial pathogens, although the importance of pathogen transmission itself in resistance spread is well appreciated^9,10^. We find that indirectly transmitted pathogens have significantly more resistance than directly transmitted pathogens across both the EOM and ARM datasets (Figures 2, 3, 8). One possible explanation for this correlation is that it results from the difficulty of controlling indirect transmission relative to direct transmission. For instance, the spread of directly transmitted pathogens can be effectively constrained by minimizing contact with an infected host or reservoir. However, indirectly transmitted pathogens may be likely to continue transmitting via air, water, fomites, or vectors even when infection status is already known, and drug therapy is underway. If future research confirms this hypothesis, efforts to better understand the relationship between the mode of transmission and resistance may offer novel approaches towards mitigating the spread of resistance.

Zoonotic pathogens are often assumed to be associated with high rates of antibiotic resistance, largely due to the use of antibiotics in farm animal rearing^10,32–34^. Here we find that zoonotic pathogens are significantly associated with lower levels of resistance in the EOM dataset (Figure 8A). Although this seemingly contradicts expectations, further analysis revealed that the observed pattern in our data comes from pathogens with wild animal reservoirs. Pathogens with wild animal reservoirs exhibit significantly lower levels of resistance than non-zoonotic pathogens, while in contrast, pathogens with domesticated animal reservoirs exhibit levels of resistance that are statistically indistinguishable from non-zoonotic pathogens (Supplemental Figure 1). Limited antibiotic exposure within wild animal reservoirs, and little or no transmission between human hosts (e.g., Lyme disease, anthrax, rickettsia, etc.) constrain the selection and spread of resistance among such zoonotic pathogens resulting in lower levels of resistance. This pattern was not seen in the ARM dataset (Table 2), but since the ARM dataset contained only one pathogen with a wild animal reservoir, similar statistical associations would not be expected.

Our results also show that differences in resistance are associated with gram-status (Table 2) in that anaerobic bacteria had lower levels of resistance than gram-positive and gram-unclassified bacteria in the ARM dataset (Figure 8). There has been a strong focus on drug resistance management for gram-negative bacteria because there are few drugs to treat them^15,16,35^. The impenetrability of the lipopolysaccharide (LPS) layer in gram-negative cell walls is known to render these bacteria inherently resistant to many drug classes^15,16,35^. It has been suggested that the LPS layer in gram-negative cell walls may also heighten acquired resistance mechanisms^36,37^. However, our analyses do not find that gram-negative bacteria have higher acquired resistance levels than gram-positive, anaerobic, or gram-unclassified bacteria. This may indicate that there is no difference in the rate at which resistance arises and spreads between gram-negative pathogens and the remaining bacterial pathogen types, or that efforts to control resistance among gram-negative bacteria have been effective enough to counteract differences in evolutionary potential that might exist. Our results do, however, show that resistance levels are higher in gram-positive and gram-unclassified pathogens than in aerobic bacteria (Figure 8, Table 2), highlighting the need to focus on sustainable approaches to treat these pathogens.

A vast body of literature has implicated horizontal gene transfer in the spread of drug resistance^9,10,18,19,38^. Our results also highlight the importance of the horizontal gene transfer mechanisms. The factor ‘conjugation’ emerged as an important predictor of resistance for the EOM dataset (Table 2), and ‘natural competence’ emerged as an important factor in both the EOM and ARM datasets when ‘commensal’ was removed (Figure 7). We also found that horizontal gene transfer mechanisms positively correlate with being gram-negative and being nosocomial (Figure 5). Horizontal gene transfer has been implicated in the emergence of multidrug resistance in hospital environments and among gram-negative pathogens^18,19,39–41^. Taken together, our results help explain why multidrug resistance is common in these pathogens and environments.

Growing evidence shows that antibiotic resistance is common in the human microbiome, especially in the gut^21,42^, leading to concerns that the microbiome may serve as a reservoir of antibiotic resistance genes^21,43,44^. Our analysis finds that being commensal is a robust predictor of resistance in bacterial pathogens in the EOM dataset (Figure 6A, B). Note that the factor ‘commensal’ as used in this study refers specifically to the subset of commensal bacteria that can be pathogenic, rather than all microbes in the human microbiome (Table 1). Previous work has suggested that inadvertent selection on off-target, commensal bacteria during antibiotic therapy, also known as by-stander selection^23^, may lead to the accumulation of antibiotic resistance genes in the microbiome^23,44,45^. Such accumulation of resistance may not only be concerning for infections resulting from commensal pathogens but also threaten to promote the spread of resistance through horizontal gene transfer to co-infecting, non-commensal pathogens^23,42,44^.

Other factors that emerged as robust predictors in one of the two datasets include drug date and research effort pathogen for the EOM dataset, and human-to-human transmission for the ARM dataset. Although, we did not find other studies that have specifically examined how the discovery date of drugs or person-to-person transmission affect resistance evolution, the associations we find are unsurprising. Our single factor analyses show a significant negative correlation between drug date and resistance score in the EOM dataset meaning that drugs discovered earlier and in use longer have higher levels of resistance than more recently developed drugs. Similarly, our single factors analysis identified a trend of increased resistance among pathogens with human-to-human transmission, which may indicate that such transmission facilitates the spread of resistance once it has emerged.

As far as we are aware, this is the first study to identify pathogen and drug features that correlate with resistance by systematically collecting and analyzing data from a large set of common human bacterial pathogens. Studies to this point have focused on single bacterial pathogens, small sets of bacterial pathogens, or the bacterial pathogens where resistance poses the greatest threats to human health^6,8,38,46–49^, but these approaches neglect comparisons between pathogens where resistance is not a problem and those where it is. Our expansion to such a wide range of pathogens allows for these comparisons, but also poses some inherent challenges. First, few data are available for systems where drug resistance is considered uncommon. Second, many bacterial-drug combinations do not have clear definitions of resistance that can be used to objectively categorize bacterial isolates as resistant or not (e.g., CLSI, EUCAST). Third, resistance is rarely studied in pathogens that are difficult to culture. To overcome these challenges, we devised two complementary approaches for resistance assessment (EOM and ARM, Figure 1), each with distinct strengths and limitations. The EOM approach allowed for the integration of multiple types of data from numerous sources to characterize resistance (Figure 1). This integration is useful in data-limited systems such as for bacteria that are hard to culture, where resistance is poorly defined, and where resistance has never been systematically quantified. However, any systematic biases that exist in the field and appear in literature descriptions of ‘pathogen x drug combinations’ may be reflected in EOM resistance scores. This approach thus offers high coverage of combinations, but it includes inherent subjectivity and coarse estimates of resistance prevalence. Although subjectivity was minimized in the study by using consensus among multiple evaluators, some degree of subjectivity is inevitable (see methods for details). The ARM approach in contrast removed virtually all subjectivity on our part but introduced an alternative form of bias in that data were unavailable for many combinations, particularly those where resistance was difficult to measure or considered to be unimportant to human health. Additionally, the availability of ARM data varies widely across different geographical locations, which can introduce bias when resistance levels vary by location, as they often do^50,51^. The EOM approach reduces this bias by using multiple sources of information to gain a more holistic view of resistance. By using both approaches, and carefully considering the strengths and weaknesses of each, we were able to, for the first time, glean novel insights from a near complete set of common human bacterial pathogens.

The goal of this research was to gain broad insights into patterns of drug resistance evolution and our study thus examined resistance data from all over the globe. While we were able to identify factors that associate with elevated or reduced levels of resistance at such large scales, we note that over shorter timescales and more localized spatial scales key factors may differ. For example, we did not find an effect of total drug use on resistance, but drug use is known to drive resistance increases at finer scales such as within individual hospitals or countries^30,52,53^. We also note that being an observational study, the associations identified here are not necessarily causative. We consider the inherent correlations between the factors examined in the study (Figure 3) and utilize different statistical approaches to identify robust associations with resistance.

We note that there are at least three sets of additional constraints that were unavoidable in our study. First, we have assumed in our analyses that the random effects of each bacterial species and drug class are independent of each other. This may not be correct, if for example, phylogenetic relatedness between bacteria caused closely related bacteria to evolve resistance at similar rates. However, bacterial phylogenies are inherently complex and inconclusive due to horizontal gene transfer between species^54–56^, making it unclear how such relationships could be incorporated into our models. Relationships between drug classes are similarly difficult to incorporate. We did explicitly examine the mechanism of drug action (bactericidal/bacteriostatic, see Table 1) in our analyses but did not find any evidence of a significant association with resistance. Second, we assumed that our categorization of drug and pathogen traits are correct even though some traits were difficult to categorize due to limited data availability. Although we used the most current available classification for each pathogen (Table 1), we note that these classifications may change over the years as new research emerges. For instance, although it is challenging to demonstrate the phenomenon of natural transformation in laboratory conditions, the requisite genetic machinery for competence has been discovered in many species. As a result, the number of human bacterial pathogens considered to be naturally competent has doubled over a span of 20 years^57,58^. Third, the sample size of our study was constrained by virtue of there being a finite number of human bacterial pathogens and drugs used to treat them. It would only be possible to overcome this limitation by expanding to non-human pathogens, but such an expansion would fundamentally change the questions that we could ask^22^.

At the beginning we posed key outstanding questions about drug resistance evolution. These questions include: How well do various factors explain resistance evolution? How important are these factors relative to each other? How might predictors of drug resistance interact and affect the evolution of resistance? Are there strong predictors of low resistance levels? Our results provide some new insights towards addressing these questions. In our single factor analyses, we showed how well each factor alone explains resistance evolution (Figures 2 and 3). In our AIC weight calculations, we showed the relative importance of each factor after accounting for correlations (Figure 6 and 7). In our correlation analysis, we showed the inherent correlations that exist between the pathogen and drug traits examined in this study (Figure 5). Finally, using model-averaged parameters estimates (Figure 8) we identify the factors that predict low levels of drug resistance, such as being zoonotic, directly transmitted, anaerobic and non-nosocomial.

The evolution of drug resistance is among the greatest public health threats of modern times. Here, our goal was to (a) document variation in drug resistance across a wide range of bacterial pathogens and the drugs used to treat them, and (b) identify factors that best explain this variation. Our results confirm the importance of factors like hospital transmission and horizontal gene transfer in promoting resistance spread, highlight areas that may require greater focus (e.g.: acquired resistance in gram-positive pathogens), and motivate future work to explore novel associations (e.g.: the link between pathogen transmission mode and resistance). Taken together, these findings provide novel insights and opportunities to explore interventions that could lead to more sustainable use of available drugs and manage the emergence and spread of drug resistance.

## Methods

### Data collection

Drug resistance for ‘Pathogen x Drug’ combinations was measured using two approaches (a) Expert Opinion Method (EOM) and (b) Algorithmic Review Method (ARM). In the first approach, EOM, all 182 ‘pathogen x drug class’ combinations were classified into one of three resistance levels with each level assigned a numeric score: Very Rare/None (0), Rare (1), or Not Rare (2). These categories were defined as described in Figure 1A. Two evaluators (AB and AA) independently conducted literature surveys for each ‘Pathogen x Drug class’ combination to classify them into one of three levels. A third evaluator (DAK) reviewed all combinations where the calls made by the two evaluators differed (25% of total calls). A final call was made on these combinations by consensus among all three evaluators after discussion (Supplemental Table 3). The second approach, ‘ARM’, used resistance prevalence data (number of resistant and not resistant isolates) collected from papers published between 01/01/2016 and 06/30/2020 available on PubMed using the search term “Pathogen Name and Antibiotic Name Resistance”. Prevalence data for each combination was derived from the three most recent publications that met a comprehensive set of criteria listed in Figure 1b. For ARM, literature searches were expanded to include specific antibiotics within each drug class. Prevalence data for each combination was derived from publications that met a comprehensive set of criteria listed in Figure 1b. Resistance prevalence data was obtained for 376 ‘Pathogen x Antibiotic’ combinations (Supplemental Table 4). These combinations encompassed 169 of the 182 ‘Pathogen x Drug Class’ combinations that were included in the EOM approach, including 49 out of 57 pathogens and all 15 drug classes. For the remaining 13 combinations (including 8 pathogens), data that met the ARM criteria (Figure 1b) were not available.

### Data collection for drug and pathogen characteristics

To identify the factors that best explain the observed variation in resistance, 14 different factors believed to be associated with resistance evolution were investigated (Table 1, Supplemental Table 5). Focal pathogen traits include whether the pathogen is nosocomial (i.e. transmitted in hospital settings), if the pathogen has animal (zoonosis), human microbiome (commensal), or environmental reservoirs, whether the pathogen is gram-positive, gram-negative, anaerobic or other (pathogen type), whether it has direct or indirect transmission modes^59^, and whether human-to-human transmission has been documented. Two modes of horizontal gene transfer, natural competence and conjugation, were also examined. Consistent with currently accepted standards in the field, pathogens were noted to have natural competence or conjugation if evidence was available that a pathogen has the requisite genetic machinery. Drug factors examined in our analyses included how long a drug has been in use (drug date; noted as the date of earliest publication for each drug class name or antibiotic name on PubMed), global drug use (determined by calculating the total drug use from annual per capita drug use data in 2015 for the three countries that report the highest drug usage, i.e. China, India, and USA^60 61^). We also classified each drug as bactericidal (killing) or bacteriostatic (growth-inhibiting). For drugs that can have either mechanism depending on dose, the more common mode was chosen. For example, macrolides, that are primarily bacteriostatic but might act bactericidally in some cases, were classified as bacteriostatic. Finally, differences in research effort devoted to different pathogens and drugs were accounted for by recording the total number of ‘results’ returned on the database PubMed when conducting a general search for each of the 57 pathogen names, 15 drug classes, and 53 antibiotics.

### Statistical Analyses

All statistical analyses were performed on RStudio (Version 1.2.5033). Mixed effects models were used to determine the factors that best explained the variation observed in drug resistance evolution. Models were fit with the ‘lme4’ package^62^ in R Studio using the functions ‘lmer’ for the EOM data and ‘glmer’ with option “family=binomial” for the ARM data.

#### Single Factor Analyses

Linear mixed effects models were used to analyze EOM data, while binomial generalized linear mixed effects models were used for ARM analyses. For the single factor analyses (Figures 3 and 4), EOM data were analyzed using mixed effects models with single factors as fixed effects and ‘Pathogen Name’ and ‘Drug Class’ included in each model as random effects. ARM data were analyzed using binomial generalized linear mixed effects models with single factors again used as fixed effects and ‘Pathogen Name’, ‘Drug class’, and ‘Antibiotic’ as random effects.

#### Best-fitting models

To assess the relative importance of 14 different factors implicated in resistance evolution, each of the EOM and ARM datasets were fit to a suite of mixed-effects models representing all possible combinations of the 14 factors as fixed effects (16,384 total models for each dataset). EOM data were analyzed using linear mixed effects models with ‘Pathogen Name’ and ‘Drug Class’ as random effects. ARM data were analyzed using binomial generalized linear mixed effects models with ‘Pathogen Name’, ‘Drug Class’ as well as ‘Antibiotics’ as random effects. The models with the lowest AIC score for each respective dataset (EOM and ARM) were identified as the best fitting models. The factors included in these best models are listed in Table 2.

#### ΔAIC scores and AIC weights

The relative important of the 14 different factors was assessed using two AIC-based metrics: ΔAIC score and AIC weight. ΔAIC is the difference in AIC scores between the best model and the focal model. The ΔAIC score represents the degree of support for one model over another with smaller values indicating stronger model support^63^. It can also be used to determine the importance of including a particular factor in the best model by either removing a factor that was already in the model or adding a factor that was not part of the model and measuring the change in the AIC score. For factors already included in the best model, a large change in AIC (large ΔAIC) following removal of a factor indicates that the factor is important to explaining the data. For factors not included in the best model, a small change in AIC (small ΔAIC) following the introduction of a factor indicates that the factor partially compensates for the increased model complexity, although not sufficiently to be included in the best model.

The AIC weight of a model is the probability of that model being the “true” model given that the set of all models contains the true model^63^. The AIC weight of a factor is similarly the probability that a particular factor is included in the true model. We calculate AIC weights for each factor by summing the AIC weights for each model that contains the focal factor. In practice, these AIC weights are calculated as

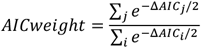

where *j* represents the set of models that include the factor of interest and *i* represents the set of all the models in the analysis. AIC weights greater than 0.5 for a particular factor imply that it is more likely than not to be included in the true model, demonstrating statistical support for the factor. Larger AIC weights demonstrate more support.

#### Missing factors analysis

To test for robustness and potential confounding regarding the importance of individual factors, we again used AIC weights measured in 14 different scenarios. In each scenario one of the 14 factors was eliminated from analysis and the AIC weights of the remaining 13 factors were calculated as described above. Calculated AIC weights should remain nearly constant if the analysis is robust to removing factors. Large changes in AIC weights indicate potential confounding between factors.

#### Model-averaged parameter estimates

Model-averaged parameter estimates for each factor were calculated across all models that included the factor, to estimate effect sizes for each focal factor. In practice, this was performed using the ‘modavg’ package^64^ in R Studio. Factors with 95% confidence intervals not overlapping ‘0’ were considered statistically significant. The mean and 95% CI of parameter estimates for all factors in EOM and ARM are plotted in Figures 8A and 8B respectively.

## Acknowledgements

We thank Zoe Dinges for help with code to design some of the figures in the manuscript. We thank Andrew Read, Beth McGraw, Clara Shaw, and Elsa Hansen for stimulating discussions.

