## Supplemental Materials for "Exceptions to the rule: Why does resistance evolution not undermine antibiotic therapy in all bacterial infections?"

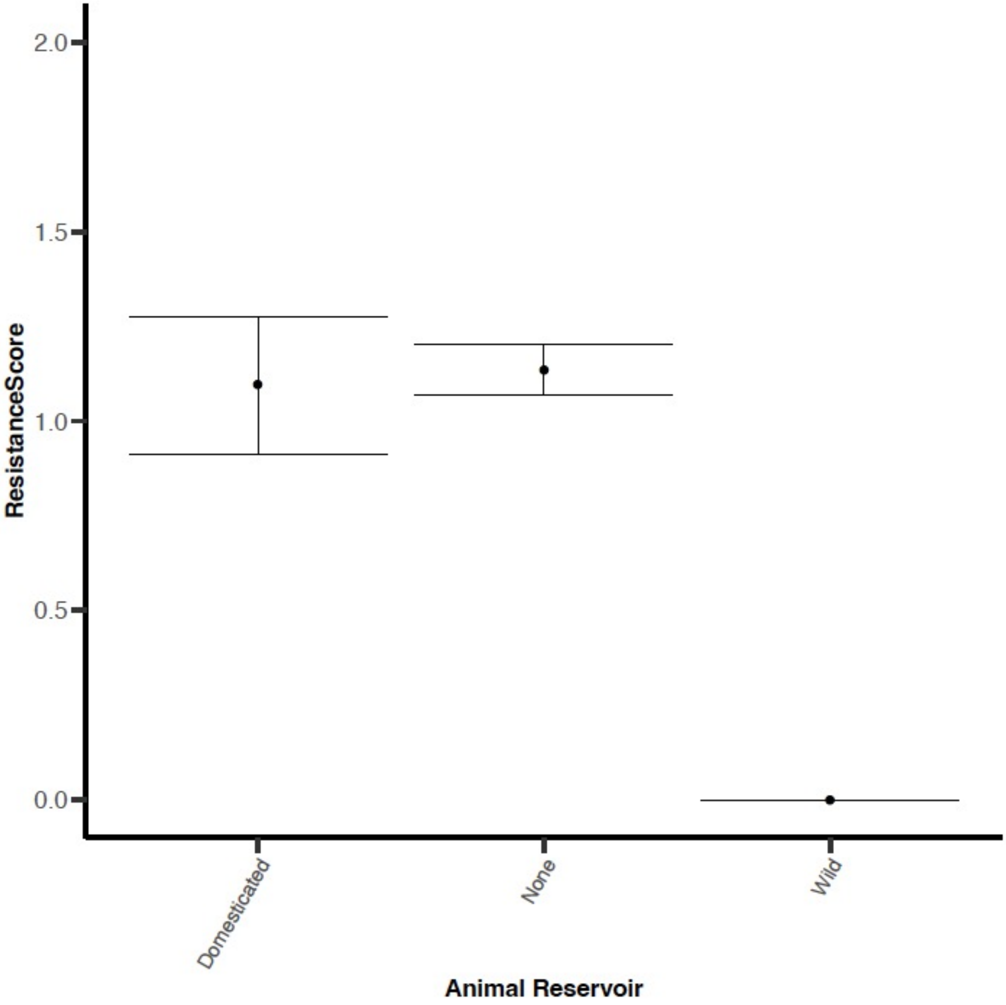

**Supplemental Figure 1:** Zoonotic pathogens with wild animal reservoirs have significantly lower resistance levels than zoonotic pathogens with domesticated reservoirs, and non-zoonotic pathogens. Mean +/- 1 standard error of resistance score data from the Expert Opinion Method data classified by type of animal reservoir. Zoonotic pathogens with wild animal reservoirs have significantly lower resistance scores than those with domesticated animal reservoirs or no animal reservoirs ($F(2,179) = 12.72$, $p < 0.001$).

Supplemental Table 1: Expert Opinion Method Data

| Category | Pathogens | Drug Classes | Pathogen x Drug combinations |
|---|---|---|---|
| Pathogens | 57 | | |
| Drug Classes | | 15 | |
| Combinations | | | 182 |
| Nosocomial | | | |
| Yes | 25 | N/A | 96 |
| No | 32 | N/A | 86 |
| Zoonosis | | | |
| Yes | 12 | N/A | 34 |
| No | 45 | N/A | 148 |
| Commensal | | | |
| Yes | 30 | N/A | 121 |
| No | 27 | N/A | 61 |
| Naturally competent | | | |
| Yes | 12 | N/A | 48 |
| No | 45 | N/A | 134 |
| Conjugation | | | |
| Yes | 44 | N/A | 150 |
| No | 13 | N/A | 32 |
| Human-human transmission | | | |
| Yes | 45 | N/A | 136 |
| No | 12 | N/A | 46 |
| Transmission mode | | | |
| Direct | 33 | N/A | 117 |
| Indirect | 24 | N/A | 65 |

<table>
<tr><th>Category</th><th colspan="2">Pathogens</th><th colspan="2">Drug Classes</th><th colspan="2">Pathogen x Drug combinations</th></tr>
<tr><td colspan="7">Pathogen type</td></tr>
<tr><td>Gram positive</td><td colspan="2">11</td><td colspan="2">N/A</td><td colspan="2">33</td></tr>
<tr><td>Gram negative</td><td colspan="2">26</td><td colspan="2">N/A</td><td colspan="2">89</td></tr>
<tr><td>Anaerobic</td><td colspan="2">8</td><td colspan="2">N/A</td><td colspan="2">35</td></tr>
<tr><td>Other</td><td colspan="2">12</td><td colspan="2">N/A</td><td colspan="2">25</td></tr>
<tr><td colspan="7">Environmental reservoir</td></tr>
<tr><td>Yes</td><td colspan="2">12</td><td colspan="2">N/A</td><td colspan="2">26</td></tr>
<tr><td>No</td><td colspan="2">45</td><td colspan="2">N/A</td><td colspan="2">156</td></tr>
<tr><td colspan="7">Drug mechanism</td></tr>
<tr><td>Bactericidal</td><td colspan="2">N/A</td><td colspan="2">13</td><td colspan="2">136</td></tr>
<tr><td>Bacteriostatic</td><td colspan="2">N/A</td><td colspan="2">2</td><td colspan="2">46</td></tr>
<tr><td colspan="7">Research effort pathogens (log$_{10}$ transformed)</td></tr>
<tr><td>56 unique values</td><td>Mean</td><td>Min.</td><td>1st Qu.</td><td>Median</td><td>3rd Qu.</td><td>Max.</td></tr>
<tr><td></td><td>3.85</td><td>1.0</td><td>3.53</td><td>3.85</td><td>4.24</td><td>5.59</td></tr>
<tr><td colspan="7">Research effort drugs (log$_{10}$ transformed)</td></tr>
<tr><td>15 unique values</td><td>Mean</td><td>Min.</td><td>1st Qu.</td><td>Median</td><td>3rd Qu.</td><td>Max.</td></tr>
<tr><td></td><td>3.65</td><td>1.25</td><td>3.23</td><td>3.98</td><td>4.24</td><td>4.83</td></tr>
<tr><td colspan="7">Drug date</td></tr>
<tr><td>14 unique dates</td><td>Mean</td><td>Min.</td><td>1st Qu.</td><td>Median</td><td>3rd Qu.</td><td>Max.</td></tr>
<tr><td></td><td>1970</td><td>1944</td><td>1950</td><td>1974</td><td>1985</td><td>2008</td></tr>
<tr><td colspan="7">Drug use (log$_{10}$ transformed)</td></tr>
<tr><td>10 unique values<br>(Generation-specific cephalosporin use not available)</td><td>Mean</td><td>Min.</td><td>1st Qu.</td><td>Median</td><td>3rd Qu.</td><td>Max.</td></tr>
<tr><td></td><td>9.42</td><td>7.73</td><td>9.39</td><td>9.63</td><td>10.02</td><td>10.02</td></tr>
</table>

Supplemental Table 2: Algorithmic Review Data

| Category | Pathogens | Antibiotics | Pathogen x Antibiotic combinations |
|---|---|---|---|
| Pathogens | 49 | | |
| Antibiotics | | 53 | |
| Combinations | | | 376 |
| Nosocomial | | | |
| Yes | 25 | N/A | 238 |
| No | 24 | N/A | 138 |
| Zoonosis | | | |
| Yes | 7 | N/A | 52 |
| No | 42 | N/A | 324 |
| Commensal | | | |
| Yes | 29 | N/A | 281 |
| No | 20 | N/A | 95 |
| Naturally competent | | | |
| Yes | 12 | N/A | 140 |
| No | 37 | N/A | 236 |
| Conjugation | | | |
| Yes | 41 | N/A | 343 |
| No | 8 | N/A | 33 |
| Human-human transmission | | | |
| Yes | 40 | N/A | 312 |
| No | 9 | N/A | 64 |
| Transmission mode | | | |
| Direct | 25 | N/A | 181 |
| Indirect | 24 | N/A | 195 |

| Category | Pathogens | Drug Classes | Pathogen x Antibiotic combinations |
|---|---|---|---|
| Pathogen type | | | |
| Gram positive | 11 | N/A | 69 |
| Gram negative | 24 | N/A | 221 |
| Anaerobic | 7 | N/A | 58 |
| Other | 7 | N/A | 28 |
| Environmental reservoir | | | |
| Yes | 10 | N/A | 50 |
| No | 39 | N/A | 326 |
| Drug mechanism | | | |
| Bactericidal | N/A | 44 | 306 |
| Bacteriostatic | N/A | 9 | 70 |

| | | | | | | |
|---|---|---|---|---|---|---|
| Research effort pathogens ($log_{10}$ transformed) | | | | | | |
| 46 unique values | Mean | Min. | 1st Qu. | Median | 3rd Qu. | Max. |
| | 4.11 | 2.03 | 3.76 | 4.12 | 4.53 | 5.59 |
| Research effort antibiotics ($log_{10}$ transformed) | | | | | | |
| 51 unique values | Mean | Min. | 1st Qu. | Median | 3rd Qu. | Max. |
| | 3.94 | 2.18 | 3.68 | 4.00 | 4.25 | 5.04 |
| Antibitoic date | | | | | | |
| 33 unique dates | Mean | Min. | 1st Qu. | Median | 3rd Qu. | Max. |
| | 1975 | 1932 | 1968 | 1980 | 1986 | 2010 |
| Drug use (log transformed) | | | | | | |
| 10 unique values | Mean | Min. | 1st Qu. | Median | 3rd Qu. | Max. |
| (Generation-specific cephalosporin use not available) | 9.49 | 7.73 | 9.39 | 9.63 | 10.02 | 10.02 |

### Supplemental Table 3 : Expert opinion method raw data and citations

| Pathogen Name | Drug Class | Classification | Citations |
|---|---|---|---|
| *Acinetobacter* spp | Carbapenems | **Not Rare** | 1–5 |
| *Actinomyces spp* | Beta-lactams /Penicillins | **None/ Very Rare** | 6–10 |
| *Actinomyces spp* | Carbapenems | **None/ Very Rare** | 9,11 |
| *Actinomyces spp* | Macrolides | **None/ Very Rare** | 9,11 |
| *Bacillus anthracis* | Beta-lactams /Penicillins | **None/ Very Rare** | 12–15 |
| *Bacillus anthracis* | Fluoroquinolones | **None/ Very Rare** | 12,16,17 |
| *Bacillus anthracis* | Tetracyclines | **None/ Very Rare** | 3,12,18 |
| *Bacteroides spp* | Beta-lactams/ Penicillins | **Not Rare** | 19–22 |
| *Bacteroides spp* | Carbapenems | **Rare** | 19,20,23–27 |
| *Bacteroides spp* | Cephalosporins (1st gen) | **Not Rare** | 22,28,29 |
| *Bacteroides spp* | Cephalosporins (2nd gen) | **Rare** | 21,22,27,28,30–32 |
| *Bacteroides spp* | Cephalosporins (3rd gen) | **Not Rare** | 25,28,33,34 |
| *Bacteroides spp* | Macrolides | **Not Rare** | 19,20,28,35,36 |
| *Bacteroides spp* | Nitroimidazoles | **Rare** | 20,28,37,38 |
| *Bacteroides spp* | Tetracyclines | **Not Rare** | 19,20,23,30 |
| *Bordetella pertussis* | Macrolides | **Not Rare** | 39–42 |
| *Borrelia burgdorferi* | Macrolides | **None/ Very Rare** | 43–46 |
| *Borrelia burgdorferi* | Tetracyclines | **None/ Very Rare** | 43,44 |
| *Brucella spp* | Aminoglycosides | **None/ Very Rare** | 47–49 |
| *Brucella spp* | Fluoroquinolones | **None/ Very Rare** | 16,50 |
| *Brucella spp* | Rifampin | **Rare** | 47,49,50 |
| *Brucella spp* | Tetracyclines | **None/ Very Rare** | 16,47,50 |
| *Brucella spp* | Trimethoprim-sulfamethoxazole | **Rare** | 47,50 |
| *Campylobacter jejuni* | Fluoroquinolones | **Not Rare** | 3,51,52 |
| *Campylobacter jejuni* | Macrolides | **Rare** | 51,53,54 |
| *Chlamydia pneumoniae* | Macrolides | **None/ Very Rare** | 55–58 |
| *Chlamydia pneumoniae* | Tetracyclines | **None/ Very Rare** | 58–60 |
| *Chlamydia psittaci* | Macrolides | **None/ Very Rare** | 55,61,62 |

| | | | |
|---|---|---|---|
| *Chlamydia psittaci* | Tetracyclines | **None/ Very Rare** | 62 |
| *Chlamydia trachomatis* | Macrolides | **None / Very Rare** | 55,56,63,64 |
| *Chlamydia trachomatis* | Tetracyclines | **None / Very Rare** | 56,58,59,64 |
| *Citrobacter* spp | Aminoglycosides | **Rare** | 65–68 |
| *Citrobacter* spp | Beta-lactams /Penicillins | **Not Rare** | 65,67,69 |
| *Citrobacter* spp | Carbapenems | **Rare** | 65,67,69–72 |
| *Citrobacter* spp | Cephalosporins (2nd gen) | **Not Rare** | 67,69,73 |
| *Citrobacter* spp | Cephalosporins (3rd gen) | **Not Rare** | 67,69,70,74,75 |
| *Citrobacter* spp | Macrolides | **None/ Very Rare** | 67 |
| *Citrobacter* spp | Tetracyclines | **Not Rare** | 67,76–78 |
| *Citrobacter* spp | Trimethoprim-sulfamethoxazole | **Rare** | 65–67,72,79 |
| *Clostridium difficile* | Vancomycin | **Rare** | 80–83 |
| *Clostridium perfringens* | Beta-lactams /Penicillins | **Rare** | 81,84–88 |
| *Clostridium perfringens* | Tetracyclines | **Not Rare** | 81,84,89,90 |
| *Clostridium spp* | Carbapenems | **None/ Very Rare** | 7,86,91 |
| *Clostridium spp* | Tetracyclines | **None/ Very Rare** | 28,92 |
| *Clostridium spp* | Beta-lactams /Penicillins | **Rare** | 7,86,91–93 |
| *Clostridium tetani* | Tetracyclines | **None/ Very Rare** | 94 |
| *Corynebacterium diptheriae* | Beta-lactams /Penicillins | **Rare** | 95–97 |
| *Corynebacterium diptheriae* | Macrolides | **Rare** | 95–97 |
| *Enterobacter aerogenes* | Carbapenems | **Rare** | 98–100 |
| *Enterobacter aerogenes* | Cephalosporins (2nd gen) | **Not Rare** | 98,101 |
| *Enterobacter aerogenes* | Cephalosporins (3rd gen) | **Not Rare** | 98,101 |
| *Enterobacter aerogenes* | Cephalosporins (4th gen) | **Rare** | 98,100,102,103 |
| *Enterobacter aerogenes* | Fluoroquinolones | **Rare** | 100,101,104–106 |
| *Enterococcus faecalis* | Aminoglycosides | **Not Rare** | 107,108 |
| *Enterococcus faecalis* | Carbapenems | **Rare** | 109–114 |
| *Escherichia coli* | Carbapenems | **Rare** | 115–117 |

| | | | |
|---|---|---|---|
| *Escherichia coli* | Cephalosporins (1st gen) | **Not Rare** | 115 |
| *Escherichia coli* | Cephalosporins (2nd gen) | **Not Rare** | 118,119 |
| *Escherichia coli* | Cephalosporins (3rd gen) | **Not Rare** | 115,118 |
| *Escherichia coli* | Cephalosporins (4th gen) | **Not Rare** | 119–121 |
| *Escherichia coli* | Cephalosporins (5th gen) | **Not Rare** | 122–124 |
| *Escherichia coli* | Fluoroquinolones | **Not Rare** | 125 |
| *Francisella tularensis* | Aminoglycosides | **None/ Very Rare** | 126–128 |
| *Francisella tularensis* | Fluoroquinolones | **None/ Very Rare** | 126,129,130 |
| *Francisella tularensis* | Tetracyclines | **None/ Very Rare** | 128,131 |
| *Fusobacterium spp* | Beta-lactams /Penicillins | **Rare** | 28,92,132–134 |
| *Fusobacterium spp* | Carbapenems | **None/ Very Rare** | 28,92,134,135 |
| *Fusobacterium spp* | Cephalosporins (2nd gen) | **None/ Very Rare** | 7,28,133,134,136–138 |
| *Gardnerella vaginalis* | Carbapenems | **None/ Very Rare** | 139 |
| *Gardnerella vaginalis* | Macrolides | **Not Rare** | 139–141 |
| *Gardnerella vaginalis* | Metronidazole | **Not Rare** | 139–141 |
| *GPAC* | Carbapenems | **None/ Very Rare** | 142–145 |
| *GPAC* | Cephalosporins (1st gen) | **None/ Very Rare** | 142,146 |
| *GPAC* | Cephalosporins (2nd gen) | **None/ Very Rare** | 142,144,145 |
| *GPAC* | Cephalosporins (3rd gen) | **None/ Very Rare** | 142,146 |
| *GPAC* | Macrolides | **Not Rare** | 142–144 |
| *GPAC* | Metronidazole | **Rare** | 142–145,147 |
| *GPAC* | Tetracyclines | **Not Rare** | 142,144,148 |
| *GPAC* | Beta-lactams /Penicillins | **Rare** | 142–145 |
| *Haemophilus influenzae* | Beta-lactams /Penicillins | **Not Rare** | 149,150 |
| *Haemophilus influenzae* | Carbapenems | **None / Very Rare** | 151–153 |
| *Haemophilus influenzae* | Cephalosporins (2nd gen) | **Not Rare** | 151,153–159 |
| *Haemophilus influenzae* | Cephalosporins (3rd gen) | **Rare** | 152,153,156,157,160 |
| *Haemophilus influenzae* | Cephalosporins (5th gen) | **Very Rare/ None** | 122,151,161 |

| | | | |
|---|---|---|---|
| *Haemophilus influenzae* | Fluoroquinolones | **Rare** | 150,151,153,154,162 |
| *Haemophilus influenzae* | Macrolides | **Rare** | 150,152–154,160 |
| *Haemophilus influenzae* | Trimethoprim-sulfamethoxazole | **Not Rare** | 151,153,155,160 |
| *Klebsiella oxytoca* | Cephalosporins (5th gen) | **Rare** | 163–165 |
| *Klebsiella pneumoniae* | Carbapenems | **Not Rare** | 166–170 |
| *Klebsiella pneumoniae* | Cephalosporins (1st gen) | **Not Rare** | 171–173 |
| *Klebsiella pneumoniae* | Cephalosporins (2nd gen) | **Not Rare** | 167,172–174 |
| *Klebsiella pneumoniae* | Cephalosporins (3rd gen) | **Not Rare** | 166,167,175,176 |
| *Klebsiella pneumoniae* | Cephalosporins (4th gen) | **Not Rare** | 166,167,170 |
| *Klebsiella pneumoniae* | Cephalosporins (5th gen) | **Not Rare** | 171,177,178 |
| *Klebsiella pneumoniae* | Fluoroquinolones | **Not Rare** | 170,179,180 |
| *Klebsiella spp* | Aminoglycosides | **Not Rare** | 68,181–183 |
| *Legionella pneumophila* | Fluoroquinolones | **Rare** | 184–186 |
| *Legionella pneumophila* | Macrolides | **Rare** | 184–186 |
| *Leptospira interrogans* | Beta-lactams /Penicillins | **None / Very Rare** | 187,188 |
| *Leptospira interrogans* | Cephalosporins (3rd gen) | **None/ Very Rare** | 187 |
| *Leptospira interrogans* | Tetracyclines | **None/ Very Rare** | 187 |
| *Listeria monocytogenes* | Carbapenems | **None/ Very Rare** | 178,189–191 |
| *Moraxella catarrhalis* | Beta-lactams /Penicillins | **Not Rare** | 192–194 |
| *Moraxella catarrhalis* | Cephalosporins (2nd gen) | **Rare** | 178,189,191,195–198 |
| *Moraxella catarrhalis* | Fluoroquinolones | **Rare** | 189–191,195 |
| *Moraxella catarrhalis* | Macrolides | **Rare** | 178,189,191,196–198 |
| *Moraxella catarrhalis* | Tetracyclines | **Rare** | 178,189–191 |
| *Mycoplasma pneumoniae* | Macrolides | **Not Rare** | 199,200 |
| *Mycoplasma pneumoniae* | Tetracyclines | **Very Rare/ None** | 199,200 |
| *Neisseria gonorrhoeae* | Carbapenems | **Very Rare/ None** | 201 |
| *Neisseria gonorrhoeae* | Cephalosporins (2nd gen) | **Very Rare/ None** | 202,203 |
| *Neisseria gonorrhoeae* | Cephalosporins (3rd gen) | **Rare** | 202–206 |

| | | | |
|---|---|---|---|
| *Neisseria meningitidis* | Beta-lactams /Penicillins | **Rare** | 207–213 |
| *Neisseria meningitidis* | Carbapenems | **None / Very Rare** | 214–216 |
| *Neisseria meningitidis* | Cephalosporins (3rd gen) | **Rare** | 209–212,214,217–219 |
| *Nocardia spp* | Carbapenems | **Not Rare** | 220–225 |
| *Nocardia spp* | Trimethoprim-sulfamethoxazole | **Rare** | 220,222,226,227 |
| *Non-typhoidal Salmonella* | Cephalosporins (3rd gen) | **Rare** | 228,229 |
| *Non-typhoidal Salmonella* | Macrolides | **Rare** | 230–232 |
| *Propiniobacterium acnes* | Beta-lactams /Penicillins | **Very Rare/ None** | 233–236 |
| *Propiniobacterium acnes* | Carbapenems | **Very Rare / None** | 85,134,235,237,238 |
| *Propiniobacterium acnes* | Cephalosporins (1st gen) | **Very Rare / None** | 234,235,239,240 |
| *Propiniobacterium acnes* | Cephalosporins (2nd gen) | **Very Rare / None** | 237,239,241,242 |
| *Propiniobacterium acnes* | Cephalosporins (3rd gen) | **Very Rare / None** | 85,235,237,240 |
| *Propiniobacterium acnes* | Macrolides | **Not Rare** | 243,244 |
| *Propiniobacterium acnes* | Metronidazole | **Not Rare** | 237,239,242,245 |
| *Propiniobacterium acnes* | Tetracyclines | **Not Rare** | 243,246–248 |
| *Propiniobacterium acnes* | Trimethoprim-sulfamethoxazole | **Not Rare** | 247–249 |
| *Proteus mirabilis* | Cephalosporins (1st gen) | **Not Rare** | 250–252 |
| *Proteus mirabilis* | Cephalosporins (2nd gen) | **Not Rare** | 250–253 |
| *Proteus mirabilis* | Cephalosporins (3rd gen) | **Rare** | 250–254 |
| *Proteus mirabilis* | Cephalosporins (4th gen) | **Rare** | 250–252,254,255 |
| *Proteus mirabilis* | Fluoroquinolones | **Not Rare** | 250–253 |
| *Proteus spp* | Carbapenems | **Rare** | 256–259 |
| *Providencia spp* | Carbapenems | **Very Rare / None** | 70,260,261 |
| *Pseudomonas aeruginosa* | Aminoglycosides | **Not Rare** | 262–264 |
| *Pseudomonas aeruginosa* | Carbapenems | **Not Rare** | 262,263,265,266 |
| *Pseudomonas aeruginosa* | Cephalosporins (3rd gen) | **Not Rare** | 4,265,267 |
| *Pseudomonas aeruginosa* | Cephalosporins (4th gen) | **Not Rare** | 262–264 |
| *Rickettsia rickettsii* | Tetracyclines | **Very Rare / None** | 268,269 |

| | | | |
|---|---|---|---|
| *Salmonella typhi* | Carbapenems | **Very Rare / None** | 270,271 |
| *Salmonella typhi* | Cephalosporins (3rd gen) | **Rare** | 270–273 |
| *Salmonella typhi* | Fluoroquinolones | **Not Rare** | 272–274 |
| *Salmonella typhi* | Macrolides | **Rare** | 270–272 |
| *Serratia marcescens* | Carbapenems | **Rare** | 275–277 |
| *Serratia marcescens* | Cephalosporins (3rd gen) | **Not Rare** | 276–279 |
| *Serratia marcescens* | Fluoroquinolones | **Rare** | 277,279 |
| *Shigella* species | Fluoroquinolones | **Not Rare** | 280–285 |
| *Shigella species* | Macrolides | **Not Rare** | 280,281,286 |
| *Staphylococcus aureus* | Carbapenems | **Not Rare** | 287–290 |
| *Staphylococcus aureus* | Cephalosporins (1st gen) | **Not Rare** | 287,288,290 |
| *Staphylococcus aureus* | Cephalosporins (2nd gen) | **Not Rare** | 287,288,291–296 |
| *Staphylococcus aureus* | Cephalosporins (4th gen) | **Not Rare** | 287,288,291,297 |
| *Staphylococcus aureus* | Cephalosporins (5th gen) | **Rare** | 178,288,290,292–296 |
| *Staphylococcus aureus* | Macrolides | **Not Rare** | 298–300 |
| *Staphylococcus epidermidis* | Carbapenems | **Rare** | 301–304 |
| *Staphylococcus epidermidis* | Cephalosporins (1st gen) | **Not Rare** | 302,304 |
| *Streptococcus agalactiae* | Aminoglycosides | **Very Rare / None** | 305–307 |
| *Streptococcus agalactiae* | Cephalosporins (5th gen) | **Very Rare / None** | 165,308,309 |
| *Streptococcus pneumoniae* | Beta-lactams /Penicillins | **Not Rare** | 310–312 |
| *Streptococcus pneumoniae* | Carbapenems | **Not Rare** | 287,310,313,314 |
| *Streptococcus pneumoniae* | Cephalosporins (2nd gen) | **Not Rare** | 312,313,315,316 |
| *Streptococcus pneumoniae* | Cephalosporins (3rd gen) | **Rare** | 151,310,317 |
| *Streptococcus pneumoniae* | Cephalosporins (4th gen) | **Rare** | 313,318–320 |
| *Streptococcus pneumoniae* | Macrolides | **Not Rare** | 310,321–325 |
| *Streptococcus pyogenes* | Beta-lactams /Penicillins | **None / Very Rare** | 309,326,327 |
| *Streptococcus pyogenes* | Cephalosporins (1st gen) | **None / Very Rare** | 328,329 |
| *Streptococcus pyogenes* | Cephalosporins (2nd gen) | **None / Very Rare** | 295,315,330,331 |

| | | | |
|---|---|---|---|
| *Streptococcus pyogenes* | Cephalosporins (3rd gen) | **None / Very Rare** | 309,326,331,332 |
| *Streptococcus pyogenes* | Cephalosporins (4th gen) | **None / Very Rare** | 333,334 |
| *Streptococcus pyogenes* | Cephalosporins (5th gen) | **None / Very Rare** | 165,308,309 |
| *Streptococcus pyogenes* | Macrolides | **Not Rare** | 326–328,333 |
| *Streptococcus viridans* | Beta-lactams /Penicillins | **Rare** | 300,335–340 |
| *Streptococcus viridans* | Cephalosporins (4th gen) | **Rare** | 341–343 |
| *Treponema pallidum* | Beta-lactams /Penicillins | **None / Very Rare** | 344–346 |
| *Treponema pallidum* | Macrolides | **Not Rare** | 344,347,348 |
| *Treponema pallidum pertenue* | Beta-lactams /Penicillins | **None / Very Rare** | 349,350 |
| *Ureaplasma urealyticum* | Fluoroquinolones | **Not Rare** | 351–353 |
| *Ureaplasma urealyticum* | Macrolides | **Not Rare** | 351–354 |
| *Ureaplasma urealyticum* | Tetracyclines | **Rare** | 351–356 |
| *Vibrio cholerae* | Tetracyclines | **Not Rare** | 357–359 |
| *Yersinia pestis* | Aminoglycosides | **None / Very Rare** | 360,361 |
| *Yersinia pestis* | Tetracyclines | **None / Very Rare** | 16,360,361 |

7. Marchand-Austin, A. *et al.* Antimicrobial susceptibility of clinical isolates of anaerobic bacteria in

Ontario, 2010-2011. *Anaerobe* **28**, 120–125 (2014).

24. Cobo, F. *et al.* Clinical findings and antimicrobial susceptibility of anaerobic bacteria isolated in

bloodstream infections. *Antibiotics* **9**, 1–9 (2020).

Genes and Macrolides Resistance. *Pediatr. Infect. Dis. J.* **38**, 370–376 (2019).

71. Lalaoui, R. *et al.* Genomic characterization of Citrobacter freundii strains coproducing OXA-48 and VIM-1 carbapenemase enzymes isolated in leukemic patient in Spain. doi:10.1186/s13756-019-

0630-3.

91. Pathogen Safety Data Sheets: Infectious Substances – Clostridium spp. *Canada Department of Health* https://www.canada.ca/en/public-health/services/laboratory-biosafety-biosecurity/pathogen-safety-data-sheets-risk-assessment/clostridium.html.

102. Thiolas, A., Bornet, C., Davin-Régli, A., Pagès, J. M. & Bollet, C. Resistance to imipenem, cefepime, and cefpirome associated with mutation in Omp36 osmoporin of Enterobacter aerogenes.

*Biochem. Biophys. Res. Commun.* **317**, 851–856 (2004).

103. Eugene Sanders, W. E. & Sanders, C. C. Enterobacter spp.: Pathogens poised to flourish at the turn of the century. *Clin. Microbiol. Rev.* **10**, 220–241 (1997).

104. Mishra, M. P., Sarangi, R. & Padhy, R. N. Prevalence of multidrug resistant uropathogenic bacteria in pediatric patients of a tertiary care hospital in eastern India. *J. Infect. Public Health* **9**, 308–314 (2016).

105. G, J.-G. *et al.* Susceptibility evolution to antibiotics of Enterobacter cloacae, Morganella morganii, Klebsiella aerogenes and Citrobacter freundii involved in urinary tract infections: an 11-year epidemiological surveillance study. *Enfermedades Infecc. y Microbiol. Clin. (English ed.)* **38**, 166–169 (2020).

106. Akhtar, N., Alqurashi, A. & Twibah, M. In vitro ciprofloxacin resistance profiles among gram-negative bacteria isolated from clinical specimens in a teaching hospital. *J. Pakistan Med. Assoc.* (2010).

146. Kuriyama, T., Karasawa, T., Nakagawa, K., Yamamoto, E. & Nakamura, S. Bacteriology and antimicrobial susceptibility of gram-positive cocci isolated from pus specimens of orofacial

odontogenic infections. *Oral Microbiol. Immunol.* **17**, 132–135 (2002).

161. Sader, H. S., Flamm, R. K., Streit, J. M., Carvalhaes, C. G. & Mendes, R. E. Antimicrobial activity of ceftaroline and comparator agents tested against organisms isolated from patients with

community-acquired bacterial pneumonia in Europe, Asia, and Latin America. *Int. J. Infect. Dis.* **77**, 82–86 (2018).

171. Bui, T. & Preuss, C. V. *Cephalosporins*. *NCBI Bookshelf* (StatPearls Publishing LLC, 2021).

232. DRUG-RESISTANT NONTYPHOIDAL SALMONELLA.

238. Shames, R., Satti, F., Vellozzi, E. M. & Smith, M. A. Susceptibilities of Propionibacterium acnes

ophthalmic isolates to ertapenem, meropenem, cefepime. *J. Clin. Microbiol.* **44**, 4227–4228 (2006).

268. Government of Canada. Pathogen Safety Data Sheet: Rickettsia rickettsii.

300. Kim, D. *et al.* Increasing resistance to extended-spectrum cephalosporins, fluoroquinolone, and

carbapenem in gram-negative bacilli and the emergence of carbapenem non-susceptibility in klebsiella pneumoniae: Analysis of Korean Antimicrobial Resistance Monitoring System . *Ann. Lab. Med.* **37**, 231–239 (2017).

315. Zafar, A. *et al.* Antibiotic susceptibility in Streptococcus pneumoniae, Haemophilus influenzae and Streptococcus pyogenes in Pakistan: A review of results from the survey of antibiotic

resistance (SOAR) 2002-15. *J. Antimicrob. Chemother.* **71**, i103–i109 (2016).

*Antimicrob. Chemother.* **71**, i71–i83 (2016).

#### Supplemental Table 4: Algorthmic review method raw data and citations

| Pathogen Name | Drug Class | Antibiotic | Resistant | Not Resistant | Total Sample Size | Citation |
|---|---|---|---|---|---|---|
| Acinetobacterspp | Carbapenems | Doripenem | 49 | 101 | 150 | 1 |
| Acinetobacterspp | Carbapenems | Imipenem | 23 | 18 | 41 | 2 |
| Acinetobacterspp | Carbapenems | Imipenem | 33 | 5 | 38 | 3 |
| Acinetobacterspp | Carbapenems | Imipenem | 1495 | 643 | 2138 | 4 |
| Acinetobacterspp | Carbapenems | Meropenem | 188 | 47 | 235 | 5 |
| Acinetobacterspp | Carbapenems | Meropenem | 105 | 45 | 150 | 1 |
| Acinetobacterspp | Carbapenems | Meropenem | 236 | 247 | 483 | 6 |
| Actinomycesspp | Penicillins | Penicillin | 0 | 89 | 89 | 7 |
| Actinomycesspp | Penicillins | Unasyn(ampicillin/sulbactam) | 0 | 360 | 360 | 7 |
| Bacillusanthracis | Penicillins | Ampicillin | 89 | 49 | 138 | 8 |
| Bacteroidesspp | Penicillins | Penicillin | 61 | 1 | 62 | 9 |
| Bacteroidesspp | Penicillins | Penicillin | 157 | 0 | 157 | 10 |
| Bacteroidesspp | Penicillins | Penicillin | 77 | 5 | 82 | 11 |
| Bacteroidesspp | Penicillins | Amoxicillin | 96 | 5 | 101 | 12 |
| Bacteroidesspp | Penicillins | Ampicillin | 157 | 0 | 157 | 10 |
| Bacteroidesspp | Penicillins | Ampicillin | 77 | 2 | 79 | 13 |
| Bacteroidesspp | Penicillins | Ampicillin | 20 | 0 | 20 | 14 |

| | | | | | | |
|---|---|---|---|---|---|---|
| Bacteroidesspp | Penicillins | Augmentin(amoxicillin/clavulanate) | 20 | 42 | 62 | 9 |
| Bacteroidesspp | Penicillins | Augmentin(amoxicillin/clavulanate) | 27 | 180 | 207 | 10 |
| Bacteroidesspp | Penicillins | Augmentin(amoxicillin/clavulanate) | 16 | 66 | 82 | 11 |
| Bacteroidesspp | Penicillins | Unasyn(ampicillin/sulbactam) | 6 | 73 | 79 | 13 |
| Bacteroidesspp | Penicillins | Unasyn(ampicillin/sulbactam) | 3 | 17 | 20 | 14 |
| Bacteroidesspp | Penicillins | Zosyn(pipercillin/tazobactam) | 9 | 53 | 62 | 9 |
| Bacteroidesspp | Penicillins | Zosyn(pipercillin/tazobactam) | 28 | 179 | 207 | 10 |
| Bacteroidesspp | Penicillins | Zosyn(pipercillin/tazobactam) | 3 | 76 | 79 | 13 |
| Bacteroidesspp | Carbapenems | Doripenem | 7 | 72 | 79 | 13 |
| Bacteroidesspp | Carbapenems | Imipenem | 2 | 60 | 62 | 9 |
| Bacteroidesspp | Carbapenems | Imipenem | 16 | 99 | 115 | 15 |
| Bacteroidesspp | Carbapenems | Imipenem | 2 | 205 | 207 | 10 |
| Bacteroidesspp | Carbapenems | Meropenem | 16 | 99 | 115 | 15 |
| Bacteroidesspp | Carbapenems | Meropenem | 2 | 205 | 207 | 10 |
| Bacteroidesspp | Carbapenems | Meropenem | 2 | 80 | 82 | 11 |
| Bacteroidesspp | Carbapenems | Ertapenem | 27 | 111 | 138 | 16 |
| Bacteroidesspp | Cephalosporins(2ndgen) | Cefmetazole | 44 | 35 | 79 | 13 |
| Bacteroidesspp | Cephalosporins(2ndgen) | Cefmetazole | 2 | 18 | 20 | 14 |
| Bacteroidesspp | Cephalosporins(2ndgen) | Cefoxitin | 12 | 103 | 115 | 15 |

| | | | | | | |
|---|---|---|---|---|---|---|
| Bacteroidesspp | Cephalosporins(2ndgen ) | Cefoxitin | 73 | 134 | 207 | 10 |
| Bacteroidesspp | Cephalosporins(3rdgen ) | Ceftriaxone | 11 | 9 | 20 | 14 |
| Bacteroidesspp | Macrolides | Clindamycin | 28 | 34 | 62 | 9 |
| Bacteroidesspp | Macrolides | Clindamycin | 92 | 23 | 115 | 15 |
| Bacteroidesspp | Macrolides | Clindamycin | 110 | 97 | 207 | 10 |
| Bacteroidesspp | Nitroimidazoles | Metronidazole | 2 | 60 | 62 | 9 |
| Bacteroidesspp | Nitroimidazoles | Metronidazole | 6 | 109 | 115 | 15 |
| Bacteroidesspp | Nitroimidazoles | Metronidazole | 12 | 195 | 207 | 10 |
| Bordetellapertussis | Macrolides | Erythromycin | 293 | 42 | 335 | 17 |
| Bordetellapertussis | Macrolides | Erythromycin | 51 | 54 | 105 | 18 |
| Bordetellapertussis | Macrolides | Erythromycin | 81 | 60 | 141 | 19 |
| Bordetellapertussis | Macrolides | Clarithromycin | 51 | 54 | 105 | 18 |
| Bordetellapertussis | Macrolides | Clarithromycin | 0 | 135 | 135 | 20 |
| Bordetellapertussis | Macrolides | Azithromycin | 51 | 54 | 105 | 18 |
| Bordetellapertussis | Macrolides | Azithromycin | 81 | 60 | 141 | 19 |
| Bordetellapertussis | Macrolides | Azithromycin | 95 | 31 | 126 | 21 |
| Bordetellapertussis | Macrolides | Clindamycin | 51 | 54 | 105 | 18 |
| Bordetellapertussis | Macrolides | Clindamycin | 95 | 31 | 126 | 21 |
| Brucellaspp | Aminoglycosides | Gentamicin | 0 | 85 | 85 | 22 |
| Brucellaspp | Aminoglycosides | Gentamicin | 0 | 57 | 57 | 23 |

| | | | | | | |
|---|---|---|---|---|---|---|
| Brucellaspp | Fluoroquinolones | Ciprofloxacin | 0 | 85 | 85 | 22 |
| Brucellaspp | Fluoroquinolones | Ciprofloxacin | 0 | 57 | 57 | 23 |
| Brucellaspp | Fluoroquinolones | Levofloxacin | 0 | 85 | 85 | 22 |
| Brucellaspp | Fluoroquinolones | Moxifloxacin | 0 | 57 | 57 | 23 |
| Brucellaspp | Rifamycins | Rifampin | 1 | 59 | 60 | 24 |
| Brucellaspp | Rifamycins | Rifampin | 1 | 84 | 85 | 22 |
| Brucellaspp | Rifamycins | Rifampin | 0 | 57 | 57 | 23 |
| Brucellaspp | Tetracyclines | Doxycyline | 0 | 85 | 85 | 22 |
| Brucellaspp | Tetracyclines | Doxycyline | 0 | 57 | 57 | 23 |
| Brucellaspp | Tetracyclines | Minocycline | 0 | 85 | 85 | 22 |
| Brucellaspp | Tetracyclines | Tetracycline | 0 | 85 | 85 | 22 |
| Brucellaspp | Trimethoprim-sulfamethoxazole | Trimethoprim-sulfamethoxazole | 6 | 79 | 85 | 22 |
| Brucellaspp | Trimethoprim-sulfamethoxazole | Trimethoprim-sulfamethoxazole | 0 | 57 | 57 | 23 |
| Campylobacterjejuni | Fluoroquinolones | Ciprofloxacin | 11 | 0 | 11 | 25 |
| Campylobacterjejuni | Fluoroquinolones | Ciprofloxacin | 111 | 88 | 199 | 26 |
| Campylobacterjejuni | Fluoroquinolones | Ciprofloxacin | 114 | 34 | 148 | 27 |
| Campylobacterjejuni | Fluoroquinolones | Norfloxacin | 1 | 10 | 11 | 28 |
| Campylobacterjejuni | Macrolides | Erythromycin | 4 | 195 | 199 | 26 |
| Campylobacterjejuni | Macrolides | Erythromycin | 4 | 1196 | 1200 | 29 |
| Campylobacterjejuni | Macrolides | Erythromycin | 0 | 97 | 97 | 30 |

| | | | | | | |
|---|---|---|---|---|---|---|
| Campylobacterjejuni | Macrolides | Azithromycin | 1 | 10 | 11 | 25 |
| Campylobacterjejuni | Macrolides | Azithromycin | 1 | 104 | 105 | 31 |
| Campylobacterjejuni | Macrolides | Azithromycin | 29 | 567 | 596 | 32 |
| Chlamydiapneumoniae | Macrolides | Erythromycin | 44 | 26 | 70 | 33 |
| Chlamydiatrachomatis | Macrolides | Erythromycin | 0 | 58 | 58 | 34 |
| Chlamydiatrachomatis | Macrolides | Clarithromycin | 0 | 58 | 58 | 34 |
| Chlamydiatrachomatis | Tetracyclines | Doxycyline | 0 | 58 | 58 | 34 |
| Chlamydiatrachomatis | Tetracyclines | Minocycline | 0 | 58 | 58 | 34 |
| Citrobacterspp | Aminoglycosides | Gentamicin | 6 | 379 | 385 | 35 |
| Citrobacterspp | Aminoglycosides | Amikacin | 53 | 3724 | 3777 | 36 |
| Citrobacterspp | Aminoglycosides | Amikacin | 6 | 379 | 385 | 35 |
| Citrobacterspp | Aminoglycosides | Tobramycin | 13 | 372 | 385 | 35 |
| Citrobacterspp | Penicillins | Ampicillin | 385 | 0 | 385 | 35 |
| Citrobacterspp | Penicillins | Ampicillin | 24 | 0 | 24 | 37 |
| Citrobacterspp | Penicillins | Augmentin(amoxicillin/clavulanate) | 2198 | 1579 | 3777 | 36 |
| Citrobacterspp | Penicillins | Unasyn(ampicillin/sulbactam) | 221 | 164 | 385 | 35 |
| Citrobacterspp | Penicillins | Unasyn(ampicillin/sulbactam) | 20 | 6 | 26 | 37 |
| Citrobacterspp | Penicillins | Zosyn(pipercillin/tazobactam) | 2 | 12 | 14 | 38 |
| Citrobacterspp | Penicillins | Zosyn(pipercillin/tazobactam) | 47 | 338 | 385 | 35 |
| Citrobacterspp | Penicillins | Zosyn(pipercillin/tazobactam) | 16 | 182 | 198 | 39 |

| | | | | | | |
|---|---|---|---|---|---|---|
| Citrobacterspp | Carbapenems | Doripenem | 45 | 3732 | 3777 | 36 |
| Citrobacterspp | Carbapenems | Imipenem | 198 | 23 | 221 | 40 |
| Citrobacterspp | Carbapenems | Imipenem | 11 | 3766 | 3777 | 36 |
| Citrobacterspp | Carbapenems | Meropenem | 198 | 23 | 221 | 40 |
| Citrobacterspp | Carbapenems | Meropenem | 11 | 3766 | 3777 | 36 |
| Citrobacterspp | Cephalosporins(2ndgen ) | Cefoxitin | 7 | 9 | 16 | 38 |
| Citrobacterspp | Cephalosporins(2ndgen ) | Cefoxitin | 213 | 172 | 385 | 35 |
| Citrobacterspp | Cephalosporins(3rdgen ) | Ceftazidime | 816 | 2961 | 3777 | 36 |
| Citrobacterspp | Cephalosporins(3rdgen ) | Ceftazidime | 20 | 7 | 27 | 37 |
| Citrobacterspp | Cephalosporins(3rdgen ) | Ceftazidime | 55 | 330 | 385 | 35 |
| Citrobacterspp | Cephalosporins(3rdgen ) | Ceftriaxone | 19 | 6 | 25 | 37 |
| Citrobacterspp | Cephalosporins(3rdgen ) | Ceftriaxone | 55 | 330 | 385 | 35 |
| Citrobacterspp | Tetracyclines | Doxycyline | 20 | 178 | 198 | 39 |
| Citrobacterspp | Tetracyclines | Minocycline | 18 | 180 | 198 | 39 |
| Citrobacterspp | Tetracyclines | Tetracycline | 25 | 173 | 198 | 39 |
| Citrobacterspp | Tetracyclines | Tetracycline | 22 | 363 | 385 | 35 |
| Citrobacterspp | Trimethoprim-sulfamethoxazole | Trimethoprim-sulfamethoxazole | 29 | 356 | 385 | 35 |
| Clostridiumdifficile | Glycopeptideantibiotics | Vancomycin | 0 | 70 | 70 | 10 |
| Clostridiumdifficile | Glycopeptideantibiotics | Vancomycin | 0 | 414 | 414 | 41 |

| | | | | | | |
|---|---|---|---|---|---|---|
| Clostridiumdifficile | Glycopeptideantibiotics | Vancomycin | 0 | 174 | 174 | 42 |
| Clostridiumdifficile | Glycopeptideantibiotics | Teicoplanin | 0 | 146 | 146 | 43 |
| Clostridiumperfringens | Penicillins | Penicillin | 8 | 305 | 313 | 44 |
| Clostridiumperfringens | Penicillins | Penicillin | 7 | 72 | 79 | 45 |
| Clostridiumperfringens | Penicillins | Unasyn(ampicillin/sulbactam) | 0 | 15 | 15 | 46 |
| Clostridiumperfringens | Penicillins | Zosyn(pipercillin/tazobactam) | 0 | 15 | 15 | 46 |
| Clostridiumspp | Penicillins | Penicillin | 3 | 31 | 34 | 9 |
| Clostridiumspp | Penicillins | Penicillin | 4 | 55 | 59 | 44 |
| Clostridiumspp | Penicillins | Augmentin(amoxicillin/clavulanate) | 1 | 33 | 34 | 9 |
| Clostridiumspp | Penicillins | Unasyn(ampicillin/sulbactam) | 0 | 22 | 22 | 46 |
| Clostridiumspp | Penicillins | Zosyn(pipercillin/tazobactam) | 2 | 20 | 22 | 46 |
| Clostridiumspp | Carbapenems | Imipenem | 0 | 34 | 34 | 9 |
| Clostridiumspp | Carbapenems | Imipenem | 0 | 22 | 22 | 46 |
| Clostridiumspp | Carbapenems | Meropenem | 0 | 22 | 22 | 46 |
| Corynebacteriumdiptheriae | Penicillins | Penicillin | 0 | 45 | 45 | 47 |
| Corynebacteriumdiptheriae | Penicillins | Penicillin | 0 | 108 | 108 | 48 |
| Corynebacteriumdiptheriae | Penicillins | Penicillin | 0 | 41 | 41 | 49 |
| Corynebacteriumdiptheriae | Macrolides | Erythromycin | 3 | 105 | 108 | 48 |
| Corynebacteriumdiptheriae | Macrolides | Erythromycin | 0 | 41 | 41 | 49 |

| | | | | | | |
|---|---|---|---|---|---|---|
| Corynebacteriumdiptheria e | Macrolides | Erythromycin | 0 | 157 | 157 | 50 |
| Enterobacteraerogenes | Carbapenems | Doripenem | 1 | 18 | 19 | 51 |
| Enterobacteraerogenes | Carbapenems | Imipenem | 3 | 45 | 48 | 52 |
| Enterobacteraerogenes | Carbapenems | Imipenem | 3 | 354 | 357 | 53 |
| Enterobacteraerogenes | Carbapenems | Imipenem | 2 | 17 | 19 | 51 |
| Enterobacteraerogenes | Carbapenems | Meropenem | 2 | 17 | 19 | 51 |
| Enterobacteraerogenes | Carbapenems | Ertapenem | 4 | 15 | 19 | 51 |
| Enterobacteraerogenes | Cephalosporins(3rdgen ) | Cefotaxime | 12 | 7 | 19 | 51 |
| Enterobacteraerogenes | Cephalosporins(3rdgen ) | Ceftazidime | 13 | 6 | 19 | 51 |
| Enterobacteraerogenes | Cephalosporins(4thgen ) | Cefepime | 31 | 69 | 100 | 54 |
| Enterobacteraerogenes | Cephalosporins(4thgen ) | Cefepime | 1 | 18 | 19 | 51 |
| Enterobacteraerogenes | Fluoroquinolones | Ciprofloxacin | 11 | 8 | 19 | 51 |
| Enterobacteraerogenes | Fluoroquinolones | Levofloxacin | 10 | 9 | 19 | 51 |
| Enterococcusfaecalis | Aminoglycosides | Gentamicin | 30 | 70 | 100 | 55 |
| Enterococcusfaecalis | Aminoglycosides | Gentamicin | 21 | 22 | 43 | 56 |
| Enterococcusfaecalis | Aminoglycosides | Gentamicin | 31 | 38 | 69 | 57 |
| Enterococcusfaecalis | Aminoglycosides | Amikacin | 63 | 5 | 68 | 58 |
| Enterococcusfaecalis | Aminoglycosides | Amikacin | 19 | 55 | 74 | 59 |
| Enterococcusfaecalis | Aminoglycosides | Amikacin | 0 | 10 | 10 | 60 |

| | | | | | | |
|---|---|---|---|---|---|---|
| Enterococcusfaecalis | Carbapenems | Imipenem | 0 | 63 | 63 | 61 |
| Enterococcusfaecalis | Carbapenems | Imipenem | 0 | 93 | 93 | 62 |
| Escherichiacoli | Carbapenems | Doripenem | 2 | 55 | 57 | 63 |
| Escherichiacoli | Carbapenems | Doripenem | 8 | 314 | 322 | 64 |
| Escherichiacoli | Carbapenems | Imipenem | 1 | 199 | 200 | 65 |
| Escherichiacoli | Carbapenems | Imipenem | 0 | 157 | 157 | 61 |
| Escherichiacoli | Carbapenems | Imipenem | 31 | 1632 | 1663 | 66 |
| Escherichiacoli | Carbapenems | Meropenem | 0 | 157 | 157 | 61 |
| Escherichiacoli | Carbapenems | Meropenem | 0 | 12 | 12 | 67 |
| Escherichiacoli | Carbapenems | Meropenem | 46 | 1617 | 1663 | 66 |
| Escherichiacoli | Carbapenems | Ertapenem | 0 | 157 | 157 | 61 |
| Escherichiacoli | Carbapenems | Ertapenem | 12 | 204 | 216 | 68 |
| Escherichiacoli | Carbapenems | Ertapenem | 3 | 54 | 57 | 63 |
| Escherichiacoli | Cephalosporins(1stgen) | Cefazolin | 25 | 153 | 178 | 69 |
| Escherichiacoli | Cephalosporins(1stgen) | Cefazolin | 28 | 28 | 56 | 63 |
| Escherichiacoli | Cephalosporins(1stgen) | Cefazolin | 447 | 251 | 698 | 70 |
| Escherichiacoli | Cephalosporins(1stgen) | Cephalothin | 0 | 47 | 47 | 71 |
| Escherichiacoli | Cephalosporins(1stgen) | Cephalothin | 44 | 16 | 60 | 72 |
| Escherichiacoli | Cephalosporins(1stgen) | Cephalothin | 38 | 89 | 127 | 73 |
| Escherichiacoli | Cephalosporins(1stgen) | Cephradine | 27 | 0 | 27 | 74 |

| | | | | | | |
|---|---|---|---|---|---|---|
| Escherichiacoli | Cephalosporins(1stgen) | Cephalexin | 2863 | 6833 | 9696 | 75 |
| Escherichiacoli | Cephalosporins(1stgen) | Cephalexin | 31 | 793 | 824 | 76 |
| Escherichiacoli | Cephalosporins(1stgen) | Cephalexin | 175 | 33 | 208 | 77 |
| Escherichiacoli | Cephalosporins(2ndgen ) | Cefmetazole | 31 | 1272 | 1303 | 78 |
| Escherichiacoli | Cephalosporins(2ndgen ) | Cefmetazole | 0 | 128 | 128 | 79 |
| Escherichiacoli | Cephalosporins(2ndgen ) | Cefoxitin | 36 | 180 | 216 | 68 |
| Escherichiacoli | Cephalosporins(2ndgen ) | Cefoxitin | 590 | 2194 | 2784 | 80 |
| Escherichiacoli | Cephalosporins(2ndgen ) | Cefoxitin | 484 | 66 | 550 | 81 |
| Escherichiacoli | Cephalosporins(2ndgen ) | Cefuroxime | 10 | 0 | 10 | 82 |
| Escherichiacoli | Cephalosporins(2ndgen ) | Cefuroxime | 56 | 15 | 71 | 83 |
| Escherichiacoli | Cephalosporins(2ndgen ) | Cefuroxime | 38 | 355 | 393 | 84 |
| Escherichiacoli | Cephalosporins(2ndgen ) | Cefaclor | 2849 | 6800 | 9649 | 75 |
| Escherichiacoli | Cephalosporins(2ndgen ) | Cefaclor | 37 | 9 | 46 | 85 |
| Escherichiacoli | Cephalosporins(2ndgen ) | Cefaclor | 28 | 7 | 35 | 86 |
| Escherichiacoli | Cephalosporins(3rdgen ) | Cefotaxime | 32 | 81 | 113 | 87 |
| Escherichiacoli | Cephalosporins(3rdgen ) | Cefotaxime | 82 | 134 | 216 | 68 |
| Escherichiacoli | Cephalosporins(3rdgen ) | Cefotaxime | 44 | 35 | 79 | 88 |

| | | | | | | |
|---|---|---|---|---|---|---|
| Escherichiacoli | Cephalosporins(3rdgen ) | Ceftazidime | 43 | 173 | 216 | 68 |
| Escherichiacoli | Cephalosporins(3rdgen ) | Ceftazidime | 630 | 1588 | 2218 | 89 |
| Escherichiacoli | Cephalosporins(3rdgen ) | Ceftazidime | 42 | 37 | 79 | 88 |
| Escherichiacoli | Cephalosporins(3rdgen ) | Ceftriaxone | 84 | 132 | 216 | 68 |
| Escherichiacoli | Cephalosporins(3rdgen ) | Ceftriaxone | 230 | 10 | 240 | 90 |
| Escherichiacoli | Cephalosporins(3rdgen ) | Ceftriaxone | 4652 | 3061 | 7713 | 91 |
| Escherichiacoli | Cephalosporins(3rdgen ) | Cefpodoxime | 60 | 40 | 100 | 92 |
| Escherichiacoli | Cephalosporins(3rdgen ) | Cefpodoxime | 409 | 289 | 698 | 70 |
| Escherichiacoli | Cephalosporins(3rdgen ) | Cefpodoxime | 182 | 14 | 196 | 93 |
| Escherichiacoli | Cephalosporins(3rdgen ) | Cefixime | 54 | 46 | 100 | 92 |
| Escherichiacoli | Cephalosporins(3rdgen ) | Cefixime | 258 | 440 | 698 | 70 |
| Escherichiacoli | Cephalosporins(3rdgen ) | Cefixime | 192 | 0 | 192 | 94 |
| Escherichiacoli | Cephalosporins(4thgen ) | Cefepime | 77 | 14 | 91 | 95 |
| Escherichiacoli | Cephalosporins(4thgen ) | Cefepime | 15 | 107 | 122 | 87 |
| Escherichiacoli | Cephalosporins(4thgen ) | Cefepime | 58 | 158 | 216 | 68 |
| Escherichiacoli | Cephalosporins(5thgen ) | Ceftaroline | 167 | 681 | 848 | 96 |
| Escherichiacoli | Cephalosporins(5thgen ) | Ceftaroline | 3 | 23 | 26 | 97 |

| | | | | | | |
|---|---|---|---|---|---|---|
| Escherichiacoli | Cephalosporins(5thgen ) | Ceftaroline | 24 | 314 | 338 | 98 |
| Escherichiacoli | Fluoroquinolones | Ciprofloxacin | 9 | 1 | 10 | 67 |
| Escherichiacoli | Fluoroquinolones | Ciprofloxacin | 109 | 107 | 216 | 68 |
| Escherichiacoli | Fluoroquinolones | Ciprofloxacin | 9 | 34 | 43 | 99 |
| Escherichiacoli | Fluoroquinolones | Levofloxacin | 121 | 200 | 321 | 100 |
| Escherichiacoli | Fluoroquinolones | Levofloxacin | 101 | 115 | 216 | 68 |
| Escherichiacoli | Fluoroquinolones | Levofloxacin | 0 | 160 | 160 | 69 |
| Escherichiacoli | Fluoroquinolones | Moxifloxacin | 775 | 208 | 983 | 101 |
| Escherichiacoli | Fluoroquinolones | Norfloxacin | 114 | 147 | 261 | 102 |
| Escherichiacoli | Fluoroquinolones | Norfloxacin | 20 | 16 | 36 | 103 |
| Escherichiacoli | Fluoroquinolones | Norfloxacin | 45 | 60 | 105 | 104 |
| Escherichiacoli | Fluoroquinolones | Ofloxacin | 27 | 4 | 31 | 105 |
| Escherichiacoli | Fluoroquinolones | Ofloxacin | 47 | 30 | 77 | 106 |
| Escherichiacoli | Fluoroquinolones | Ofloxacin | 56 | 65 | 121 | 107 |
| Francisellatularensis | Aminoglycosides | Gentamicin | 0 | 59 | 59 | 108 |
| Francisellatularensis | Aminoglycosides | Gentamicin | 0 | 128 | 128 | 109 |
| Francisellatularensis | Aminoglycosides | Tobramycin | 0 | 59 | 59 | 108 |
| Francisellatularensis | Fluoroquinolones | Ciprofloxacin | 0 | 59 | 59 | 108 |
| Francisellatularensis | Fluoroquinolones | Ciprofloxacin | 0 | 128 | 128 | 109 |
| Francisellatularensis | Fluoroquinolones | Levofloxacin | 0 | 59 | 59 | 108 |

| | | | | | | |
|---|---|---|---|---|---|---|
| Francisellatularensis | Fluoroquinolones | Moxifloxacin | 0 | 59 | 59 | 108 |
| Francisellatularensis | Fluoroquinolones | Ofloxacin | 0 | 59 | 59 | 108 |
| Francisellatularensis | Tetracyclines | Doxycyline | 0 | 59 | 59 | 108 |
| Francisellatularensis | Tetracyclines | Doxycyline | 0 | 128 | 128 | 109 |
| Fusobacteriumspp | Penicillins | Penicillin | 2 | 28 | 30 | 10 |
| Fusobacteriumspp | Penicillins | Ampicillin | 2 | 28 | 30 | 10 |
| Fusobacteriumspp | Penicillins | Augmentin(amoxicillin/clavulanate) | 1 | 29 | 30 | 10 |
| Fusobacteriumspp | Penicillins | Unasyn(ampicillin/sulbactam) | 0 | 14 | 14 | 13 |
| Fusobacteriumspp | Penicillins | Unasyn(ampicillin/sulbactam) | 0 | 19 | 19 | 14 |
| Fusobacteriumspp | Penicillins | Zosyn(pipercillin/tazobactam) | 1 | 29 | 30 | 10 |
| Fusobacteriumspp | Penicillins | Zosyn(pipercillin/tazobactam) | 0 | 14 | 14 | 13 |
| Fusobacteriumspp | Carbapenems | Doripenem | 0 | 14 | 14 | 13 |
| Fusobacteriumspp | Carbapenems | Imipenem | 1 | 29 | 30 | 10 |
| Fusobacteriumspp | Carbapenems | Imipenem | 0 | 14 | 14 | 13 |
| Fusobacteriumspp | Carbapenems | Meropenem | 1 | 29 | 30 | 10 |
| Fusobacteriumspp | Cephalosporins(2ndgen) | Cefmetazole | 0 | 14 | 14 | 13 |
| Fusobacteriumspp | Cephalosporins(2ndgen) | Cefmetazole | 0 | 19 | 19 | 14 |
| Fusobacteriumspp | Cephalosporins(2ndgen) | Cefoxitin | 1 | 29 | 30 | 10 |
| Gardnerellavaginalis | Macrolides | Clindamycin | 0 | 10 | 10 | 110 |

| | | | | | | |
|---|---|---|---|---|---|---|
| Gardnerellavaginalis | Macrolides | Clindamycin | 0 | 110 | 110 | 111 |
| Gardnerellavaginalis | Macrolides | Clindamycin | 14 | 190 | 204 | 112 |
| Gardnerellavaginalis | Nitroimidazoles | Metronidazole | 0 | 10 | 10 | 110 |
| Gardnerellavaginalis | Nitroimidazoles | Metronidazole | 30 | 80 | 110 | 111 |
| Gardnerellavaginalis | Nitroimidazoles | Metronidazole | 122 | 82 | 204 | 112 |
| Gardnerellavaginalis | Nitroimidazoles | Tinidazole | 123 | 81 | 204 | 112 |
| GPAC | Penicillins | Penicillin | 1 | 21 | 22 | 9 |
| GPAC | Penicillins | Penicillin | 27 | 47 | 74 | 113 |
| GPAC | Penicillins | Penicillin | 4 | 218 | 222 | 114 |
| GPAC | Penicillins | Augmentin(amoxicillin/clavulanate) | 0 | 22 | 22 | 9 |
| GPAC | Penicillins | Augmentin(amoxicillin/clavulanate) | 0 | 41 | 41 | 115 |
| GPAC | Penicillins | Unasyn(ampicillin/sulbactam) | 1 | 21 | 22 | 9 |
| GPAC | Penicillins | Unasyn(ampicillin/sulbactam) | 0 | 16 | 16 | 11 |
| GPAC | Penicillins | Unasyn(ampicillin/sulbactam) | 0 | 41 | 41 | 115 |
| GPAC | Penicillins | Zosyn(pipercillin/tazobactam) | 0 | 26 | 26 | 116 |
| GPAC | Penicillins | Zosyn(pipercillin/tazobactam) | 1 | 21 | 22 | 11 |
| GPAC | Penicillins | Zosyn(pipercillin/tazobactam) | 0 | 16 | 16 | 11 |
| GPAC | Carbapenems | Imipenem | 0 | 22 | 22 | 9 |
| GPAC | Carbapenems | Imipenem | 0 | 16 | 16 | 11 |
| GPAC | Carbapenems | Imipenem | 0 | 74 | 74 | 113 |

| | | | | | | |
|---|---|---|---|---|---|---|
| GPAC | Cephalosporins(2ndgen) | Cefotetan | 6 | 68 | 74 | 113 |
| GPAC | Cephalosporins(2ndgen) | Cefoxitin | 0 | 74 | 74 | 113 |
| GPAC | Cephalosporins(2ndgen) | Cefoxitin | 9 | 32 | 41 | 115 |
| GPAC | Macrolides | Clindamycin | 12 | 10 | 22 | 9 |
| GPAC | Macrolides | Clindamycin | 9 | 7 | 16 | 11 |
| GPAC | Macrolides | Clindamycin | 14 | 60 | 74 | 113 |
| GPAC | Nitroimidazoles | Metronidazole | 1 | 21 | 22 | 9 |
| GPAC | Nitroimidazoles | Metronidazole | 2 | 14 | 16 | 11 |
| GPAC | Nitroimidazoles | Metronidazole | 0 | 74 | 74 | 113 |
| GPAC | Tetracyclines | Tetracycline | 30 | 44 | 74 | 113 |
| Haemophilusinfluenzae | Penicillins | Penicillin | 0 | 18 | 18 | 117 |
| Haemophilusinfluenzae | Penicillins | Amoxicillin | 35 | 204 | 239 | 118 |
| Haemophilusinfluenzae | Penicillins | Amoxicillin | 126 | 132 | 258 | 119 |
| Haemophilusinfluenzae | Penicillins | Amoxicillin | 12 | 87 | 99 | 120 |
| Haemophilusinfluenzae | Penicillins | Ampicillin | 25 | 214 | 239 | 118 |
| Haemophilusinfluenzae | Penicillins | Ampicillin | 112 | 146 | 258 | 119 |
| Haemophilusinfluenzae | Penicillins | Ampicillin | 6 | 93 | 99 | 120 |
| Haemophilusinfluenzae | Penicillins | Augmentin(amoxicillin/clavulanate) | 2 | 237 | 239 | 118 |
| Haemophilusinfluenzae | Penicillins | Augmentin(amoxicillin/clavulanate) | 54 | 204 | 258 | 119 |

| | | | | | | |
|---|---|---|---|---|---|---|
| Haemophilusinfluenzae | Penicillins | Augmentin(amoxicillin/clavulanate) | 6 | 93 | 99 | 120 |
| Haemophilusinfluenzae | Penicillins | Unasyn(ampicillin/sulbactam) | 12 | 39 | 51 | 121 |
| Haemophilusinfluenzae | Penicillins | Unasyn(ampicillin/sulbactam) | 365 | 1708 | 2073 | 122 |
| Haemophilusinfluenzae | Penicillins | Unasyn(ampicillin/sulbactam) | 118 | 163 | 281 | 123 |
| Haemophilusinfluenzae | Penicillins | Zosyn(pipercillin/tazobactam) | 0 | 281 | 281 | 123 |
| Haemophilusinfluenzae | Carbapenems | Imipenem | 0 | 51 | 51 | 124 |
| Haemophilusinfluenzae | Carbapenems | Imipenem | 0 | 17 | 17 | 125 |
| Haemophilusinfluenzae | Carbapenems | Meropenem | 4 | 2069 | 2073 | 122 |
| Haemophilusinfluenzae | Carbapenems | Meropenem | 0 | 23 | 23 | 126 |
| Haemophilusinfluenzae | Cephalosporins(2ndgen) | Cefuroxime | 0 | 239 | 239 | 118 |
| Haemophilusinfluenzae | Cephalosporins(2ndgen) | Cefuroxime | 26 | 232 | 258 | 119 |
| Haemophilusinfluenzae | Cephalosporins(2ndgen) | Cefuroxime | 0 | 99 | 99 | 120 |
| Haemophilusinfluenzae | Cephalosporins(2ndgen) | Cefaclor | 1 | 238 | 239 | 118 |
| Haemophilusinfluenzae | Cephalosporins(2ndgen) | Cefaclor | 52 | 206 | 258 | 119 |
| Haemophilusinfluenzae | Cephalosporins(2ndgen) | Cefaclor | 1 | 98 | 99 | 120 |
| Haemophilusinfluenzae | Cephalosporins(3rdgen) | Cefotaxime | 122 | 1951 | 2073 | 122 |
| Haemophilusinfluenzae | Cephalosporins(3rdgen) | Cefotaxime | 22 | 588 | 610 | 127 |
| Haemophilusinfluenzae | Cephalosporins(3rdgen) | Cefotaxime | 0 | 92 | 92 | 128 |

| | | | | | | |
|---|---|---|---|---|---|---|
| Haemophilusinfluenzae | Cephalosporins(3rdgen ) | Cefdinir | 5 | 89 | 94 | 129 |
| Haemophilusinfluenzae | Cephalosporins(3rdgen ) | Cefdinir | 0 | 99 | 99 | 120 |
| Haemophilusinfluenzae | Cephalosporins(3rdgen ) | Ceftriaxone | 0 | 239 | 239 | 118 |
| Haemophilusinfluenzae | Cephalosporins(3rdgen ) | Ceftriaxone | 33 | 225 | 258 | 119 |
| Haemophilusinfluenzae | Cephalosporins(3rdgen ) | Ceftriaxone | 0 | 99 | 99 | 120 |
| Haemophilusinfluenzae | Cephalosporins(3rdgen ) | Cefpodoxime | 3 | 91 | 94 | 129 |
| Haemophilusinfluenzae | Cephalosporins(3rdgen ) | Cefpodoxime | 0 | 218 | 218 | 130 |
| Haemophilusinfluenzae | Cephalosporins(3rdgen ) | Cefpodoxime | 0 | 55 | 55 | 131 |
| Haemophilusinfluenzae | Cephalosporins(3rdgen ) | Cefixime | 93 | 165 | 258 | 119 |
| Haemophilusinfluenzae | Cephalosporins(3rdgen ) | Cefixime | 5 | 213 | 218 | 130 |
| Haemophilusinfluenzae | Cephalosporins(3rdgen ) | Cefixime | 0 | 77 | 77 | 132 |
| Haemophilusinfluenzae | Cephalosporins(5thgen ) | Ceftaroline | 71 | 1101 | 1172 | 126 |
| Haemophilusinfluenzae | Cephalosporins(5thgen ) | Ceftaroline | 6 | 422 | 428 | 133 |
| Haemophilusinfluenzae | Cephalosporins(5thgen ) | Ceftaroline | 0 | 487 | 487 | 97 |
| Haemophilusinfluenzae | Fluoroquinolones | Ciprofloxacin | 1 | 779 | 780 | 134 |
| Haemophilusinfluenzae | Fluoroquinolones | Ciprofloxacin | 0 | 51 | 51 | 124 |
| Haemophilusinfluenzae | Fluoroquinolones | Ciprofloxacin | 0 | 208 | 208 | 135 |

| | | | | | | |
|---|---|---|---|---|---|---|
| Haemophilusinfluenzae | Fluoroquinolones | Levofloxacin | 18 | 221 | 239 | 118 |
| Haemophilusinfluenzae | Fluoroquinolones | Levofloxacin | 61 | 197 | 258 | 119 |
| Haemophilusinfluenzae | Fluoroquinolones | Levofloxacin | 0 | 99 | 99 | 120 |
| Haemophilusinfluenzae | Fluoroquinolones | Moxifloxacin | 0 | 294 | 294 | 136 |
| Haemophilusinfluenzae | Fluoroquinolones | Moxifloxacin | 0 | 167 | 167 | 137 |
| Haemophilusinfluenzae | Fluoroquinolones | Moxifloxacin | 1 | 779 | 780 | 134 |
| Haemophilusinfluenzae | Fluoroquinolones | Ofloxacin | 1 | 706 | 707 | 134 |
| Haemophilusinfluenzae | Macrolides | Erythromycin | 11 | 13 | 24 | 138 |
| Haemophilusinfluenzae | Macrolides | Clarithromycin | 2 | 237 | 239 | 118 |
| Haemophilusinfluenzae | Macrolides | Clarithromycin | 27 | 231 | 258 | 119 |
| Haemophilusinfluenzae | Macrolides | Clarithromycin | 0 | 99 | 99 | 120 |
| Haemophilusinfluenzae | Macrolides | Azithromycin | 1 | 779 | 780 | 134 |
| Haemophilusinfluenzae | Macrolides | Azithromycin | 517 | 1100 | 1617 | 122 |
| Haemophilusinfluenzae | Macrolides | Azithromycin | 4 | 204 | 208 | 135 |
| Haemophilusinfluenzae | Trimethoprim-sulfamethoxazole | Trimethoprim-sulfamethoxazole | 67 | 172 | 239 | 118 |
| Haemophilusinfluenzae | Trimethoprim-sulfamethoxazole | Trimethoprim-sulfamethoxazole | 156 | 102 | 258 | 119 |
| Haemophilusinfluenzae | Trimethoprim-sulfamethoxazole | Trimethoprim-sulfamethoxazole | 31 | 68 | 99 | 120 |
| Klebsiellaoxytoca | Cephalosporins(5thgen) | Ceftaroline | 0 | 97 | 97 | 98 |
| Klebsiellaoxytoca | Cephalosporins(5thgen) | Ceftaroline | 14 | 60 | 74 | 139 |

| | | | | | | |
|---|---|---|---|---|---|---|
| Klebsiellapneumoniae | Carbapenems | Imipenem | 61 | 304 | 365 | 140 |
| Klebsiellapneumoniae | Carbapenems | Imipenem | 47 | 65 | 112 | 141 |
| Klebsiellapneumoniae | Carbapenems | Imipenem | 0 | 44 | 44 | 142 |
| Klebsiellapneumoniae | Carbapenems | Meropenem | 54 | 279 | 333 | 140 |
| Klebsiellapneumoniae | Carbapenems | Meropenem | 46 | 66 | 112 | 141 |
| Klebsiellapneumoniae | Carbapenems | Meropenem | 0 | 22 | 22 | 143 |
| Klebsiellapneumoniae | Carbapenems | Ertapenem | 88 | 105 | 193 | 68 |
| Klebsiellapneumoniae | Carbapenems | Ertapenem | 72 | 5069 | 5141 | 144 |
| Klebsiellapneumoniae | Carbapenems | Ertapenem | 9 | 91 | 100 | 145 |
| Klebsiellapneumoniae | Carbapenems | Imipenem/Cilastatin | 2 | 687 | 689 | 146 |
| Klebsiellapneumoniae | Cephalosporins(1stgen) | Cefazolin | 97 | 15 | 112 | 141 |
| Klebsiellapneumoniae | Cephalosporins(1stgen) | Cefazolin | 0 | 79 | 79 | 147 |
| Klebsiellapneumoniae | Cephalosporins(1stgen) | Cefazolin | 25 | 16 | 41 | 148 |
| Klebsiellapneumoniae | Cephalosporins(1stgen) | Cephalothin | 20 | 2 | 22 | 149 |
| Klebsiellapneumoniae | Cephalosporins(1stgen) | Cephalothin | 52 | 75 | 127 | 150 |
| Klebsiellapneumoniae | Cephalosporins(1stgen) | Cephalothin | 5566 | 1230 | 6796 | 151 |
| Klebsiellapneumoniae | Cephalosporins(1stgen) | Cephalothin | 15 | 11 | 26 | 152 |
| Klebsiellapneumoniae | Cephalosporins(1stgen) | Cephalexin | 11 | 11 | 22 | 153 |
| Klebsiellapneumoniae | Cephalosporins(2ndgen ) | Cefotetan | 147 | 286 | 433 | 154 |
| Klebsiellapneumoniae | Cephalosporins(2ndgen ) | Cefotetan | 17 | 200 | 217 | 70 |

| | | | | | | |
|---|---|---|---|---|---|---|
| Klebsiellapneumoniae | Cephalosporins(2ndgen ) | Cefoxitin | 71 | 37 | 108 | 141 |
| Klebsiellapneumoniae | Cephalosporins(2ndgen ) | Cefoxitin | 96 | 97 | 193 | 68 |
| Klebsiellapneumoniae | Cephalosporins(2ndgen ) | Cefoxitin | 1 | 43 | 44 | 142 |
| Klebsiellapneumoniae | Cephalosporins(2ndgen ) | Cefuroxime | 91 | 20 | 111 | 141 |
| Klebsiellapneumoniae | Cephalosporins(2ndgen ) | Cefuroxime | 20 | 2 | 22 | 149 |
| Klebsiellapneumoniae | Cephalosporins(2ndgen ) | Cefuroxime | 0 | 79 | 79 | 147 |
| Klebsiellapneumoniae | Cephalosporins(2ndgen ) | Cefaclor | 116 | 101 | 217 | 70 |
| Klebsiellapneumoniae | Cephalosporins(3rdgen ) | Cefotaxime | 147 | 46 | 193 | 68 |
| Klebsiellapneumoniae | Cephalosporins(3rdgen ) | Cefotaxime | 15 | 3 | 18 | 88 |
| Klebsiellapneumoniae | Cephalosporins(3rdgen ) | Cefotaxime | 23 | 0 | 23 | 37 |
| Klebsiellapneumoniae | Cephalosporins(3rdgen ) | Ceftazidime | 37 | 4 | 41 | 155 |
| Klebsiellapneumoniae | Cephalosporins(3rdgen ) | Ceftazidime | 21 | 10 | 31 | 141 |
| Klebsiellapneumoniae | Cephalosporins(3rdgen ) | Ceftazidime | 134 | 59 | 193 | 68 |
| Klebsiellapneumoniae | Cephalosporins(3rdgen ) | Ceftriaxone | 37 | 4 | 41 | 155 |
| Klebsiellapneumoniae | Cephalosporins(3rdgen ) | Ceftriaxone | 83 | 27 | 110 | 141 |
| Klebsiellapneumoniae | Cephalosporins(3rdgen ) | Ceftriaxone | 150 | 43 | 193 | 68 |
| Klebsiellapneumoniae | Cephalosporins(3rdgen ) | Cefpodoxime | 117 | 100 | 217 | 70 |

| | | | | | | |
|---|---|---|---|---|---|---|
| Klebsiellapneumoniae | Cephalosporins(3rdgen ) | Cefpodoxime | 71 | 99 | 170 | 156 |
| Klebsiellapneumoniae | Cephalosporins(3rdgen ) | Cefixime | 75 | 142 | 217 | 70 |
| Klebsiellapneumoniae | Cephalosporins(4thgen ) | Cefepime | 64 | 46 | 110 | 141 |
| Klebsiellapneumoniae | Cephalosporins(4thgen ) | Cefepime | 129 | 64 | 193 | 68 |
| Klebsiellapneumoniae | Cephalosporins(4thgen ) | Cefepime | 8 | 216 | 224 | 157 |
| Klebsiellapneumoniae | Cephalosporins(5thgen ) | Ceftaroline | 43 | 479 | 522 | 96 |
| Klebsiellapneumoniae | Cephalosporins(5thgen ) | Ceftaroline | 6 | 235 | 241 | 98 |
| Klebsiellapneumoniae | Cephalosporins(5thgen ) | Ceftaroline | 12 | 97 | 109 | 158 |
| Klebsiellapneumoniae | Fluoroquinolones | Ciprofloxacin | 80 | 29 | 109 | 141 |
| Klebsiellapneumoniae | Fluoroquinolones | Ciprofloxacin | 139 | 54 | 193 | 68 |
| Klebsiellapneumoniae | Fluoroquinolones | Ciprofloxacin | 39 | 93 | 132 | 159 |
| Klebsiellapneumoniae | Fluoroquinolones | Levofloxacin | 78 | 34 | 112 | 141 |
| Klebsiellapneumoniae | Fluoroquinolones | Levofloxacin | 129 | 64 | 193 | 68 |
| Klebsiellapneumoniae | Fluoroquinolones | Levofloxacin | 3 | 40 | 43 | 160 |
| Klebsiellapneumoniae | Fluoroquinolones | Ofloxacin | 5 | 18 | 23 | 58 |
| Klebsiellapneumoniae | Fluoroquinolones | Ofloxacin | 14 | 41 | 55 | 161 |
| Klebsiellaspp | Aminoglycosides | Gentamicin | 16 | 2 | 18 | 162 |
| Klebsiellaspp | Aminoglycosides | Gentamicin | 22 | 5 | 27 | 163 |
| Klebsiellaspp | Aminoglycosides | Gentamicin | 19 | 3 | 22 | 164 |

| | | | | | | |
|---|---|---|---|---|---|---|
| Klebsiellaspp | Aminoglycosides | Amikacin | 43 | 160 | 203 | 100 |
| Klebsiellaspp | Aminoglycosides | Amikacin | 4 | 7 | 11 | 103 |
| Klebsiellaspp | Aminoglycosides | Amikacin | 13 | 25 | 38 | 165 |
| Legionellapneumophila | Fluoroquinolones | Levofloxacin | 0 | 149 | 149 | 166 |
| Legionellapneumophila | Fluoroquinolones | Moxifloxacin | 0 | 149 | 149 | 166 |
| Legionellapneumophila | Macrolides | Erythromycin | 0 | 149 | 149 | 166 |
| Legionellapneumophila | Macrolides | Azithromycin | 25 | 124 | 149 | 166 |
| Listeriamonocytogenes | Carbapenems | Meropenem | 1 | 17 | 18 | 167 |
| Moraxellacatarrhalis | Penicillins | Penicillin | 35 | 2 | 37 | 168 |
| Moraxellacatarrhalis | Penicillins | Amoxicillin | 42 | 2 | 44 | 169 |
| Moraxellacatarrhalis | Penicillins | Ampicillin | 38 | 9 | 47 | 170 |
| Moraxellacatarrhalis | Penicillins | Ampicillin | 1 | 139 | 140 | 136 |
| Moraxellacatarrhalis | Penicillins | Ampicillin | 132 | 46 | 178 | 171 |
| Moraxellacatarrhalis | Penicillins | Augmentin(amoxicillin/clavulanate) | 0 | 178 | 178 | 171 |
| Moraxellacatarrhalis | Penicillins | Augmentin(amoxicillin/clavulanate) | 0 | 164 | 164 | 123 |
| Moraxellacatarrhalis | Penicillins | Augmentin(amoxicillin/clavulanate) | 0 | 147 | 147 | 172 |
| Moraxellacatarrhalis | Cephalosporins(2ndgen) | Cefuroxime | 2 | 42 | 44 | 169 |
| Moraxellacatarrhalis | Cephalosporins(2ndgen) | Cefuroxime | 25 | 160 | 185 | 173 |
| Moraxellacatarrhalis | Cephalosporins(2ndgen) | Cefuroxime | 8 | 132 | 140 | 136 |

| | | | | | | |
|---|---|---|---|---|---|---|
| Moraxellacatarrhalis | Cephalosporins(2ndgen ) | Cefaclor | 4 | 40 | 44 | 169 |
| Moraxellacatarrhalis | Cephalosporins(2ndgen ) | Cefaclor | 8 | 132 | 140 | 136 |
| Moraxellacatarrhalis | Cephalosporins(2ndgen ) | Cefaclor | 0 | 40 | 40 | 174 |
| Moraxellacatarrhalis | Fluoroquinolones | Ciprofloxacin | 0 | 164 | 164 | 123 |
| Moraxellacatarrhalis | Fluoroquinolones | Ciprofloxacin | 0 | 147 | 147 | 172 |
| Moraxellacatarrhalis | Fluoroquinolones | Ciprofloxacin | 0 | 44 | 44 | 169 |
| Moraxellacatarrhalis | Fluoroquinolones | Levofloxacin | 0 | 613 | 613 | 126 |
| Moraxellacatarrhalis | Fluoroquinolones | Levofloxacin | 0 | 100 | 100 | 175 |
| Moraxellacatarrhalis | Fluoroquinolones | Levofloxacin | 0 | 140 | 140 | 136 |
| Moraxellacatarrhalis | Fluoroquinolones | Moxifloxacin | 0 | 140 | 140 | 136 |
| Moraxellacatarrhalis | Fluoroquinolones | Moxifloxacin | 0 | 577 | 577 | 176 |
| Moraxellacatarrhalis | Macrolides | Erythromycin | 43 | 135 | 178 | 171 |
| Moraxellacatarrhalis | Macrolides | Erythromycin | 32 | 5 | 37 | 168 |
| Moraxellacatarrhalis | Macrolides | Clarithromycin | 0 | 140 | 140 | 136 |
| Moraxellacatarrhalis | Macrolides | Clarithromycin | 1 | 576 | 577 | 176 |
| Moraxellacatarrhalis | Macrolides | Clarithromycin | 0 | 197 | 197 | 177 |
| Moraxellacatarrhalis | Macrolides | Azithromycin | 26 | 18 | 44 | 169 |
| Moraxellacatarrhalis | Macrolides | Azithromycin | 0 | 140 | 140 | 136 |
| Moraxellacatarrhalis | Macrolides | Azithromycin | 49 | 171 | 220 | 178 |
| Moraxellacatarrhalis | Macrolides | Clindamycin | 27 | 17 | 44 | 169 |

| | | | | | | |
|---|---|---|---|---|---|---|
| Moraxellacatarrhalis | Tetracyclines | Doxycyline | 1 | 43 | 44 | 169 |
| Moraxellacatarrhalis | Tetracyclines | Tetracycline | 3 | 910 | 913 | 39 |
| Moraxellacatarrhalis | Tetracyclines | Tetracycline | 4 | 609 | 613 | 126 |
| Moraxellacatarrhalis | Tetracyclines | Tetracycline | 1 | 43 | 44 | 169 |
| Mycoplasmapneumoniae | Macrolides | Erythromycin | 107 | 3 | 110 | 179 |
| Mycoplasmapneumoniae | Macrolides | Erythromycin | 123 | 31 | 154 | 180 |
| Mycoplasmapneumoniae | Macrolides | Erythromycin | 53 | 28 | 81 | 181 |
| Mycoplasmapneumoniae | Macrolides | Azithromycin | 107 | 3 | 110 | 179 |
| Mycoplasmapneumoniae | Macrolides | Azithromycin | 123 | 31 | 154 | 180 |
| Mycoplasmapneumoniae | Macrolides | Azithromycin | 53 | 28 | 81 | 181 |
| Mycoplasmapneumoniae | Tetracyclines | Minocycline | 0 | 75 | 75 | 182 |
| Mycoplasmapneumoniae | Tetracyclines | Tetracycline | 0 | 81 | 81 | 180 |
| Mycoplasmapneumoniae | Tetracyclines | Tetracycline | 0 | 75 | 75 | 182 |
| Neisseriagonorrhoeae | Carbapenems | Ertapenem | 4 | 39 | 43 | 183 |
| Neisseriagonorrhoeae | Cephalosporins(2ndgen ) | Cefoxitin | 0 | 21 | 21 | 184 |
| Neisseriagonorrhoeae | Cephalosporins(2ndgen ) | Cefuroxime | 0 | 64 | 64 | 185 |
| Neisseriagonorrhoeae | Cephalosporins(3rdgen ) | Cefotaxime | 0 | 80 | 80 | 186 |
| Neisseriagonorrhoeae | Cephalosporins(3rdgen ) | Cefotaxime | 532 | 11236 | 11768 | 187 |
| Neisseriagonorrhoeae | Cephalosporins(3rdgen ) | Cefotaxime | 13 | 596 | 609 | 183 |

| | | | | | | |
|---|---|---|---|---|---|---|
| Neisseriagonorrhoeae | Cephalosporins(3rdgen ) | Ceftazidime | 0 | 300 | 300 | 188 |
| Neisseriagonorrhoeae | Cephalosporins(3rdgen ) | Ceftriaxone | 2 | 395 | 397 | 189 |
| Neisseriagonorrhoeae | Cephalosporins(3rdgen ) | Ceftriaxone | 0 | 2596 | 2596 | 190 |
| Neisseriagonorrhoeae | Cephalosporins(3rdgen ) | Ceftriaxone | 0 | 361 | 361 | 191 |
| Neisseriagonorrhoeae | Cephalosporins(3rdgen ) | Cefpodoxime | 0 | 379 | 379 | 192 |
| Neisseriagonorrhoeae | Cephalosporins(3rdgen ) | Cefpodoxime | 30 | 196 | 226 | 193 |
| Neisseriagonorrhoeae | Cephalosporins(3rdgen ) | Cefpodoxime | 105 | 985 | 1090 | 194 |
| Neisseriagonorrhoeae | Cephalosporins(3rdgen ) | Cefixime | 0 | 2596 | 2596 | 190 |
| Neisseriagonorrhoeae | Cephalosporins(3rdgen ) | Cefixime | 0 | 150 | 150 | 195 |
| Neisseriagonorrhoeae | Cephalosporins(3rdgen ) | Cefixime | 19 | 210 | 229 | 196 |
| Neisseriameningitidis | Penicillins | Penicillin | 3 | 209 | 212 | 197 |
| Neisseriameningitidis | Penicillins | Penicillin | 14 | 262 | 276 | 197 |
| Neisseriameningitidis | Penicillins | Penicillin | 0 | 2888 | 2888 | 198 |
| Neisseriameningitidis | Penicillins | Amoxicillin | 0 | 107 | 107 | 199 |
| Neisseriameningitidis | Penicillins | Ampicillin | 0 | 2888 | 2888 | 198 |
| Neisseriameningitidis | Penicillins | Ampicillin | 0 | 25 | 25 | 200 |
| Neisseriameningitidis | Carbapenems | Meropenem | 0 | 25 | 25 | 200 |
| Neisseriameningitidis | Cephalosporins(3rdgen ) | Cefotaxime | 1 | 73 | 74 | 201 |

| | | | | | | |
|---|---|---|---|---|---|---|
| Neisseriameningitidis | Cephalosporins(3rdgen ) | Ceftriaxone | 0 | 210 | 210 | 197 |
| Neisseriameningitidis | Cephalosporins(3rdgen ) | Ceftriaxone | 0 | 274 | 274 | 197 |
| Neisseriameningitidis | Cephalosporins(3rdgen ) | Ceftriaxone | 0 | 2888 | 2888 | 198 |
| Nocardiaspp | Carbapenems | Imipenem | 160 | 110 | 270 | 202 |
| Nocardiaspp | Carbapenems | Imipenem | 5 | 48 | 53 | 203 |
| Nocardiaspp | Carbapenems | Imipenem | 13 | 39 | 52 | 204 |
| Nocardiaspp | Trimethoprim-sulfamethoxazole | Trimethoprim-sulfamethoxazole | 0 | 18 | 18 | 205 |
| Nocardiaspp | Trimethoprim-sulfamethoxazole | Trimethoprim-sulfamethoxazole | 25 | 245 | 270 | 202 |
| Nocardiaspp | Trimethoprim-sulfamethoxazole | Trimethoprim-sulfamethoxazole | 14 | 39 | 53 | 203 |
| Non-typhoidalSalmonella | Cephalosporins(3rdgen ) | Cefotaxime | 42 | 136 | 178 | 206 |
| Non-typhoidalSalmonella | Cephalosporins(3rdgen ) | Cefotaxime | 5 | 132 | 137 | 207 |
| Non-typhoidalSalmonella | Cephalosporins(3rdgen ) | Cefotaxime | 14 | 19 | 33 | 208 |
| Non-typhoidalSalmonella | Cephalosporins(3rdgen ) | Ceftazidime | 9 | 128 | 137 | 207 |
| Non-typhoidalSalmonella | Cephalosporins(3rdgen ) | Ceftriaxone | 136 | 728 | 864 | 209 |
| Non-typhoidalSalmonella | Cephalosporins(3rdgen ) | Ceftriaxone | 43 | 243 | 286 | 210 |
| Non-typhoidalSalmonella | Cephalosporins(3rdgen ) | Ceftriaxone | 0 | 826 | 826 | 211 |
| Non-typhoidalSalmonella | Macrolides | Azithromycin | 129 | 735 | 864 | 209 |

| | | | | | | |
|---|---|---|---|---|---|---|
| Non-typhoidalSalmonella | Macrolides | Azithromycin | 22 | 11 | 33 | 208 |
| Non-typhoidalSalmonella | Macrolides | Azithromycin | 6 | 48 | 54 | 212 |
| Propiniobacteriumacnes | Penicillins | Penicillin | 0 | 22 | 22 | 213 |
| Propiniobacteriumacnes | Penicillins | Penicillin | 4 | 62 | 66 | 214 |
| Propiniobacteriumacnes | Penicillins | Penicillin | 0 | 106 | 106 | 215 |
| Propiniobacteriumacnes | Penicillins | Ampicillin | 0 | 63 | 63 | 216 |
| Propiniobacteriumacnes | Penicillins | Augmentin(amoxicillin/clavulanate) | 0 | 17 | 17 | 115 |
| Propiniobacteriumacnes | Penicillins | Unasyn(ampicillin/sulbactam) | 0 | 63 | 63 | 216 |
| Propiniobacteriumacnes | Penicillins | Unasyn(ampicillin/sulbactam) | 0 | 17 | 17 | 115 |
| Propiniobacteriumacnes | Carbapenems | Imipenem | 0 | 63 | 63 | 216 |
| Propiniobacteriumacnes | Carbapenems | Imipenem | 0 | 17 | 17 | 115 |
| Propiniobacteriumacnes | Cephalosporins(2ndgen) | Cefoxitin | 0 | 63 | 63 | 216 |
| Propiniobacteriumacnes | Cephalosporins(2ndgen) | Cefoxitin | 0 | 17 | 17 | 115 |
| Propiniobacteriumacnes | Cephalosporins(3rdgen) | Ceftriaxone | 0 | 22 | 22 | 213 |
| Propiniobacteriumacnes | Cephalosporins(3rdgen) | Ceftriaxone | 0 | 63 | 63 | 216 |
| Propiniobacteriumacnes | Cephalosporins(3rdgen) | Ceftriaxone | 0 | 1005 | 1005 | 217 |
| Propiniobacteriumacnes | Macrolides | Erythromycin | 20 | 116 | 136 | 217 |
| Propiniobacteriumacnes | Macrolides | Erythromycin | 31 | 32 | 63 | 216 |
| Propiniobacteriumacnes | Macrolides | Erythromycin | 34 | 35 | 69 | 218 |

| | | | | | | |
|---|---|---|---|---|---|---|
| Propiniobacteriumacnes | Macrolides | Clindamycin | 41 | 964 | 1005 | 217 |
| Propiniobacteriumacnes | Macrolides | Clindamycin | 2 | 20 | 22 | 213 |
| Propiniobacteriumacnes | Macrolides | Clindamycin | 0 | 63 | 63 | 216 |
| Propiniobacteriumacnes | Nitroimidazoles | Metronidazole | 63 | 0 | 63 | 216 |
| Propiniobacteriumacnes | Tetracyclines | Minocycline | 0 | 63 | 63 | 216 |
| Propiniobacteriumacnes | Tetracyclines | Minocycline | 0 | 66 | 66 | 214 |
| Propiniobacteriumacnes | Tetracyclines | Tetracycline | 3 | 133 | 136 | 217 |
| Propiniobacteriumacnes | Tetracyclines | Tetracycline | 6 | 60 | 66 | 214 |
| Proteusmirabilis | Cephalosporins(1stgen) | Cefazolin | 53 | 17 | 70 | 141 |
| Proteusmirabilis | Cephalosporins(1stgen) | Cefazolin | 6 | 36 | 42 | 219 |
| Proteusmirabilis | Cephalosporins(1stgen) | Cefazolin | 86 | 15 | 101 | 220 |
| Proteusmirabilis | Cephalosporins(1stgen) | Cephalothin | 3930 | 456 | 4386 | 151 |
| Proteusmirabilis | Cephalosporins(1stgen) | Cephalothin | 4 | 6 | 10 | 152 |
| Proteusmirabilis | Cephalosporins(1stgen) | Cephalothin | 24 | 38 | 62 | 153 |
| Proteusmirabilis | Cephalosporins(1stgen) | Cephalexin | 21 | 42 | 63 | 221 |
| Proteusmirabilis | Cephalosporins(1stgen) | Cephalexin | 19 | 43 | 62 | 153 |
| Proteusmirabilis | Cephalosporins(2ndgen) | Cefoxitin | 10 | 60 | 70 | 141 |
| Proteusmirabilis | Cephalosporins(2ndgen) | Cefoxitin | 667 | 3719 | 4386 | 151 |
| Proteusmirabilis | Cephalosporins(2ndgen) | Cefoxitin | 0 | 0 | 0 | 152 |

| | | | | | | |
|---|---|---|---|---|---|---|
| Proteusmirabilis | Cephalosporins(2ndgen ) | Cefuroxime | 47 | 22 | 69 | 141 |
| Proteusmirabilis | Cephalosporins(2ndgen ) | Cefuroxime | 0 | 960 | 960 | 222 |
| Proteusmirabilis | Cephalosporins(2ndgen ) | Cefuroxime | 0 | 10 | 10 | 223 |
| Proteusmirabilis | Cephalosporins(3rdgen ) | Cefotaxime | 16 | 150 | 166 | 224 |
| Proteusmirabilis | Cephalosporins(3rdgen ) | Cefotaxime | 0 | 4386 | 4386 | 151 |
| Proteusmirabilis | Cephalosporins(3rdgen ) | Ceftazidime | 72 | 1073 | 1145 | 225 |
| Proteusmirabilis | Cephalosporins(3rdgen ) | Ceftazidime | 6 | 20 | 26 | 141 |
| Proteusmirabilis | Cephalosporins(3rdgen ) | Ceftazidime | 5 | 96 | 101 | 220 |
| Proteusmirabilis | Cephalosporins(3rdgen ) | Ceftriaxone | 40 | 29 | 69 | 141 |
| Proteusmirabilis | Cephalosporins(3rdgen ) | Ceftriaxone | 1 | 62 | 63 | 221 |
| Proteusmirabilis | Cephalosporins(3rdgen ) | Ceftriaxone | 0 | 1117 | 1117 | 226 |
| Proteusmirabilis | Cephalosporins(3rdgen ) | Cefixime | 2 | 14 | 16 | 227 |
| Proteusmirabilis | Cephalosporins(3rdgen ) | Cefixime | 17 | 159 | 176 | 222 |
| Proteusmirabilis | Cephalosporins(4thgen ) | Cefepime | 23 | 47 | 70 | 141 |
| Proteusmirabilis | Cephalosporins(4thgen ) | Cefepime | 0 | 33 | 33 | 157 |
| Proteusmirabilis | Cephalosporins(4thgen ) | Cefepime | 6 | 95 | 101 | 220 |
| Proteusmirabilis | Fluoroquinolones | Ciprofloxacin | 58 | 12 | 70 | 141 |

| | | | | | | |
|---|---|---|---|---|---|---|
| Proteusmirabilis | Fluoroquinolones | Ciprofloxacin | 8 | 55 | 63 | 221 |
| Proteusmirabilis | Fluoroquinolones | Ciprofloxacin | 418 | 5549 | 5967 | 228 |
| Proteusmirabilis | Fluoroquinolones | Levofloxacin | 58 | 12 | 70 | 141 |
| Proteusmirabilis | Fluoroquinolones | Levofloxacin | 5 | 50 | 55 | 160 |
| Proteusmirabilis | Fluoroquinolones | Levofloxacin | 16 | 115 | 131 | 222 |
| Proteusmirabilis | Fluoroquinolones | Moxifloxacin | 3 | 7 | 10 | 152 |
| Proteusmirabilis | Fluoroquinolones | Norfloxacin | 7 | 56 | 63 | 221 |
| Proteusmirabilis | Fluoroquinolones | Norfloxacin | 0 | 16 | 16 | 227 |
| Proteusmirabilis | Fluoroquinolones | Ofloxacin | 302 | 815 | 1117 | 226 |
| Proteusspp | Carbapenems | Imipenem | 0 | 49 | 49 | 222 |
| Proteusspp | Carbapenems | Imipenem | 1 | 11 | 12 | 229 |
| Proteusspp | Carbapenems | Imipenem | 4 | 97 | 101 | 220 |
| Proteusspp | Carbapenems | Meropenem | 0 | 16 | 16 | 230 |
| Proteusspp | Carbapenems | Meropenem | 9 | 20 | 29 | 231 |
| Proteusspp | Carbapenems | Meropenem | 0 | 52 | 52 | 232 |
| Pseudomonasaeruginosa | Aminoglycosides | Gentamicin | 274 | 1117 | 1391 | 233 |
| Pseudomonasaeruginosa | Aminoglycosides | Gentamicin | 72 | 95 | 167 | 141 |
| Pseudomonasaeruginosa | Aminoglycosides | Gentamicin | 0 | 227 | 227 | 234 |
| Pseudomonasaeruginosa | Aminoglycosides | Amikacin | 15 | 177 | 192 | 235 |
| Pseudomonasaeruginosa | Aminoglycosides | Amikacin | 548 | 5608 | 6156 | 225 |

| | | | | | | |
|---|---|---|---|---|---|---|
| Pseudomonasaeruginosa | Aminoglycosides | Amikacin | 37 | 42 | 79 | 236 |
| Pseudomonasaeruginosa | Aminoglycosides | Tobramycin | 0 | 227 | 227 | 234 |
| Pseudomonasaeruginosa | Aminoglycosides | Tobramycin | 20 | 747 | 767 | 134 |
| Pseudomonasaeruginosa | Aminoglycosides | Tobramycin | 58 | 356 | 414 | 237 |
| Pseudomonasaeruginosa | Aminoglycosides | Plazomicin | 25 | 77 | 102 | 238 |
| Pseudomonasaeruginosa | Aminoglycosides | Plazomicin | 29 | 74 | 103 | 239 |
| Pseudomonasaeruginosa | Carbapenems | Doripenem | 56 | 377 | 433 | 240 |
| Pseudomonasaeruginosa | Carbapenems | Doripenem | 10 | 95 | 105 | 241 |
| Pseudomonasaeruginosa | Carbapenems | Doripenem | 15 | 155 | 170 | 242 |
| Pseudomonasaeruginosa | Carbapenems | Imipenem | 0 | 149 | 149 | 140 |
| Pseudomonasaeruginosa | Carbapenems | Imipenem | 17 | 175 | 192 | 235 |
| Pseudomonasaeruginosa | Carbapenems | Imipenem | 1674 | 4482 | 6156 | 225 |
| Pseudomonasaeruginosa | Carbapenems | Meropenem | 37 | 30 | 67 | 243 |
| Pseudomonasaeruginosa | Carbapenems | Meropenem | 0 | 141 | 141 | 140 |
| Pseudomonasaeruginosa | Carbapenems | Meropenem | 348 | 131 | 479 | 244 |
| Pseudomonasaeruginosa | Carbapenems | Ertapenem | 0 | 13 | 13 | 245 |
| Pseudomonasaeruginosa | Carbapenems | Ertapenem | 5746 | 0 | 5746 | 151 |
| Pseudomonasaeruginosa | Carbapenems | Imipenem/Cilastatin | 25 | 871 | 896 | 246 |
| Pseudomonasaeruginosa | Carbapenems | Imipenem/Cilastatin | 22 | 823 | 845 | 146 |
| Pseudomonasaeruginosa | Cephalosporins(3rdgen ) | Cefotaxime | 41 | 5 | 46 | 247 |

| | | | | | | |
|---|---|---|---|---|---|---|
| Pseudomonasaeruginosa | Cephalosporins(3rdgen ) | Cefotaxime | 37 | 21 | 58 | 248 |
| Pseudomonasaeruginosa | Cephalosporins(3rdgen ) | Cefotaxime | 68 | 7 | 75 | 249 |
| Pseudomonasaeruginosa | Cephalosporins(3rdgen ) | Ceftazidime | 1404 | 4752 | 6156 | 225 |
| Pseudomonasaeruginosa | Cephalosporins(3rdgen ) | Ceftazidime | 51 | 116 | 167 | 141 |
| Pseudomonasaeruginosa | Cephalosporins(3rdgen ) | Ceftazidime | 91 | 136 | 227 | 234 |
| Pseudomonasaeruginosa | Cephalosporins(3rdgen ) | Ceftriaxone | 5 | 35 | 40 | 250 |
| Pseudomonasaeruginosa | Cephalosporins(3rdgen ) | Ceftriaxone | 15 | 13 | 28 | 251 |
| Pseudomonasaeruginosa | Cephalosporins(3rdgen ) | Ceftriaxone | 29 | 2 | 31 | 37 |
| Pseudomonasaeruginosa | Cephalosporins(4thgen ) | Cefepime | 211 | 16 | 227 | 234 |
| Pseudomonasaeruginosa | Cephalosporins(4thgen ) | Cefepime | 37 | 130 | 167 | 141 |
| Pseudomonasaeruginosa | Cephalosporins(4thgen ) | Cefepime | 34 | 98 | 132 | 68 |
| Salmonellatyphi | Carbapenems | Doripenem | 3 | 125 | 128 | 252 |
| Salmonellatyphi | Carbapenems | Imipenem | 0 | 43 | 43 | 253 |
| Salmonellatyphi | Carbapenems | Imipenem | 0 | 128 | 128 | 252 |
| Salmonellatyphi | Carbapenems | Meropenem | 0 | 42 | 42 | 254 |
| Salmonellatyphi | Carbapenems | Meropenem | 0 | 239 | 239 | 221 |
| Salmonellatyphi | Carbapenems | Ertapenem | 4 | 124 | 128 | 252 |
| Salmonellatyphi | Cephalosporins(3rdgen ) | Cefotaxime | 0 | 135 | 135 | 255 |

| | | | | | | |
|---|---|---|---|---|---|---|
| Salmonellatyphi | Cephalosporins(3rdgen ) | Cefotaxime | 0 | 144 | 144 | 256 |
| Salmonellatyphi | Cephalosporins(3rdgen ) | Cefotaxime | 0 | 78 | 78 | 257 |
| Salmonellatyphi | Cephalosporins(3rdgen ) | Ceftazidime | 9 | 33 | 42 | 254 |
| Salmonellatyphi | Cephalosporins(3rdgen ) | Ceftriaxone | 0 | 133 | 133 | 258 |
| Salmonellatyphi | Cephalosporins(3rdgen ) | Ceftriaxone | 423 | 42996 | 43419 | 259 |
| Salmonellatyphi | Cephalosporins(3rdgen ) | Ceftriaxone | 1 | 41 | 42 | 254 |
| Salmonellatyphi | Cephalosporins(3rdgen ) | Cefpodoxime | 6 | 122 | 128 | 252 |
| Salmonellatyphi | Cephalosporins(3rdgen ) | Cefixime | 0 | 133 | 133 | 258 |
| Salmonellatyphi | Cephalosporins(3rdgen ) | Cefixime | 0 | 192 | 192 | 260 |
| Salmonellatyphi | Cephalosporins(3rdgen ) | Cefixime | 0 | 395 | 395 | 261 |
| Salmonellatyphi | Fluoroquinolones | Ciprofloxacin | 130 | 3 | 133 | 258 |
| Salmonellatyphi | Fluoroquinolones | Ciprofloxacin | 0 | 42 | 42 | 254 |
| Salmonellatyphi | Fluoroquinolones | Ciprofloxacin | 3 | 390 | 393 | 262 |
| Salmonellatyphi | Fluoroquinolones | Levofloxacin | 0 | 42 | 42 | 254 |
| Salmonellatyphi | Fluoroquinolones | Levofloxacin | 6 | 68 | 74 | 263 |
| Salmonellatyphi | Fluoroquinolones | Levofloxacin | 3 | 90 | 93 | 264 |
| Salmonellatyphi | Fluoroquinolones | Moxifloxacin | 4 | 124 | 128 | 252 |
| Salmonellatyphi | Fluoroquinolones | Ofloxacin | 6 | 36 | 42 | 254 |

| | | | | | | |
|---|---|---|---|---|---|---|
| Salmonellatyphi | Fluoroquinolones | Ofloxacin | 8 | 154 | 162 | 265 |
| Salmonellatyphi | Fluoroquinolones | Ofloxacin | 144 | 11 | 155 | 266 |
| Serratiamarcescens | Carbapenems | Imipenem | 1 | 44 | 45 | 267 |
| Serratiamarcescens | Carbapenems | Imipenem | 0 | 30 | 30 | 268 |
| Serratiamarcescens | Carbapenems | Imipenem | 0 | 738 | 738 | 269 |
| Serratiamarcescens | Carbapenems | Meropenem | 15 | 25 | 40 | 270 |
| Serratiamarcescens | Carbapenems | Meropenem | 45 | 2912 | 2957 | 271 |
| Serratiamarcescens | Carbapenems | Meropenem | 24 | 5982 | 6006 | 269 |
| Serratiamarcescens | Cephalosporins(3rdgen ) | Cefotaxime | 16 | 0 | 16 | 272 |
| Serratiamarcescens | Cephalosporins(3rdgen ) | Cefotaxime | 1 | 157 | 158 | 273 |
| Serratiamarcescens | Cephalosporins(3rdgen ) | Cefotaxime | 1 | 44 | 45 | 267 |
| Serratiamarcescens | Cephalosporins(3rdgen ) | Ceftazidime | 31 | 127 | 158 | 273 |
| Serratiamarcescens | Cephalosporins(3rdgen ) | Ceftazidime | 15 | 25 | 40 | 270 |
| Serratiamarcescens | Cephalosporins(3rdgen ) | Ceftriaxone | 36 | 122 | 158 | 273 |
| Serratiamarcescens | Cephalosporins(3rdgen ) | Ceftriaxone | 372 | 2585 | 2957 | 271 |
| Serratiamarcescens | Cephalosporins(3rdgen ) | Ceftriaxone | 15 | 25 | 40 | 270 |
| Serratiamarcescens | Fluoroquinolones | Ciprofloxacin | 1 | 44 | 45 | 267 |
| Serratiamarcescens | Fluoroquinolones | Ciprofloxacin | 0 | 30 | 30 | 268 |
| Serratiamarcescens | Fluoroquinolones | Ciprofloxacin | 2 | 38 | 40 | 270 |

| | | | | | | |
|---|---|---|---|---|---|---|
| Serratiamarcescens | Fluoroquinolones | Levofloxacin | 0 | 30 | 30 | 268 |
| Serratiamarcescens | Fluoroquinolones | Levofloxacin | 15 | 25 | 40 | 270 |
| Serratiamarcescens | Fluoroquinolones | Levofloxacin | 398 | 6346 | 6744 | 269 |
| Shigellaspecies | Fluoroquinolones | Ciprofloxacin | 5 | 108 | 113 | 274 |
| Shigellaspecies | Fluoroquinolones | Ciprofloxacin | 0 | 18 | 18 | 275 |
| Shigellaspecies | Fluoroquinolones | Ciprofloxacin | 0 | 26 | 26 | 276 |
| Shigellaspecies | Fluoroquinolones | Levofloxacin | 0 | 50 | 50 | 277 |
| Shigellaspecies | Fluoroquinolones | Norfloxacin | 1 | 18 | 19 | 278 |
| Shigellaspecies | Fluoroquinolones | Norfloxacin | 0 | 50 | 50 | 277 |
| Shigellaspecies | Fluoroquinolones | Norfloxacin | 0 | 17 | 17 | 279 |
| Shigellaspecies | Fluoroquinolones | Ofloxacin | 0 | 17 | 17 | 279 |
| Shigellaspecies | Macrolides | Azithromycin | 15 | 178 | 193 | 280 |
| Shigellaspecies | Macrolides | Azithromycin | 9 | 32 | 41 | 281 |
| Shigellaspecies | Macrolides | Azithromycin | 7 | 337 | 344 | 282 |
| Staphylococcusaureus | Carbapenems | Imipenem | 5 | 114 | 119 | 283 |
| Staphylococcusaureus | Carbapenems | Imipenem | 0 | 45 | 45 | 284 |
| Staphylococcusaureus | Carbapenems | Imipenem | 0 | 24 | 24 | 285 |
| Staphylococcusaureus | Cephalosporins(1stgen) | Cefazolin | 11 | 127 | 138 | 286 |
| Staphylococcusaureus | Cephalosporins(1stgen) | Cefazolin | 17 | 88 | 105 | 287 |
| Staphylococcusaureus | Cephalosporins(1stgen) | Cefazolin | 0 | 7146 | 7146 | 288 |

| | | | | | | |
|---|---|---|---|---|---|---|
| Staphylococcusaureus | Cephalosporins(1stgen) | Cephalothin | 47 | 32 | 79 | 289 |
| Staphylococcusaureus | Cephalosporins(1stgen) | Cephalexin | 321 | 59 | 380 | 290 |
| Staphylococcusaureus | Cephalosporins(1stgen) | Cephalexin | 7 | 3 | 10 | 291 |
| Staphylococcusaureus | Cephalosporins(2ndgen ) | Cefoxitin | 16 | 34 | 50 | 292 |
| Staphylococcusaureus | Cephalosporins(2ndgen ) | Cefoxitin | 0 | 194 | 194 | 293 |
| Staphylococcusaureus | Cephalosporins(2ndgen ) | Cefoxitin | 377 | 331 | 708 | 5 |
| Staphylococcusaureus | Cephalosporins(2ndgen ) | Cefuroxime | 47 | 13 | 60 | 294 |
| Staphylococcusaureus | Cephalosporins(2ndgen ) | Cefuroxime | 0 | 16 | 16 | 295 |
| Staphylococcusaureus | Cephalosporins(2ndgen ) | Cefuroxime | 52 | 82 | 134 | 33 |
| Staphylococcusaureus | Cephalosporins(4thgen ) | Cefepime | 0 | 100 | 100 | 157 |
| Staphylococcusaureus | Cephalosporins(5thgen ) | Ceftaroline | 0 | 67 | 67 | 296 |
| Staphylococcusaureus | Cephalosporins(5thgen ) | Ceftaroline | 0 | 80 | 80 | 297 |
| Staphylococcusaureus | Cephalosporins(5thgen ) | Ceftaroline | 25 | 25183 | 25208 | 298 |
| Staphylococcusaureus | Macrolides | Erythromycin | 93 | 226 | 319 | 299 |
| Staphylococcusaureus | Macrolides | Erythromycin | 23 | 130 | 153 | 300 |
| Staphylococcusaureus | Macrolides | Erythromycin | 14 | 36 | 50 | 292 |
| Staphylococcusaureus | Macrolides | Clarithromycin | 2 | 10 | 12 | 227 |
| Staphylococcusaureus | Macrolides | Azithromycin | 572 | 852 | 1424 | 134 |

| | | | | | | |
|---|---|---|---|---|---|---|
| Staphylococcusaureus | Macrolides | Azithromycin | 115 | 71 | 186 | 301 |
| Staphylococcusaureus | Macrolides | Azithromycin | 129 | 201 | 330 | 135 |
| Staphylococcusaureus | Macrolides | Clindamycin | 24 | 20 | 44 | 81 |
| Staphylococcusaureus | Macrolides | Clindamycin | 132 | 29 | 161 | 141 |
| Staphylococcusaureus | Macrolides | Clindamycin | 14 | 139 | 153 | 300 |
| Staphylococcusepidermidis | Carbapenems | Meropenem | 18 | 0 | 18 | 302 |
| Staphylococcusepidermidis | Cephalosporins(1stgen) | Cefazolin | 36 | 10 | 46 | 303 |
| Streptococcusagalactiae | Aminoglycosides | Gentamicin | 24 | 8733 | 8757 | 304 |
| Streptococcusagalactiae | Aminoglycosides | Amikacin | 572 | 8185 | 8757 | 304 |
| Streptococcusagalactiae | Cephalosporins(5thgen ) | Ceftaroline | 0 | 44 | 44 | 98 |
| Streptococcuspneumoniae | Penicillins | Penicillin | 7 | 37 | 44 | 305 |
| Streptococcuspneumoniae | Penicillins | Penicillin | 0 | 14 | 14 | 306 |
| Streptococcuspneumoniae | Penicillins | Penicillin | 77 | 102 | 179 | 118 |
| Streptococcuspneumoniae | Penicillins | Oxacillin | 12 | 299 | 311 | 307 |
| Streptococcuspneumoniae | Penicillins | Oxacillin | 205 | 426 | 631 | 308 |
| Streptococcuspneumoniae | Penicillins | Oxacillin | 0 | 96 | 96 | 135 |
| Streptococcuspneumoniae | Penicillins | Amoxicillin | 22 | 157 | 179 | 118 |
| Streptococcuspneumoniae | Penicillins | Amoxicillin | 54 | 206 | 260 | 119 |
| Streptococcuspneumoniae | Penicillins | Amoxicillin | 0 | 78 | 78 | 120 |
| Streptococcuspneumoniae | Penicillins | Ampicillin | 22 | 157 | 179 | 118 |

| | | | | | | |
|---|---|---|---|---|---|---|
| Streptococcuspneumoniae | Penicillins | Ampicillin | 137 | 123 | 260 | 119 |
| Streptococcuspneumoniae | Penicillins | Ampicillin | 6 | 72 | 78 | 120 |
| Streptococcuspneumoniae | Penicillins | Augmentin(amoxicillin/clavulan ate) | 1 | 178 | 179 | 118 |
| Streptococcuspneumoniae | Penicillins | Augmentin(amoxicillin/clavulan ate) | 54 | 206 | 260 | 119 |
| Streptococcuspneumoniae | Penicillins | Augmentin(amoxicillin/clavulan ate) | 0 | 78 | 78 | 120 |
| Streptococcuspneumoniae | Penicillins | Unasyn(ampicillin/sulbactam) | 1 | 33 | 34 | 309 |
| Streptococcuspneumoniae | Penicillins | Zosyn(pipercillin/tazobactam) | 1 | 33 | 34 | 309 |
| Streptococcuspneumoniae | Carbapenems | Doripenem | 0 | 304 | 304 | 1 |
| Streptococcuspneumoniae | Carbapenems | Imipenem | 0 | 150 | 150 | 310 |
| Streptococcuspneumoniae | Carbapenems | Meropenem | 81 | 277 | 358 | 311 |
| Streptococcuspneumoniae | Carbapenems | Meropenem | 3 | 147 | 150 | 310 |
| Streptococcuspneumoniae | Carbapenems | Meropenem | 141 | 587 | 728 | 312 |
| Streptococcuspneumoniae | Cephalosporins(2ndgen ) | Cefoxitin | 3 | 33 | 36 | 170 |
| Streptococcuspneumoniae | Cephalosporins(2ndgen ) | Cefuroxime | 110 | 248 | 358 | 311 |
| Streptococcuspneumoniae | Cephalosporins(2ndgen ) | Cefuroxime | 86 | 93 | 179 | 118 |
| Streptococcuspneumoniae | Cephalosporins(2ndgen ) | Cefuroxime | 11 | 67 | 78 | 120 |
| Streptococcuspneumoniae | Cephalosporins(2ndgen ) | Cefaclor | 114 | 65 | 179 | 118 |
| Streptococcuspneumoniae | Cephalosporins(2ndgen ) | Cefaclor | 194 | 66 | 260 | 119 |

| | | | | | | |
|---|---|---|---|---|---|---|
| Streptococcuspneumoniae | Cephalosporins(2ndgen) | Cefaclor | 11 | 67 | 78 | 120 |
| Streptococcuspneumoniae | Cephalosporins(3rdgen) | Cefotaxime | 89 | 639 | 728 | 312 |
| Streptococcuspneumoniae | Cephalosporins(3rdgen) | Cefotaxime | 191 | 1664 | 1855 | 313 |
| Streptococcuspneumoniae | Cephalosporins(3rdgen) | Cefotaxime | 0 | 170 | 170 | 314 |
| Streptococcuspneumoniae | Cephalosporins(3rdgen) | Cefdinir | 77 | 65 | 142 | 315 |
| Streptococcuspneumoniae | Cephalosporins(3rdgen) | Cefdinir | 5 | 89 | 94 | 129 |
| Streptococcuspneumoniae | Cephalosporins(3rdgen) | Cefdinir | 13 | 65 | 78 | 120 |
| Streptococcuspneumoniae | Cephalosporins(3rdgen) | Ceftriaxone | 3 | 176 | 179 | 118 |
| Streptococcuspneumoniae | Cephalosporins(3rdgen) | Ceftriaxone | 21 | 239 | 260 | 119 |
| Streptococcuspneumoniae | Cephalosporins(3rdgen) | Ceftriaxone | 0 | 78 | 78 | 120 |
| Streptococcuspneumoniae | Cephalosporins(3rdgen) | Cefpodoxime | 3 | 91 | 94 | 129 |
| Streptococcuspneumoniae | Cephalosporins(3rdgen) | Cefpodoxime | 28 | 140 | 168 | 130 |
| Streptococcuspneumoniae | Cephalosporins(3rdgen) | Cefpodoxime | 37 | 102 | 139 | 131 |
| Streptococcuspneumoniae | Cephalosporins(4thgen) | Cefepime | 29 | 121 | 150 | 310 |
| Streptococcuspneumoniae | Cephalosporins(4thgen) | Cefepime | 16 | 26 | 42 | 316 |
| Streptococcuspneumoniae | Macrolides | Erythromycin | 160 | 60 | 220 | 317 |
| Streptococcuspneumoniae | Macrolides | Erythromycin | 32 | 12 | 44 | 305 |

| | | | | | | |
|---|---|---|---|---|---|---|
| Streptococcuspneumoniae | Macrolides | Erythromycin | 86 | 93 | 179 | 118 |
| Streptococcuspneumoniae | Macrolides | Clarithromycin | 86 | 93 | 179 | 118 |
| Streptococcuspneumoniae | Macrolides | Clarithromycin | 200 | 60 | 260 | 119 |
| Streptococcuspneumoniae | Macrolides | Clarithromycin | 16 | 62 | 78 | 120 |
| Streptococcuspneumoniae | Macrolides | Azithromycin | 184 | 174 | 358 | 311 |
| Streptococcuspneumoniae | Macrolides | Azithromycin | 212 | 378 | 590 | 135 |
| Streptococcuspneumoniae | Macrolides | Azithromycin | 68 | 7 | 75 | 318 |
| Streptococcuspneumoniae | Macrolides | Clindamycin | 19 | 25 | 44 | 305 |
| Streptococcuspneumoniae | Macrolides | Clindamycin | 61 | 14 | 75 | 318 |
| Streptococcuspneumoniae | Macrolides | Clindamycin | 170 | 239 | 409 | 308 |
| Streptococcuspneumoniae | Macrolides | Pristinamycin(Streptogramin) | 0 | 51 | 51 | 319 |
| Streptococcuspyogenes | Penicillins | Penicillin | 0 | 29 | 29 | 123 |
| Streptococcuspyogenes | Penicillins | Penicillin | 0 | 63 | 63 | 320 |
| Streptococcuspyogenes | Penicillins | Penicillin | 0 | 38 | 38 | 321 |
| Streptococcuspyogenes | Penicillins | Amoxicillin | 1 | 34 | 35 | 322 |
| Streptococcuspyogenes | Penicillins | Ampicillin | 0 | 16 | 16 | 309 |
| Streptococcuspyogenes | Penicillins | Ampicillin | 0 | 29 | 29 | 123 |
| Streptococcuspyogenes | Penicillins | Ampicillin | 0 | 16 | 16 | 172 |
| Streptococcuspyogenes | Penicillins | Augmentin(amoxicillin/clavulanate) | 0 | 222 | 222 | 174 |
| Streptococcuspyogenes | Penicillins | Augmentin(amoxicillin/clavulanate) | 0 | 125 | 125 | 323 |

| | | | | | | |
|---|---|---|---|---|---|---|
| Streptococcuspyogenes | Penicillins | Unasyn(ampicillin/sulbactam) | 0 | 16 | 16 | 309 |
| Streptococcuspyogenes | Penicillins | Zosyn(pipercillin/tazobactam) | 0 | 16 | 16 | 309 |
| Streptococcuspyogenes | Cephalosporins(2ndgen ) | Cefuroxime | 0 | 222 | 222 | 174 |
| Streptococcuspyogenes | Cephalosporins(2ndgen ) | Cefuroxime | 0 | 125 | 125 | 323 |
| Streptococcuspyogenes | Cephalosporins(2ndgen ) | Cefuroxime | 0 | 78 | 78 | 324 |
| Streptococcuspyogenes | Cephalosporins(2ndgen ) | Cefaclor | 0 | 222 | 222 | 174 |
| Streptococcuspyogenes | Cephalosporins(2ndgen ) | Cefaclor | 0 | 78 | 78 | 324 |
| Streptococcuspyogenes | Cephalosporins(3rdgen ) | Cefotaxime | 0 | 54 | 54 | 320 |
| Streptococcuspyogenes | Cephalosporins(3rdgen ) | Cefotaxime | 2 | 46 | 48 | 321 |
| Streptococcuspyogenes | Cephalosporins(3rdgen ) | Cefotaxime | 0 | 140 | 140 | 325 |
| Streptococcuspyogenes | Cephalosporins(3rdgen ) | Ceftriaxone | 1 | 34 | 35 | 322 |
| Streptococcuspyogenes | Cephalosporins(3rdgen ) | Ceftriaxone | 0 | 36 | 36 | 320 |
| Streptococcuspyogenes | Cephalosporins(3rdgen ) | Ceftriaxone | 2 | 34 | 36 | 321 |
| Streptococcuspyogenes | Cephalosporins(5thgen ) | Ceftaroline | 0 | 193 | 193 | 326 |
| Streptococcuspyogenes | Cephalosporins(5thgen ) | Ceftaroline | 0 | 174 | 174 | 98 |
| Streptococcuspyogenes | Cephalosporins(5thgen ) | Ceftaroline | 0 | 312 | 312 | 98 |
| Streptococcuspyogenes | Macrolides | Erythromycin | 15 | 256 | 271 | 327 |

| | | | | | | |
|---|---|---|---|---|---|---|
| Streptococcuspyogenes | Macrolides | Erythromycin | 1 | 34 | 35 | 322 |
| Streptococcuspyogenes | Macrolides | Erythromycin | 7 | 56 | 63 | 328 |
| Streptococcuspyogenes | Macrolides | Azithromycin | 20 | 123 | 143 | 329 |
| Streptococcuspyogenes | Macrolides | Clindamycin | 14 | 257 | 271 | 327 |
| Streptococcuspyogenes | Macrolides | Clindamycin | 1 | 102 | 103 | 330 |
| Streptococcuspyogenes | Macrolides | Clindamycin | 1 | 31 | 32 | 328 |
| Streptococcusviridans | Penicillins | Penicillin | 0 | 216 | 216 | 133 |
| Streptococcusviridans | Penicillins | Penicillin | 0 | 10 | 10 | 331 |
| Streptococcusviridans | Penicillins | Penicillin | 4 | 39 | 43 | 332 |
| Streptococcusviridans | Penicillins | Oxacillin | 0 | 10 | 10 | 331 |
| Streptococcusviridans | Penicillins | Amoxicillin | 1 | 634 | 635 | 333 |
| Streptococcusviridans | Penicillins | Ampicillin | 35 | 600 | 635 | 333 |
| Streptococcusviridans | Penicillins | Ampicillin | 10 | 769 | 779 | 334 |
| Streptococcusviridans | Penicillins | Augmentin(amoxicillin/clavulanate) | 2 | 72 | 74 | 335 |
| Treponemapallidum | Macrolides | Clarithromycin | 94 | 38 | 132 | 336 |
| Treponemapallidum | Macrolides | Azithromycin | 74 | 280 | 354 | 337 |
| Treponemapallidum | Macrolides | Azithromycin | 162 | 16 | 178 | 338 |
| Ureaplasmaurealyticum | Fluoroquinolones | Ciprofloxacin | 3472 | 1226 | 4698 | 339 |
| Ureaplasmaurealyticum | Fluoroquinolones | Ciprofloxacin | 54 | 37 | 91 | 5 |
| Ureaplasmaurealyticum | Fluoroquinolones | Ciprofloxacin | 33 | 15 | 48 | 340 |

| | | | | | | |
|---|---|---|---|---|---|---|
| Ureaplasmaurealyticum | Fluoroquinolones | Levofloxacin | 61 | 13 | 74 | 236 |
| Ureaplasmaurealyticum | Fluoroquinolones | Levofloxacin | 257 | 4441 | 4698 | 339 |
| Ureaplasmaurealyticum | Fluoroquinolones | Levofloxacin | 0 | 13 | 13 | 341 |
| Ureaplasmaurealyticum | Fluoroquinolones | Moxifloxacin | 46 | 28 | 74 | 236 |
| Ureaplasmaurealyticum | Fluoroquinolones | Norfloxacin | 577 | 195 | 772 | 342 |
| Ureaplasmaurealyticum | Fluoroquinolones | Ofloxacin | 192 | 4506 | 4698 | 339 |
| Ureaplasmaurealyticum | Fluoroquinolones | Ofloxacin | 18 | 19 | 37 | 340 |
| Ureaplasmaurealyticum | Fluoroquinolones | Ofloxacin | 12 | 79 | 91 | 5 |
| Ureaplasmaurealyticum | Macrolides | Erythromycin | 4 | 70 | 74 | 236 |
| Ureaplasmaurealyticum | Macrolides | Erythromycin | 671 | 4027 | 4698 | 339 |
| Ureaplasmaurealyticum | Macrolides | Erythromycin | 14 | 458 | 472 | 343 |
| Ureaplasmaurealyticum | Macrolides | Clarithromycin | 125 | 4573 | 4698 | 339 |
| Ureaplasmaurealyticum | Macrolides | Clarithromycin | 0 | 45 | 45 | 344 |
| Ureaplasmaurealyticum | Macrolides | Clarithromycin | 4 | 468 | 472 | 343 |
| Ureaplasmaurealyticum | Macrolides | Clarithromycin | 35 | 2 | 37 | 340 |
| Ureaplasmaurealyticum | Macrolides | Azithromycin | 136 | 4562 | 4698 | 339 |
| Ureaplasmaurealyticum | Macrolides | Azithromycin | 0 | 45 | 45 | 344 |
| Ureaplasmaurealyticum | Macrolides | Azithromycin | 72 | 26 | 98 | 345 |
| Ureaplasmaurealyticum | Macrolides | Clindamycin | 235 | 82 | 317 | 346 |
| Ureaplasmaurealyticum | Macrolides | Pristinamycin(Streptogramin) | 0 | 15 | 15 | 347 |

| | | | | | | |
|---|---|---|---|---|---|---|
| Ureaplasmaurealyticum | Macrolides | Pristinamycin(Streptogramin) | 19 | 18 | 37 | 340 |
| Ureaplasmaurealyticum | Tetracyclines | Doxycyline | 0 | 15 | 15 | 347 |
| Ureaplasmaurealyticum | Tetracyclines | Doxycyline | 13 | 85 | 98 | 345 |
| Ureaplasmaurealyticum | Tetracyclines | Doxycyline | 13 | 759 | 772 | 342 |
| Ureaplasmaurealyticum | Tetracyclines | Minocycline | 184 | 4514 | 4698 | 339 |
| Ureaplasmaurealyticum | Tetracyclines | Minocycline | 12 | 86 | 98 | 345 |
| Ureaplasmaurealyticum | Tetracyclines | Minocycline | 15 | 757 | 772 | 342 |
| Ureaplasmaurealyticum | Tetracyclines | Tetracycline | 12 | 3 | 15 | 347 |
| Ureaplasmaurealyticum | Tetracyclines | Tetracycline | 0 | 13 | 13 | 341 |
| Ureaplasmaurealyticum | Tetracyclines | Tetracycline | 0 | 48 | 48 | 348 |
| Vibriocholerae | Tetracyclines | Doxycyline | 63 | 8 | 71 | 349 |
| Vibriocholerae | Tetracyclines | Doxycyline | 88 | 70 | 158 | 350 |
| Vibriocholerae | Tetracyclines | Minocycline | 9 | 11 | 20 | 351 |
| Vibriocholerae | Tetracyclines | Oxytetracycline | 264 | 316 | 580 | 352 |
| Vibriocholerae | Tetracyclines | Tetracycline | 1 | 55 | 56 | 279 |
| Vibriocholerae | Tetracyclines | Tetracycline | 917 | 1467 | 2384 | 352 |
| Vibriocholerae | Tetracyclines | Tetracycline | 65 | 93 | 158 | 349 |

40. Praharaj, A. K., Khajuria, A., Kumar, M. & Grover, N. Phenotypic detection and molecular characterization of beta-lactamase genes among

Citrobacter species in a tertiary care hospital. *Avicenna J. Med.* **06**, 17–27 (2016).

105. Plantamura, J. *et al.* Molecular epidemiological of extended-spectrum β-lactamase producing Escherichia coli isolated in Djibouti. *J. Infect.*

*Dev. Ctries.* **13**, 753–758 (2019).

180. Zhao, F. *et al.* Antimicrobial susceptibility and molecular characteristics of Mycoplasma pneumoniae isolates across different regions of

China. *Antimicrob. Resist. Infect. Control* **8**, 1–8 (2019).

*Agents* **51**, 768–774 (2018).

234. Ahmed, N. *et al.* Evaluation of antibiotic resistance and virulence genes among clinical isolates of Pseudomonas aeruginosa from cancer

patients. *Asian Pacific J. Cancer Prev.* **21**, 1333–1338 (2020).

261. Khatun, H. *et al.* Clinical profile, antibiotic susceptibility pattern of bacterial isolates and factors associated with complications in culture-

proven typhoid patients admitted to an urban hospital in Bangladesh. *Trop. Med. Int. Heal.* **23**, 359–366 (2018).

301. Bastidas, C. A. *et al.* Antibiotic susceptibility profile and prevalence of mecA and lukS-PV/lukF-PV genes in Staphylococcus aureus isolated

from nasal and pharyngeal sources of medical students in Ecuador. *Infect. Drug Resist.* **12**, 2553–2560 (2019).

#### Supplemental Table 5

| Pathogen | Reservoir | Transmission Mode* | Nosocomial (Yes-1 or No-0) | Zoonotic (Yes-1 or No-0) | Human to Human transmission (Yes-1 or No-0) | Commensals (Yes-1 or No-0) | Conjugation (Yes-1 or No-0) | Naturally Competent (Yes-1 or No-0) | Citations |
|---|---|---|---|---|---|---|---|---|---|
| *Acinetobacter* spp | Environment | Vehicle-Borne | 1 | No | 1 | Yes | 1 | Yes | 1–4 |
| *Actinomyces spp* | Human | Direct Contact | 0 | No | 0 | Yes | 0 | No | 4–6 |
| *Bacillus anthracis* | Animal | Direct Contact | 0 | Yes | 0 | No | 1 | No | 4,7–10 |
| *Bacteroides spp* | Human | Vehicle-Borne | 1 | No | 0 | Yes | 1 | No | 4,11–14 |
| *Bordetella pertussis* | Human | Droplet | 1 | No | 1 | No | 1 | No | 4,15–17 |
| *Borrelia burgdorferi* | Animal | Vector-Borne | 0 | Yes | 0 | No | 0 | No | 4,18,19 |
| *Brucella spp* | Animal | Vehicle-Borne | 0 | Yes | 0 | No | 1 | No | 4,20–22 |
| *Campylobacter jejuni* | Animal | Vehicle-Borne | 0 | Yes | 0 | No | 1 | Yes | 4,23–26 |
| *Chlamydia pneumoniae* | Human | Droplet | 1 | No | 1 | No | 0 | No | 4,27–29 |
| *Chlamydia psittaci* | Animal | Airborne | 0 | Yes | 0 | No | 0 | No | 4,30–32 |
| *Chlamydia trachomatis* | Human | Direct Contact | 0 | No | 1 | No | 0 | No | 4,33,34 |

| | | | | | | | | | |
|---|---|---|---|---|---|---|---|---|---|
| *Citrobacter* spp | Human | Vehicle-Borne | 1 | No | 0 | Yes | 1 | No | 4,35–38 |
| *Clostridium difficile* | Environment | Vehicle-Borne | 1 | No | 1 | Yes | 1 | No | 4,39–41 |
| *Clostridium perfringens* | Environment | Vehicle-Borne | 0 | No | 0 | Yes | 1 | No | 4,42,43 |
| *Clostridium spp* | Environment | Direct Contact (excluding C. botulinum) | 0 | No | 0 | Yes | 1 | No | 4,44–46 |
| *Clostridium tetani* | Environment | Direct Contact | 0 | No | 0 | No | 0 | No | 4,47 |
| *Corynebacterium diptheriae* | Human | Droplet | 0 | No | 1 | No | 1 | No | 4,48,49 |
| *Enterobacter aerogenes* | Human | Vehicle-Borne | 1 | No | 1 | Yes | 1 | No | 4,50–53 |
| *Enterococcus faecalis* | Human | Vehicle-Borne | 1 | Yes | 1 | Yes | 1 | No | 4,54–57 |
| Escherichia coli (ETEC) | Animal | Vehicle-Borne | 1 | Yes | 1 | Yes | 1 | Yes | 4,58–60 |
| *Francisella tularensis* | Animal | Vehicle-Borne | 0 | Yes | 0 | No | 0 | No | 4,61–63 |
| *Fuseobacterium spp* | Human | Direct Contact | 1 | No | 0 | Yes | 1 | No | 4,64–67 |

| | | | | | | | | | |
|---|---|---|---|---|---|---|---|---|---|
| *Gardnerella vaginalis* | Human | Direct Contact | 0 | No | 1 | Yes | 0 | No | 4,68–73 |
| *GPAC* | Human | Vehicle-Borne (Fecal-Oral) | 0 | No | 1 | Yes | 0 | No | 4,74 |
| *Haemophilus influenzae* | Human | Droplet | 1 | No | 1 | Yes | 1 | Yes | 4,75–78 |
| *Klebsiella oxytoca* | Human | Direct Contact | 1 | No | 1 | Yes | 1 | No | 4,64,79–81 |
| *Klebsiella pneumoniae* | Human | Direct Contact | 1 | No | 1 | Yes | 1 | No | 4,59,82,83 |
| *Klebsiella spp* | Human | Direct Contact | 1 | No | 1 | Yes | 1 | No | 4,80,84,85 |
| *Legionella pneumophila* | Environment | Airborne | 1 | No | 0 | No | 1 | Yes | 4,30,86,87 |
| *Leptospira interrogans* | Animal | Direct Contact | 0 | Yes | 0 | No | 1 | No | 4,88–90 |
| *Listeria monocytogenes* | Environment | Vehicle-Borne | 0 | No | 1 | No | 1 | No | 4,91–93 |
| *Moraxella catarrhalis* | Human | Direct Contact | 1 | No | 0 | Yes | 1 | Yes | 4,94–97 |
| *Mycoplasma pneumoniae* | Human | Droplet | 0 | No | 1 | No | 0 | No | 4,98,99 |

| | | | | | | | | | |
|---|---|---|---|---|---|---|---|---|---|
| *Neisseria gonorrhoeae* | Human | Direct Contact | 0 | No | 1 | No | 1 | Yes | 4,100–103 |
| *Neisseria meningitidis* | Human | Direct | 0 | No | 1 | Yes | 1 | Yes | 4,104–106 |
| *Nocardia spp* | Environment | Airborne | 0 | No | 0 | No | 1 | No | 4,107–109 |
| *Non-typhoidal Salmonella* | Animal | Vehicle-Borne | 0 | Yes | 1 | No | 1 | No | 4,110–112 |
| *Propiniobacterium acnes* | Human | Vehicle-Borne | 1 | No | 0 | Yes | 1 | No | 4,113–115 |
| *Proteus mirabilis* | Human | Vehicle-Borne | 1 | No | 0 | Yes | 1 | No | 4,116–118 |
| *Proteus spp* | Human | Vehicle-Borne | 1 | No | 0 | Yes | 1 | No | 4,119–121 |
| *Providencia spp* | Human | Vehicle-Borne | 1 | No | 0 | Yes | 1 | No | 4,122–125 |
| *Pseudomonas aeruginosa* | Environment | Vehicle-Borne | 1 | No | 1 | Yes | 1 | Yes | 4,126–128 |
| *Rickettsia rickettsii* | Animal | Vector-Borne | 0 | Yes | 0 | No | 0 | No | 4,129,130 |
| *Salmonella typhi* | Human | Vehicle-Borne | 0 | No | 1 | No | 1 | No | 4,93,131,132 |
| *Serratia marcescens* | Environment | Direct Contact | 1 | No | 0 | No | 1 | No | 4,133–136 |
| *Shigella* species | Human | Vehicle-Borne | 0 | No | 1 | No | 1 | No | 4,137,138 |
| *Staphylococcus aureus* | Human | Direct Contact | 1 | No | 1 | Yes | 1 | Yes | 4,139–141 |
| *Staphylococcus epidermidis* | Human | Vehicle-Borne | 1 | No | 0 | Yes | 1 | No | 4,142,143 |

| | | | | | | | | | |
|---|---|---|---|---|---|---|---|---|---|
| *Streptococcus agalactiae* | Human | Direct Contact | 1 | No | 0 | Yes | 1 | No | 4,144–146 |
| *Streptococcus pneumoniae* | Human | Droplet | 1 | No | 1 | Yes | 1 | Yes | 4,147,148 |
| *Streptococcus pyogenes* | Human | Droplet | 1 | No | 1 | Yes | 1 | No | 4,149,150 |
| *Streptococcus viridans* | Human | Direct | 1 | No | 0 | Yes | 1 | No | 4,151–153 |
| *Treponema pallidum* | Human | Direct Contact | 0 | No | 1 | No | 0 | No | 4,154 |
| *Treponema pallidum pertenue* | Human | Direct Contact | 0 | No | 1 | No | 0 | No | 4,155 |
| *Ureaplasma urealyticum* | Human | Direct Contact | 0 | No | 1 | Yes | 1 | No | 4,156,157 |
| *Vibrio cholerae* | Environment | Vehicle-Borne | 0 | No | 1 | No | 1 | Yes | 4,158–160 |
| *Yersinia pestis* | Animal | Vector-Borne | 0 | Yes | 0 | No | 1 | No | 4,135,161 |

*Transmission modes were analysed as Direct or Indirect. Direct modes of transmission include 'Direct Contact' whereas Indirect transmission includes 'Vector-Borne', 'Vehicle-Borne' , 'Droplet', and 'Air-borne'.

1. Atrouni, A. Al, Joly-Guillou, M. L., Hamze, M. & Kempf, M. Reservoirs of non-baumannii Acinetobacter species. *Front. Microbiol.* **7**, 1–12 (2016).
2. Leungtongkam, U., Thummeepak, R., Tasanapak, K. & Sitthisak, S. Acquisition and transfer of antibiotic resistance genes in association with conjugative plasmid or class 1 integrons of Acinetobacter baumannii. *PLoS One* **13**, 1–12 (2018).
3. Venanzio, G. Di *et al.* Multidrug-resistant plasmids repress chromosomally encoded T6SS to enable their dissemination. *Proc. Natl. Acad. Sci. U. S. A.* **116**, 1378–1383 (2019).
4. Johnsborg, O., Eldholm, V. & Håvarstein, L. S. Natural genetic transformation: prevalence, mechanisms and function. *Res. Microbiol.* **158**, 767–778 (2007).
5. Könönen, E. & Wade, W. G. Actinomyces and related organisms in human infections. *Clin. Microbiol. Rev.* **28**, 419–442 (2015).
6. Bowden, G. H. W. *Actinomyces, Propionibacterium propionicus, and Streptomyces*. *Medical Microbiology* (University of Texas Medical Branch at Galveston, 1996).
7. What is Anthrax? | CDC. *Centers for Disease Control and Prevention, National Center for Emerging and Zoonotic Infectious Diseases (NCEZID)* https://www.cdc.gov/anthrax/basics/index.html (2020).
8. Koehler, T. M. Bacillus anthracis Physiology and Genetics. *Mol. Aspects Med.* **30**, 386–396 (2009).
9. Saile, E. & Koehler, T. M. Bacillus anthracis multiplication, persistence, and genetic exchange in the rhizosphere of grass plants. *Appl. Environ. Microbiol.* **72**, 3168–3174 (2006).
10. Yuan, Y., Zheng, D., Hu, X., Cai, Q. & Yuan, Z. Conjugative Transfer of Insecticidal Plasmid pHT73 from Bacillus thuringiensis to B. anthracis and Compatibility of This Plasmid with pXO1 and pXO2. *Appl. Environ. Microbiol.* **76**, 468–473 (2010).
11. Patrick, S. Bacteroides. in *Molecular Medical microbiology* 917–944 (Elsevier, 2014).
12. Shoemaker, N. B., Vlamakis, H., Hayes, K. & Salyers, A. A. Evidence for extensive resistance gene transfer among Bacteroides spp. and among Bacteroides and other genera in the human colon. *Appl. Environ. Microbiol.* **67**, 561–568 (2001).
13. Nguyen, M. & Vedantam, G. Mobile genetic elements in the genus Bacteroides, and their mechanism(s) of dissemination. *Mob. Genet. Elements* **1**, 187–196 (2011).
14. Pathogen Safety Data Sheets: Infectious Substances – Bacteroides spp. - Canada.ca. https://www.canada.ca/en/public-health/services/laboratory-biosafety-biosecurity/pathogen-safety-data-sheets-risk-assessment/bacteroides.html.
15. Weiss, A. A. & Falkow, S. Genetic analysis of phase change in Bordetella pertussis. *Infect. Immun.* **43**, 263–269 (1984).

16. Weiss, A. A. & Falkow, S. Plasmid transfer to Bordetella pertussis: Conjugation and transformation. *J. Bacteriol.* **152**, 549–552 (1982).

17. OCHMAN, H. Evolution of Bacterial Pathogens. *Princ. Bact. Pathog.* 1–41 (2001) doi:10.1016/B978-012304220-0/50002-9.

18. Transmission | Lyme Disease | CDC. https://www.cdc.gov/lyme/transmission/index.html.

19. Brisson, D., Drecktrah, D., Eggers, C. H. & Samuels, D. S. Genetics of Borrelia burgdorferi. *Annu. Rev. Genet.* **46**, 515–536 (2012).

20. Verger, J. M., Grayon, M., Chaslus-Dancla, E., Meurisse, M. & Lafont, J. P. Conjugative Transfer and in Vitro/in Vivo Stability of the Broad-Host-Range IncP R751 Plasmid in Brucella spp. *Plasmid* **29**, 142–146 (1992).

21. Wattam, A. R. *et al.* Analysis of ten Brucella genomes reveals evidence for horizontal gene transfer despite a preferred intracellular lifestyle. *J. Bacteriol.* **191**, 3569–3579 (2009).

22. Transmission | Brucellosis | CDC. https://www.cdc.gov/brucellosis/transmission/index.html.

23. Questions and Answers | Campylobacter | CDC. https://www.cdc.gov/campylobacter/faq.html.

24. Wilson, D. J. *et al.* Tracing the source of campylobacteriosis. *PLoS Genet.* **4**, (2008).

25. Zeng, X., Wu, Z., Zhang, Q. & Lin, J. A cotransformation method to identify a restriction-modification enzyme that reduces conjugation efficiency in Campylobacter jejuni. *Appl. Environ. Microbiol.* **84**, 1–13 (2018).

26. Avrain, L., Vernozy-Rozand, C. & Kempf, I. Evidence for natural horizontal transfer of tetO gene between Campylobacter jejuni strains in chickens. *J. Appl. Microbiol.* **97**, 134–140 (2004).

27. Chlamydia pneumoniae: Causes, How It Spreads, and Risk Factors | CDC. https://www.cdc.gov/pneumonia/atypical/cpneumoniae/about/causes.html.

28. Sixt, B. S. & Valdivia, R. H. Molecular Genetic Analysis of Chlamydia Species. *Annu. Rev. Microbiol.* **70**, 179–198 (2016).

29. M, R. & KA, F. Transformation of Chlamydia: current approaches and impact on our understanding of chlamydial infection biology. *Microbes Infect.* **20**, 445–450 (2018).

30. González-Rivera, E. M. *et al.* Antibiotic resistance, virulence factors and genotyping of pseudomonas aeruginosa in public hospitals of northeastern mexico. *J. Infect. Dev. Ctries.* **13**, 374–383 (2019).

31. Hooppaw, A. J. & Fisher, D. J. A coming of age story: Chlamydia in the post-genetic era. *Infect. Immun.* **84**, 612–621 (2016).

32. Psittacosis: Clinical Disease Specifics | CDC. https://www.cdc.gov/pneumonia/atypical/psittacosis/hcp/disease-specifics.html.

33. Detailed STD Facts - Chlamydia. https://www.cdc.gov/std/chlamydia/stdfact-chlamydia-detailed.htm.

41. What is C. diff? | CDC. https://www.cdc.gov/cdiff/what-is.html#factsheet.

42. C. perfringens | CDC. https://www.cdc.gov/foodsafety/diseases/clostridium-perfringens.html.

43. Wisniewski, J. A. & Rood, J. I. The Tcp conjugation system of Clostridium perfringens. *Plasmid* **91**, 28–36 (2017).

44. Pathogen Safety Data Sheets: Infectious Substances – Clostridium spp. - Canada.ca. https://www.canada.ca/en/public-health/services/laboratory-biosafety-biosecurity/pathogen-safety-data-sheets-risk-assessment/clostridium.html.

45. Philipps, G., De Vries, S. & Jennewein, S. Development of a metabolic pathway transfer and genomic integration system for the syngas-fermenting bacterium Clostridium ljungdahlii. *Biotechnol. Biofuels* **12**, 1–14 (2019).

46. Vidor, C. J. *et al.* Clostridium sordellii Pathogenicity Locus Plasmid pCS1-1 Encodes a Novel Clostridial Conjugation Locus. *Am. Soc. Microbiol.* **9**, 1–14 (2018).

47. Tetanus Causes and Transmission | CDC. https://www.cdc.gov/tetanus/about/causes-transmission.html.

48. Diphtheria: Causes and Spread to Others | CDC. https://www.cdc.gov/diphtheria/about/causes-transmission.html.

49. Hennart, M. *et al.* Population genomics and antimicrobial resistance in Corynebacterium diphtheriae. *Genome Med.* **12**, 1–18 (2020).

50. Pathogen Safety Data Sheets: Infectious Substances – Enterobacter spp. - Canada.ca. https://www.canada.ca/en/public-health/services/laboratory-biosafety-biosecurity/pathogen-safety-data-sheets-risk-assessment/enterobacter.html.

51. Davin-Regli, A. *et al.* Molecular epidemiology of Enterobacter aerogenes acquisition: One-year prospective study in two intensive care units. *J. Clin. Microbiol.* **34**, 1474–1480 (1996).

52. Burmølle, M., Bahl, M. I., Jensen, L. B., Sørensen, S. J. & Hansen, L. H. Type 3 fimbriae, encoded by the conjugative plasmid pOLA52, enhance biofilm formation and transfer frequencies in Enterobacteriaceae strains. *Microbiology* **154**, 187–195 (2008).

53. Chavda, K. D. *et al.* Comprehensive genome analysis of carbapenemase-producing Enterobacter spp.: New insights into phylogeny, population structure, and resistance mechanisms. *MBio* **7**, 1–16 (2016).

54. Audrey Wanger, V. C. et al. Chapter 6. Overview of Bacteria | Elsevier Enhanced Reader. in *Microbiology and Molecular Diagnosis in Pathology* 97–98 (2017).

55. Pathogen Safety Data Sheets: Infectious Substances – Enterococcus faecalis and Enterococcus faecium - Canada.ca. https://www.canada.ca/en/public-health/services/laboratory-biosafety-biosecurity/pathogen-safety-data-sheets-risk-assessment/enterococcus-faecalis.html.

56. Frost, S. Bacterial conjugation : everybody ' s doin ' it. *Can. J. Microbiol.* 1091–1096 (1992).

57. Hirt, H. *et al.* Enterococcus faecalis sex pheromone cCF10 enhances conjugative plasmid transfer in vivo. *MBio* **9**, (2018).

58. Enterotoxigenic E. coli (ETEC) | E. coli | CDC. https://www.cdc.gov/ecoli/etec.html.

59. Lerminiaux, N. A. & Cameron, A. D. S. Horizontal transfer of antibiotic resistance genes in clinical environments. *Can. J. Microbiol.* **65**, 34–44 (2019).

60. Murray, B. E., Evans, D. J., Penaranda, M. E. & Evans, D. G. CFA/I-ST plasmids: Comparison of enterotoxigenic Escherichia coli (ETEC) of serogroups O25, O63, O78, and O128 and mobilization from an R factor-containing epidemic ETEC isolate. *J. Bacteriol.* **153**, 566–570 (1983).

61. Transmission | Tularemia | CDC. https://www.cdc.gov/tularemia/transmission/index.html.

62. World Health Organization. *WHO guidelines on tularaemia: epidemic and pandemic alert and response*. http://www.who.int/csr/resources/publications/WHO_CDS_EPR_2007_7.pdf (2007).

63. Siddaramappa, S., Challacombe, J. F., Petersen, J. M., Pillai, S. & Kuske, C. R. Comparative analyses of a putative Francisella conjugative

element. *Genome* **57**, 137–144 (2014).

64. Garrett, W. S. & Onderdonk, A. B. Bacteroides, Prevotella, Porphyromonas, and Fusobacterium Species (and Other Medically Important Anaerobic Gram-Negative Bacilli). in *Mandell, Douglas, and Bennett's Principles and Practice of Infectious Diseases* vol. 2 2773–2780 (2014).

65. Roberts, M. C. & Lansciardi, J. Transferable Tet M in Fusobacterium nucleatum. *Antimicrob. Agents Chemother.* **34**, 1836–1838 (1990).

66. Claypool, B. M. *et al.* Mobilization and prevalence of a fusobacterial plasmid. *Plasmid* **63**, 11–19 (2010).

67. Riordan, T. Human infection with Fusobacterium necrophorum (Necrobacillosis), with a focus on Lemierre's syndrome. *Clin. Microbiol. Rev.* **20**, 622–659 (2007).

68. STD Facts - Bacterial Vaginosis. https://www.cdc.gov/std/bv/stdfact-bacterial-vaginosis.htm.

76. Haemophilus influenzae: Causes and Transmission | CDC. https://www.cdc.gov/hi-disease/about/causes-transmission.html.

77. Hegstad, K. *et al.* Role of Horizontal Gene Transfer in the Development of Multidrug Resistance in Haemophilus influenzae. *mSphere* **5**, (2020).

78. Stuy, J. H. Chromosomally integrated conjugative plasmids are common in antibiotic-resistant Haemophilus influenzae. *J. Bacteriol.* **142**,

925–930 (1980).

79. Podschun, R. & Ullmann, U. Klebsiella spp. as nosocomial pathogens: Epidemiology, taxonomy, typing methods, and pathogenicity factors. *Clin. Microbiol. Rev.* **11**, 589–603 (1998).

80. Evans, D. R. *et al.* Systematic detection of horizontal gene transfer across genera among multidrug-resistant bacteria in a single hospital. *Elife* **9**, 1–20 (2020).

81. Yigit, H. *et al.* Carbapenem-Resistant Strain of Klebsiella oxytoca Harboring Carbapenem-Hydrolyzing β-Lactamase KPC-2. *Antimicrob. Agents Chemother.* **47**, 3881–3889 (2003).

82. Klebsiella pneumoniae in Healthcare Settings | HAI | CDC. https://www.cdc.gov/hai/organisms/klebsiella/klebsiella.html.

83. Dixon, R. A. & Postgate, J. R. Transfer of nitrogen-fixation genes by conjugation in Klebsiella pneumoniae [12]. *Nature* vol. 234 47–48 (1971).

84. Samanta, I. & Bandyopadhyay, S. Klebsiella. in *Antimicrobial Resistance in Agriculture* 153–169 (Academic Press, 2020). doi:10.1016/b978-0-12-815770-1.00014-6.

85. Pathogen Safety Data Sheets: Infectious Substances – Klebsiella spp. - Canada.ca. https://www.canada.ca/en/public-health/services/laboratory-biosafety-biosecurity/pathogen-safety-data-sheets-risk-assessment/klebsiella.html.

86. Legionnaires Disease Cause and Spread | CDC. https://www.cdc.gov/legionella/about/causes-transmission.html.

87. Gomez-Valero, L. *et al.* Extensive recombination events and horizontal gene transfer shaped the Legionella pneumophila genomes. *BMC Genomics* **12**, 536 (2011).

88. Infection | Leptospirosis | CDC. https://www.cdc.gov/leptospirosis/infection/index.html.

89. Haake, D. A. *et al.* Molecular Evolution and Mosaicism of Leptospiral Outer Membrane Proteins Involves Horizontal DNA Transfer. *J. Bacteriol.* **186**, 2818–2828 (2004).

90. Picardeau, M. Conjugative transfer between Escherichia coli and Leptospira spp. as a new genetic tool. *Appl. Environ. Microbiol.* **74**, 319–322 (2008).

91. Information for Health Professionals and Laboratories | Listeria | CDC. https://www.cdc.gov/listeria/technical.html.

98. Mycoplasma pneumoniae Causes and Transmission | CDC. https://www.cdc.gov/pneumonia/atypical/mycoplasma/about/causes-transmission.html.

99. Xiao, L. *et al.* Comparative genome analysis of Mycoplasma pneumoniae. *BMC Genomics* **16**, 1–16 (2015).

100. Limeres Posse, J., Diz Dios, P. & Scully, C. Systemic Bacteria Transmissible by Kissing. *Saliva Prot. Transm. Dis.* 29–51 (2017) doi:10.1016/b978-0-12-813681-2.00003-2.

101. STD Facts - Gonorrhea. https://www.cdc.gov/std/gonorrhea/stdfact-gonorrhea.htm.

102. Cehovin, A. & Lewis, S. B. Mobile genetic elements in Neisseria gonorrhoeae: Movement for change. *Pathog. Dis.* **75**, 1–12 (2017).

103. Pachulec, E. & van der Does, C. Conjugative plasmids of Neisseria gonorrhoeae. *PLoS One* **5**, (2010).

104. Meningococcal Disease (Neisseria meningitidis) | Disease Directory | Travelers' Health | CDC. https://wwwnc.cdc.gov/travel/diseases/meningococcal-disease.

105. Brett, M. S. Y. Conjugal transfer of gonococcal βlactamase and conjugative plasmids to neisseria meningitidis. *J. Antimicrob. Chemother.* **24**, 875–879 (1989).

106. Roberts, M. C. & Knapp, J. S. Transfer of β-lactamase plasmids from Neisseria gonorrhoeae to Neisseria meningitidis and commensal Neisseria species by the 25.2-megadalton conjugative plasmid. *Antimicrob. Agents Chemother.* **32**, 1430–1432 (1988).

107. Transmission | Nocardiosis | CDC. https://www.cdc.gov/nocardiosis/transmission/index.html.

108. Pathogen Safety Data Sheets: Infectious Substances – Nocardia spp. - Canada.ca. https://www.canada.ca/en/public-

health/services/laboratory-biosafety-biosecurity/pathogen-safety-data-sheets-risk-assessment/nocardia.html.

109. Jung, C. M., Crocker, F. H., Eberly, J. O. & Indest, K. J. Horizontal gene transfer (HGT) as a mechanism of disseminating RDX-degrading activity among Actinomycete bacteria. *J. Appl. Microbiol.* **110**, 1449–1459 (2011).

110. Salmonella (non-typhoidal). https://www.who.int/news-room/fact-sheets/detail/salmonella-%28non-typhoidal%29.

119. Pathogen Safety Data Sheets: Infectious Substances – Proteus spp. - Canada.ca. https://www.canada.ca/en/public-health/services/laboratory-biosafety-biosecurity/pathogen-safety-data-sheets-risk-assessment/proteus.html.

120. Girlich, D., Bonnin, R. A., Dortet, L. & Naas, T. Genetics of Acquired Antibiotic Resistance Genes in Proteus spp. *Front. Microbiol.* **11**, 1–21 (2020).

121. Li, X. *et al.* SXT/R391 integrative and conjugative elements in Proteus species reveal abundant genetic diversity and multidrug resistance. *Sci. Rep.* **6**, 4–12 (2016).

122. Providencia species - Infectious Disease and Antimicrobial Agents. http://antimicrobe.org/b227.asp.

123. Wie, S. H. Clinical significance of providencia bacteremia or bacteriuria. *Korean J. Intern. Med.* **30**, 167–169 (2015).

124. Mahrouki, S. *et al.* Nosocomial dissemination of plasmids carrying blaTEM-24, blaDHA-1, aac(6')-Ib-cr, and qnrA6 in Providencia spp. strains isolated from a Tunisian hospital. *Diagn. Microbiol. Infect. Dis.* **81**, 50–52 (2015).

125. Olumuyiwa Olaitan, A., Diene, S. M., Victor Assous, M. & Rolain, J. M. Genomic plasticity of multidrug-resistant NDM-1 positive clinical isolate of providencia rettgeri. *Genome Biol. Evol.* **8**, 723–728 (2016).

126. Pseudomonas aeruginosa Infection | HAI | CDC. https://www.cdc.gov/hai/organisms/pseudomonas.html.

127. Botelho, J., Grosso, F. & Peixe, L. Antibiotic resistance in Pseudomonas aeruginosa – Mechanisms, epidemiology and evolution. *Drug Resist. Updat.* **44**, 100640 (2019).

128. Zeng, L. *et al.* Genetic characterization of a blaVIM-24-Carrying IncP-7ß plasmid p1160-VIM and a blaVIM-4-harboring integrative and conjugative element Tn6413 from clinical pseudomonas aeruginosa. *Front. Microbiol.* **10**, 1–9 (2019).

129. Rickettsial Diseases (Including Spotted Fever & Typhus Fever Rickettsioses, Scrub Typhus, Anaplasmosis, and Ehrlichioses) - Chapter 4 - 2020 Yellow Book | Travelers' Health | CDC. https://wwwnc.cdc.gov/travel/yellowbook/2020/travel-related-infectious-diseases/rickettsial-including-spotted-fever-and-typhus-fever-rickettsioses-scrub-typhus-anaplasmosis-and-ehr.

130. Merhej, V. & Raoult, D. Rickettsial evolution in the light of comparative genomics. *Biol. Rev.* **86**, 379–405 (2011).

131. Questions and Answers | Typhoid Fever | CDC. https://www.cdc.gov/typhoid-fever/sources.html.

137. Questions & Answers | Shigella – Shigellosis | CDC. https://www.cdc.gov/shigella/general-information.html.

138. J, I., D, S., A, S., PD, C. & A, D. Characterization of antimicrobial resistance, plasmids, and gene cassettes in Shigella spp. from patients in

vietnam. *Microb. Drug Resist.* **9 Suppl 1**, (2003).

139. Staphylococcus aureus in Healthcare Settings | HAI | CDC. https://www.cdc.gov/hai/organisms/staph.html.

146. Clinical Information about Group B Strep | CDC. https://www.cdc.gov/groupbstrep/clinicians/index.html.

147. Pinkbook: Pneumococcal Disease | CDC. https://www.cdc.gov/vaccines/pubs/pinkbook/pneumo.html.

148. Lehtinen, S. *et al.* Horizontal gene transfer rate is not the primary determinant of observed antibiotic resistance frequencies in streptococcus pneumonia. *Sci. Adv.* **6**, 1–9 (2020).

149. Pharyngitis (Strep Throat): Information For Clinicians | CDC. https://www.cdc.gov/groupastrep/diseases-hcp/strep-throat.html.

150. Del Grosso, M. *et al.* ICESpy009, a conjugative genetic element carrying mef(E) in Streptococcus pyogenes. *Antimicrob. Agents Chemother.* **60**, 3906–3912 (2016).

151. Haslam, D. B. & St. Geme, J. W. Viridans Streptococci, Abiotrophia and Granulicatella Species, and Streptococcus bovis. in *Principles and Practice of Pediatric Infectious Disease* (eds. Long, S. S., Pickering, L. K. & Prober, C. G.) 719–723 (Elsevier Saunders, 2008). doi:10.1016/b978-0-7020-3468-8.50127-9.

152. Doern, C. D. & Burnham, C. A. D. It's not easy being green: The viridans group streptococci, with a focus on pediatric clinical manifestations. *J. Clin. Microbiol.* **48**, 3829–3835 (2010).

153. Balsalobre, L., Ferrándiz, M. J., Liñares, J., Tubau, F. & De la Campa, A. G. Viridans group streptococci are donors in horizontal transfer of topoisomerase IV genes to Streptococcus pneumoniae. *Antimicrob. Agents Chemother.* **47**, 2072–2081 (2003).

154. STD Facts - Syphilis (Detailed). https://www.cdc.gov/std/syphilis/stdfact-syphilis-detailed.htm.

155. Yaws. https://www.who.int/news-room/fact-sheets/detail/yaws.

156. Pathogen Safety Data Sheets: Infectious Substances – Ureaplasma urealyticum - Canada.ca. https://www.canada.ca/en/public-health/services/laboratory-biosafety-biosecurity/pathogen-safety-data-sheets-risk-assessment/ureaplasma-urealyticum.html.

157. Waites, K. B., Katz, B. & Schelonka, R. L. Mycoplasmas and ureaplasmas as neonatal pathogens. *Clin. Microbiol. Rev.* **18**, 757–789 (2005).

158. General Information | Cholera | CDC. https://www.cdc.gov/cholera/general/index.html.

159. Verma, J. *et al.* Genomic plasticity associated with antimicrobial resistance in Vibrio cholerae. *Proc. Natl. Acad. Sci. U. S. A.* **116**, 6226–6231 (2019).

160. Nelson, J. D. & McCracken, G. H. The pediatric infectious disease journal(r) newsletter: march 2009. *Pediatr. Infect. Dis. J.* **28**, A5 (2009).

161. Ecology and Transmission | Plague | CDC. https://www.cdc.gov/plague/transmission/index.html.
